# Behavior evolves as a correlated response to selection on cuticle color in *Drosophila melanogaster* and *D. simulans*

**DOI:** 10.64898/2026.01.13.699118

**Authors:** Sarah N. Ruckman, Alexandra G. Duffy, Paulina Montes Mendez, Katelyn D. McCaffery, Samuel Miller, Addison Crews, Nicholas Tan, Lauren A. Campbell, Ashley March, Elizabeth B. Brown, David Houle

## Abstract

To predict adaptive evolution, we need to understand the degree to which selection on one trait can constrain or redirect evolutionary responses in other traits. We used artificial selection on cuticle color in *Drosophila melanogaster* and *D. simulans* to investigate whether color and behavior evolve in tandem. We selected for light and dark thoracic colors for 16 generations in two populations per species and measured correlated responses in aggression, basal activity, total activity, sleep, and geotaxis. Dark selected individuals consistently showed higher aggression and basal activity than light selected or control flies across both species and sexes, pointing to a modest but repeatable correlated response. In contrast, patterns for geotaxis, sleep, and total activity evolved unpredictably, often varying across species and populations, and showed no clear association with color. Within-population phenotypic correlations between color and aggression were significant but weak (rho = -0.035, rho² = 0.001), suggesting that color is a poor predictor of individual aggressive behavior. Populations evolving higher aggression also tended to evolve higher basal activity (rho = 0.52, p = 0.004), suggesting activity may have evolved as a correlated response rather than through a direct color-activity link. The persistence of correlated responses in aggression and basal activity in response to selection on color is consistent with a conserved genetic basis, such as pleiotropy or linkage disequilibrium, although our data do not distinguish between these mechanisms. Identifying the loci underlying variation in both color and behavior, and testing their effects experimentally, will clarify how multivariate genetic covariance shapes the direction and pace of adaptive evolution.

## Introduction

A key goal of modern evolutionary biology is to predict if and how populations will evolve adaptively in response to environmental change. Predicting adaptive evolution requires identifying constraints that might limit a population’s ability to respond to selection. Definitions of adaptive constraints vary, but they all emphasize the same core concept: that populations are not always able to respond to selection as one might predict from a simple model of adaptation (Blows and Walsh 2009; Walsh and Blows 2009; Connallon and Hall 2018; Hansen and Pélabon 2021). One important source of long-term adaptive constraint is genetic covariance, which can prevent traits from evolving independently or potentially slow adaptation (Houle 1991; Connallon and Hall 2018; Hansen and Pélabon 2021).

Numerous empirical studies in both natural and experimental populations have tested whether these genetic covariances constrain adaptation. In natural populations, selection on one trait can cause changes in another trait, such as the linked evolution of beak morphology and song in Darwin’s finches (Podos 2001) or the covariation of body shape and armor in threespine sticklebacks (Leinonen et al. 2011). Experimental evolution and artificial selection studies in model organisms allow researchers to identify the nature of underlying genomic differences associated with genetic correlations. For example, studies of locomotor behavior and body size in nematodes show that multivariate phenotypic evolution is predictable from ancestral genetic variation (Mallard et al. 2023). Gene expression evolution driven by indirect selection on genetically correlated genes during adaptation to temperature and drought stress in *Tribolium* demonstrates that genetic correlations shape the direction of adaptation (Koch et al. 2025). In *Drosophila melanogaster*, laboratory selection for decreased development time led to correlated changes in body size and lifespan, along with divergence across dozens of loci (Burke et al. 2010). Experimental evolution across thermal environments showed that wing morphology can evolve along genetic lines of least resistance, though this depends on environmental heterogeneity (Walter 2023). Collectively, these studies demonstrate that correlated trait responses are likely common and, thus, that naïve expectations about adaptation may be incorrect. Experimental studies are essential if we are to fully understand how traits constrain adaptive evolution.

Color and behavior are two ecologically important traits that are often key targets of selection and have been previously demonstrated to be correlated at the phenotypic level (Ducrest et al. 2008; Roulin and Ducrest 2011; San-Jose and Roulin 2018; Ruckman et al. 2024). One explanation for these correlations is that the biochemical pathways underlying coloration (e.g., melanin synthesis) influence neurophysiology and behavior (Ducrest et al. 2008; San-Jose and Roulin 2018). In invertebrates, for example, melanin coloration arises from dopamine metabolism, and changes in gene expression within the pathway are linked to variation in color intensity (Wittkopp and Beldade 2009; Takahashi 2013). Consequently, mutations in key pigmentation genes, such as *ebony* and *yellow,* not only alter color but also cause predictable behavioral defects, including abnormalities in circadian rhythms and mating behavior (Suh and Jackson 2007; reviewed in Takahashi 2013). However, a recent meta-analysis across Animalia (Ruckman et al. 2024) revealed a consistent positive correlation between color and behavior, which was not restricted to only melanin-based colors, challenging the exclusivity of the melanin-pleiotropy hypothesis. Instead, the results are more consistent with the ’badge of status’ hypothesis as the most parsimonious explanation, where conspicuous colors act as honest signals of an individual’s aggressive potential (Rohwer 1975; McGraw et al. 2003; Tibbetts and Dale 2004). Under this hypothesis, correlational selection jointly favors individuals possessing both dark coloration and high aggression, which can generate and maintain a genetic correlation between the traits through linkage disequilibrium or by acting on existing pleiotropic variation. Consistent with this interpretation, Ruckman et al. (2024) found no evidence for condition dependence, suggesting that the correlation between color and aggression is more likely to reflect underlying genetic covariance than condition-based signaling. The conclusion from this meta-analysis, however, was inferred from broad-scale correlational patterns, leaving the underlying genetic mechanism, pleiotropy or linkage disequilibrium, unresolved.

Distinguishing between pleiotropy and linkage disequilibrium can be challenging when testing for correlated responses to selection in a single species. Experimental evolution studies comparing responses among two or more related species subjected to the same selection regime can be a powerful tool because linkage disequilibrium is generally not expected to be conserved over longer evolutionary timescales (Falconer and Mackay 1996; Gaut and Long 2003; Slatkin 2008). A consistent pattern of correlated trait evolution observed repeatedly in different taxa would provide strong evidence for pleiotropy or widespread correlational selection jointly favoring both traits (Lande 1979; Cheverud 1988, 1996; Wagner and Zhang 2011). Moreover, even in the absence of recombination, simulations show that genetic correlations maintained by linkage are weaker than those maintained by pleiotropy, because mutation at one of the two linked loci can alter one trait without affecting the other, breaking the correlation (Chebib and Guillaume 2021). A third mechanism that can generate and maintain genetic correlations is correlational selection, in which two traits are jointly favored by selection regardless of their shared genetic architecture (Lande 1980; Chebib and Guillaume 2021).

Here, we focused on two species of fruit flies, *D. melanogaster* and *D. simulans*. These recently diverged species (estimates vary from 2-8 million years ago; Capy and Gibert 2004; Obbard et al. 2012) are important models for experimental evolution (e.g., Burke et al. 2010; Turner et al. 2011, 2013; Remolina et al. 2012; Barghi et al. 2017; Kelly and Hughes 2019; Bastide et al. 2022; Girardeau et al. 2025) due to their tractability for large-scale experiments (Hoffmann 1987; Kravitz and Huber 2003; Kravitz and Fernandez 2015; Massey and Wittkopp 2016). Furthermore, in *Drosophila*, both cuticle pigmentation (Wittkopp et al. 2011; Rajpurohit et al. 2016; Bastide et al. 2022) and behavior (Sokolowski 2001; Martin 2003; Anholt and Mackay 2015; Kravitz and Fernandez 2015) are ecologically relevant traits that influence fitness.

We hypothesized that selection for darker color would drive a correlated suite of behavioral changes (aggression, basal activity, geotaxis, total activity, and sleep) in comparison to light selected individuals for both *D. melanogaster* and *D. simulans*. We predicted dark selected individuals would be more aggressive and have increased locomotor activity (Jordan et al. 2006, 2007; Ducrest et al. 2008; Wittkopp and Beldade 2009; Takahashi 2013). Given previous evidence of a negative correlation between locomotor activity and sleep in *D. melanogaster* (Keene et al. 2010; Harbison et al. 2013), we predicted dark selected flies (i.e., more active flies) would sleep less. Finally, because activity level and geotaxis have been shown to be genetically correlated in *D. melanogaster* (Murphey and Hall 1969; Toma et al. 2002; Zhong et al. 2022), we predicted that dark selected flies would have a faster geotactic response. The evolution of one or more of these behaviors in tandem with color would support the genetic covariance hypothesis, providing direct experimental evidence that genetic covariance influences the evolution of this suite of traits. A consistent correlated response in the opposite direction would also support a shared genetic basis, but would be inconsistent with the melanin-pleiotropy literature. A correlated response in only one species or one sex would suggest the genetic basis is not fully conserved. No consistent correlated response in either species would suggest that color and behavior do not share a conserved genetic basis. If selection on color produces correlated behavioral changes in both species and sexes, this would suggest a conserved genetic basis, such as pleiotropy, linkage disequilibrium, or correlational selection in both species. Critically, the badge of status hypothesis requires that color reliably predicts aggressive potential at the individual level within populations, enabling conspecifics to assess an opponent’s fighting ability from its coloration. This predicts a strong negative phenotypic correlation between color (darker individuals have lower grey scale values) and aggression within populations. Pleiotropy and linkage disequilibrium, in contrast, can produce correlated evolutionary responses to selection without necessarily generating a strong within-population phenotypic correlation, particularly when the heritability of the behavioral trait is low.

## Methods

### Fly maintenance

We housed each species in a separate incubator (Percival Model # 136VLC8Q9) on a 12-hour light and 12-hour dark cycle. We maintained *D. melanogaster* populations (*D. mel*-1 and *D. mel*-2) at a constant temperature of 25 °C, while *D. simulans* (*D. sim*-1 and *D. sim*-2) were housed at a constant temperature of 24 °C (Krstevska and Hoffmann 1994; Ashburner and Roote 2007; Austin and Moehring 2013). We housed each treatment in a separate plexiglass population cage (26.67 cm x 15.25 cm x 20.32 cm) and randomly assorted the cages into the incubator every week. Bottles containing a standard Drosophila media composition (Ashburner and Roote 2007) were added to each cage every week and were removed after three weeks.

### Artificial Selection

To limit the effects of bottlenecks early in the experiment, we used a gradual selection regime (Kofler and Schlötterer 2014; Vlachos and Kofler 2019). Each generation, we collected 750 virgin females and 750 males for each population (Supplemental Material for origin). We randomly assigned them to three equal-sexed groups to serve as the parents for the three treatments (Figure 1). In each population, we established two selection treatments (dark “D” and light “L” selected) and one control “C” treatment. We performed selection on the cuticle coloration of the dorsal thorax. This region of the cuticle is not sexually dimorphic (Wittkopp et al. 2003; Massey and Wittkopp 2016). To perform selection, flies were anesthetized with CO_2_. We scored the color of the dorsal thorax by eye into four color categories, using the prominent macrochaeta as landmarks (David et al. 1985; see Supplemental Material for details). Color selection was conducted by a single individual and validated (n=100) for agreement with objective color quantification measurements and repeatability (see Supplemental Material). Visual bins differed from one another in the expected directions (*F*_3, 96_= 138.19, *P*<0.001; Figure S1) and were highly repeatable (ICC = 0.933; F_99,99.1_ = 29.2, p < 0.001; Figure S2).

**Figure 1:**
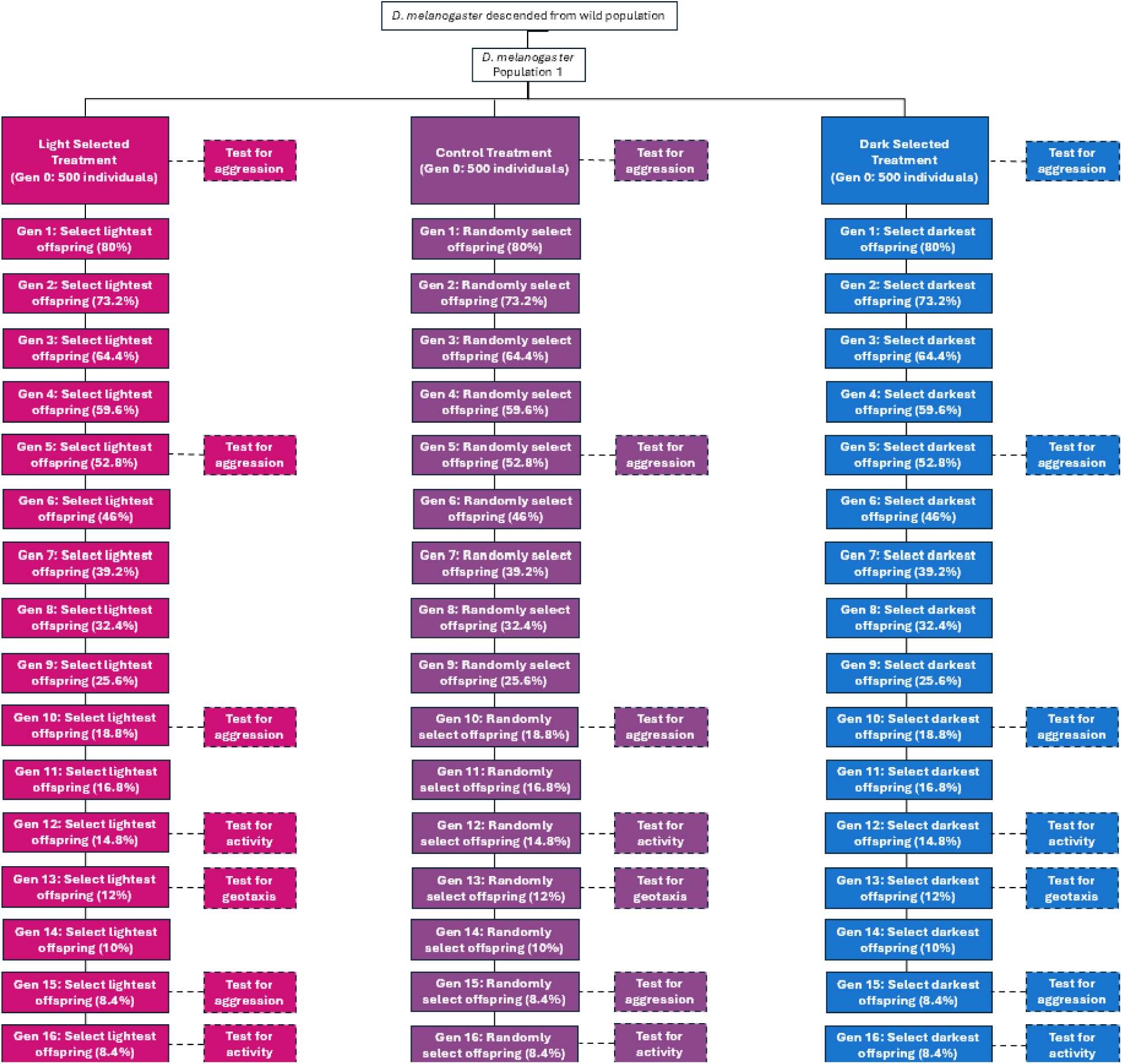
Selection and Behavioral Workflow. This workflow is designed for a single population of *D. melanogaster*. This same workflow was used for each population per species. Flies were collected from wild populations to create each population. They were then tested for aggression and split into three treatments (light selected “L”, control “C”, and dark selected “D”). The L treatment is represented by pink. The C treatment is represented by purple. The D treatment is represented by blue. Solid lines indicate the selection and establishment of a new generation. Dashed lines and dashed boxes indicate behavioral assays. Selection gradually increased, so fewer individuals were kept every generation (decreasing percentage of individuals kept every generation). The L treatment had the lightest offspring selected, while the D treatment had the darkest offspring selected each generation. The C treatment was randomly chosen each generation.

For D and L selection treatments, we gradually increased the coefficient of selection from 80% of 250 virgin females and 80% of 250 males to 8.4% of 250 virgin females and 8.4% of 250 males, over 16 generations. For example, for L, we selected the lightest 80% of 250 virgin females and 80% of 250 males in the first generation of selection. By the 16th generation, we selected only the lightest 8.4% of 250 virgin females and 8.4% of 250 males. C treatments flies were randomly chosen with respect to thorax color in each generation, while still mimicking the changes in population size experienced by the D and L treatments. Figure 1 illustrates this selection scheme in detail and indicates the generations in which phenotypes other than thorax color were assayed. No individuals used in the selection experiment were used in behavioral tests.

### Behavioral Assays

All behavioral tests were conducted on individuals 3-5 days old that were not used to establish the next generation of flies. We staggered behavioral assays across generations to ensure enough flies were available for selection experiments. For all behavioral assays, flies were isolated for 24 hours before testing. All tests were scored by observers who were blind to the hypothesis. However, the observer was not blind to the color or sex of the individual as these features were readily apparent.

#### Aggressive Behavior

Due to its previously reported association with color (Ducrest et al. 2008; San-Jose and Roulin 2018; Ruckman et al. 2024), we were particularly interested in aggression, and therefore, measured aggression at multiple generations (generations 5, 10, and 15; Figure 1) using a standard protocol (Girardeau et al. 2025). To assay aggression, we paired a focal individual with a marked opponent of the same sex, similar age, color, and size. Controlling for sex, age, and size accounts for known behavioral covariates in *Drosophila* (Hoffmann 1991; Chen et al. 2002; Baxter and Dukas 2017; Bath et al. 2021; Fernandez et al. 2023) Furthermore, we paired individuals of similar color because interactions can be modulated by an opponent’s coloration, a principle established in other species (e.g., Lank et al. 1995; Sinervo and Lively 1996; Kraft et al. 2018; Sacchi et al. 2021; Scali et al. 2021).

The focal individual was assigned randomly. The opponent’s left wing was marked to distinguish it from the focal individual. Females were first aspirated into new vials without media or water for two hours before conducting the trial to increase the likelihood of aggressive behaviors (Ueda and Kidokoro 2002). After two hours, both females were aspirated into the arena. For the male assays, flies were aspirated together into the arena 30 minutes after lights on that day. Males were not moved to empty vials prior to the assay because they do not require food deprivation to display aggressive behaviors (Chen et al. 2002; Ueda and Kidokoro 2002).

We used a standard aggression arena with a central food resource (see Supplemental Material for details; Hutchins et al. 2024; Girardeau et al. 2025). Following a five-minute acclimation period, we recorded the occurrence of fencing, lunging, boxing, and tussling (Chen et al. 2002; see Supplemental Material for behavioral descriptions) every five minutes for one hour (Altmann 1974).

#### Basal Activity Level

To measure basal activity level, we aspirated each individual into an open field arena 3 hours after lights on that day (see Supplemental Material for details). After a 10-minute acclimation period, the focal individuals were video-recorded for one minute, and we counted the number of grid lines crossed as a measure of activity (Burnet et al. 1988; Anderson et al. 2016).

#### Geotaxis Level

*Drosophila* exhibit an innate negative geotaxis response (Hirsch 1959; Toma et al. 2002). To measure geotaxis, we aspirated each individual into the geotaxis arena (Taylor and Tuxworth 2019; Zhong et al. 2022; see Supplemental Material for details). After tapping a fly to the bottom, we timed how long it took to climb a 2 cm distance. Trials were censored and assigned a score of 0 after one minute if the individual did not climb the 2 cm distance.

#### Total Activity Level and Sleep

We measured activity and sleep over 72 hours using the *Drosophila* Activity Monitoring (DAM) system (Trikinetics, Waltham, MA). Flies were individually housed in tubes with standard media and acclimated for 24 hours before data collection. Total activity is measured as total beam crossings per day. Total sleep is defined as the number of 5-minute bouts of inactivity per day (Pfeiffenberger et al. 2010a,b; see Supplemental Material for details).

### Color Quantification of Individuals Used in Behavioral Assays

Following behavioral assays, except for total activity and sleep, all individuals were photographed using a standardized apparatus (WINGMACHINE; Houle et al. 2003). The mean grayscale value of the dorsal thorax was quantified from these images using ImageJ after calibration to a color standard (Schneider et al. 2012; see Supplemental Material for the imaging and analysis protocol).

### Statistical Analyses

All statistical analyses were conducted using R version 4.5.0 (R Core Team 2025). All data and R code are available on GitHub (https://github.com/sruckman/Exp_Evo_2024). Raw data distributions are provided in Figures S4-S11. To test for direct and correlated responses to selection, we used a consistent generalized linear model (GLM) framework for each trait. All models were analyzed separately for males and females because the residual distributions differed between the sexes. For each trait, we visually inspected residuals and fitted values to verify that model assumptions (e.g., normality, homoscedasticity, goodness of fit) were met.

For the color and behavioral analyses, the full model for each trait related the dependent variable (e.g., grayscale value, aggression counts) to the fixed effects of species, treatment, and population (nested within species), including all two-way interactions. To test our prediction that selection treatments diverged in the predicted directions, we coded treatment as a continuous predictor with values -1, 0, and 1 corresponding to L, C, and D lines, respectively. This parameterization allowed us to estimate both the direction and magnitude of the evolutionary response to selection. Models were fit by maximum likelihood, and we used F-tests for Gaussian models and Wald χ² statistics for GLMs to assess the significance of main effects and interactions. Non-significant interaction terms (p > 0.05) were removed before testing main effects.

We selected error distributions appropriate to each trait: a normal distribution for thorax grayscale values, total activity, and sleep; a Poisson distribution with a log link for counts of aggressive acts; and a negative binomial distribution for basal activity counts. For geotaxis, we analyzed time-to-event data using a Cox proportional hazards mixed-effects model (coxme package; Therneau 2024) with observer as a random effect and right-censoring of individuals that did not complete the task within one minute.

When we detected a significant main effect or interaction involving treatment, we re-coded treatment as a categorical factor, and conducted post-hoc comparisons using the emmeans package (Lenth 2025). For Gaussian and negative binomial models, we report least-squares means and pairwise t-tests; for Poisson models, we used asymptotic z-tests with effectively infinite degrees of freedom. When treatment-by-population interactions were significant, we further characterized among-population differences in response to selection using simple slopes analyses implemented with the emtrends function in emmeans, which estimates the treatment slope within each population and tests it using Wald X^2^ statistics.

To examine the temporal dynamics of the response to selection and differences in evolutionary trajectories among treatments, we fit models similar to those above, but also including generation and generation^2^ as predictors, along with their interactions with treatment. For thorax color, we used a Gaussian model, and for aggression, we used a Poisson GLM with a log link. Full model specifications, diagnostics, and trait-specific details (color, aggression, basal activity, total activity, sleep, and geotaxis) are provided in the Supplementary Methods.

To test the badge of status prediction that color reliably predicts aggressive potential at the individual level, we calculated Spearman rank correlations between individual color measurements and behavioral scores within each combination of line, species, sex, and generation. This within-stratum approach avoids inflation of correlations by known species, line, sex, and generation effects. We then averaged within-stratum correlations using Fisher’s z-transformation, weighting each stratum by its sample size (n-3), and back-transformed the data to obtain a weighted average correlation with 95% confidence intervals. To test whether correlations changed over time or differed between lines, species, or sexes, we regressed Fisher z-transformed correlations on generation using weighted least squares and tested for interactions between generation and line, species, and sex. To test for significant differences between specific time points, we used pairwise z-tests comparing Fisher z-averaged correlations, with test statistics computed as the difference in z-averaged correlations divided by the square root of the sum of squared standard errors, and p-values obtained from the standard normal distribution. To examine whether the correlated response in basal activity was associated with the correlated response in aggression, we computed a Spearman rank correlation between group mean aggression and group mean basal activity. Since these traits were measured in separate assays on different individuals, correlations were computed at the group level, with each group defined as a unique combination of line, population, and sex (3 lines × 4 populations × 2 sexes = 24 groups). Mean aggression was averaged across generations 5, 10, and 15 to reduce noise from single-generation measurements. We used Spearman correlations because the behavioral measures violated the assumptions of normality. All correlation analyses were conducted in R using the cor.test and lm functions.

## Results

### Evolution of Color in Selection Treatments

To examine the temporal dynamics of the direct response to selection, we analyzed the divergence in thorax color between D and L selected lines at generations 5, 10, and 15. Since the full statistical model revealed complex interactions (Table S1), we visualized the response by plotting the mean difference (contrast) in color between D and L treatments for each population over time (Figure 2). In both sexes, the divergence between D and L lines was significant by generation 5 and increased in a roughly linear fashion throughout the experiment (Table S2; Figure 2). In males, the pattern was broadly consistent, though the *D. sim*-2 population tended to exhibit less divergence than the other three populations in most generations (Figure 2A). The response in females was remarkably consistent across all four populations (Figure 2B).

**Figure 2:**
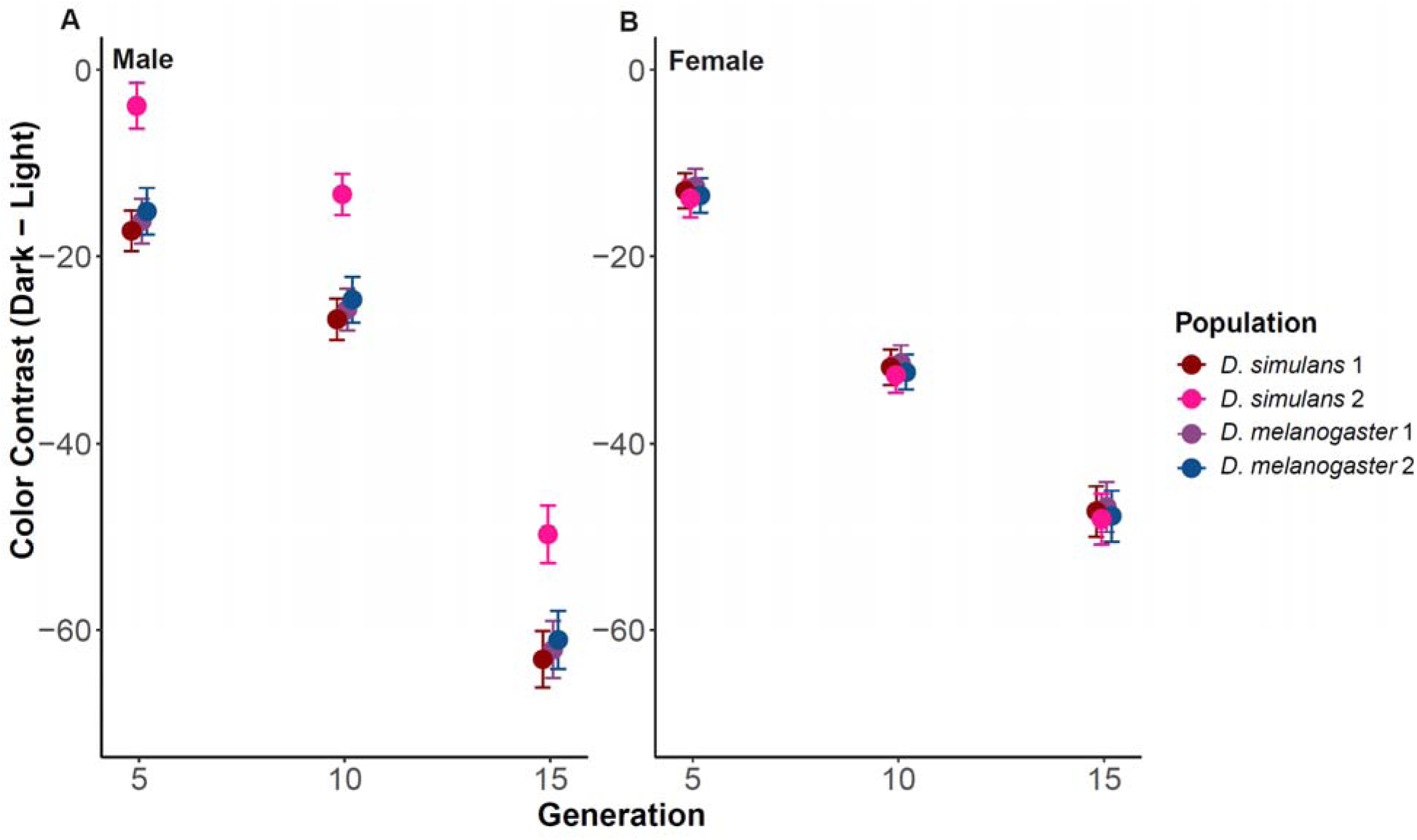
Difference (contrast) in mean thorax color between D and L flies within a population at a specific generation of selection. The colors represent the different populations. The x-axis represents the generation number in which traits were assayed, and the y-axis represents the D - L color contrast (mean ± SE). The mean is the difference between the raw data mean of the D and L treatments. The error bars represent the standard errors of raw data. However, as described in the Methods, the statistical analysis was conducted using the individual fly color measure as the dependent variable. A negative contrast indicates that D flies were darker than L flies. The left panel shows contrast values for males of both populations of both species, while the right panel shows the contrasts for females.

We next examined the magnitude and among-population structure of color differences at the end of the experiment (Table S3; Figure 3). There was a significant effect of treatment by population nested in species interaction (males: F_2,345_ = 12.43, p < 0.0001; females: F_2,337_ = 5.20, p = 0.006), indicating that the magnitude of the response to selection varied among populations within a species. However, our analysis of rates of response was in the same direction and was significantly negative in all cases (all slopes < -19.4, all X^2^_1_ > 90.5, all p < 0.0001; Table S4). Post-hoc tests further confirmed that for all populations and sexes, D treatment flies were significantly darker than L and C treatment flies (Table S5; Figure 3), and L treatment flies were significantly lighter than C treatment flies (Table S5; Figure 3).

**Figure 3:**
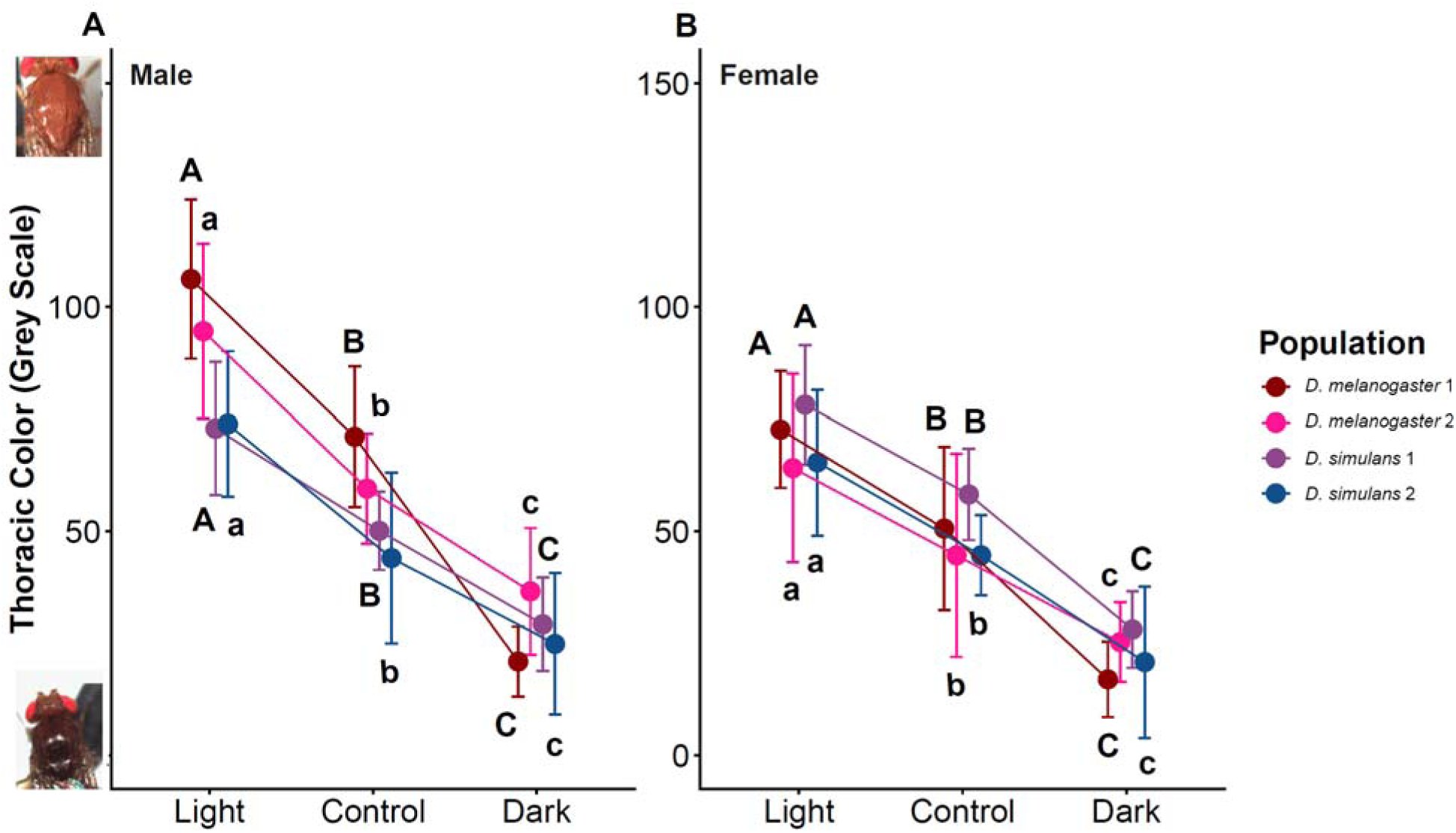
Thoracic color evolution at generation 15 for A) Male analysis B) Female analysis. The x-axis indicates the different treatments (L, C, and D). The y-axis represents the average grey-scale value, with lighter values (higher on the scale) at the top and darker values (lower on the scale) at the bottom. Photos on the y-axis represent individuals before grey-scale conversion in ImageJ. The letters above and below the data indicate significance, where groups with different letters are significantly different from each other. Uppercase letters indicate the significance of *D. mel*-1 and *D. sim*-1, and lowercase letters indicate the significance of *D. mel*-2 and *D. sim*-2 populations.

Despite the consistent direction of the response, there were notable differences among groups that explain the significant interaction. For instance, *D. melanogaster* males were, on average, lighter than *D. simulans* males, a difference driven primarily by the L and C treatments (Table S4; Figure 3A). The response in *D. mel-*1 males was particularly strong, with a selection slope twice as steep as any other population (−46.2 vs. -21.9; Table S4). Among females, there was a significant difference between the replicate *D. simulans* populations, with *D. sim*-1 females tending to be darker than *D. sim*-2 females across all treatments (Tables S5 and S6; Figure 3B).

### Correlated Evolution of Aggression in Selection Treatments

To test our *a priori* prediction about the correlated evolution of cuticle color and aggression, we assayed aggression across multiple generations during the selection experiment. To simplify interpretation (Table S7), we visualized the temporal response by plotting the divergence in the number of behaviors (contrast) between the D and L treatments over time (Figure 4). In males, this revealed a non-linear response that differed between the species (Figure 4A). The divergence in both species decreased slightly at generation 10 and was greatest at generation 15 (Figure 4). Despite divergence in color at generation 5, *D. simulans* populations did not diverge in aggressive behavior until generation 15 (Table S8; Figure 4A). However, *D. melanogaster* populations diverged in both color and behavior by generation 5 and consistently showed greater divergence than *D. simulans* populations (Table S8; Figure 4A). Females exhibited a similar non-linear trajectory, with the divergence between D and L lines also peaking at generation 15 after a minimum at generation 10 (Figure 4B). For females, however, the variation was most pronounced between populations rather than species (Figure 4B). The *D. mel*-1 population, in particular, exhibited a greater divergence between D and L treatments at each generation than the other three populations (Table S8; Figure 4B).

**Figure 4:**
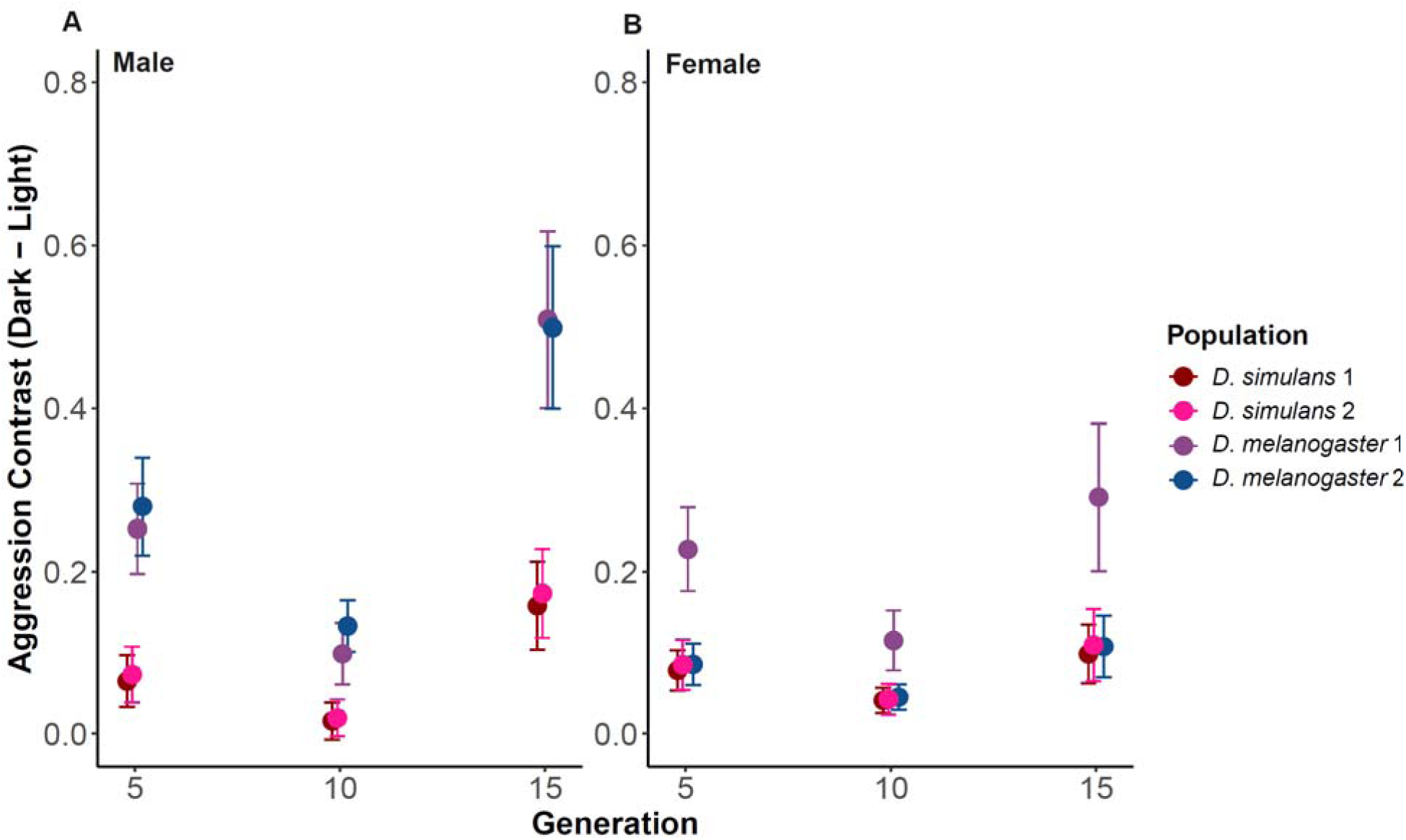
Difference (contrast) in mean aggression counts between D and L flies within a population at a specific generation of selection. The colors represent the different populations. The x-axis represents the generation number in which traits were assayed, and the y-axis represents the D - L color contrast (mean ± SE). The mean is the difference between the raw data mean of the D and L treatments. The error bars represent the standard errors of the raw data. However, as described in the Methods, the statistical analysis was conducted using the individual fly aggression counts as the dependent variable. A positive contrast indicates that D treatment flies were more aggressive than L flies. The left panel shows contrast values for males of both populations of both species, while the right panel shows the contrasts for females.

Given these nonlinear divergence trajectories and their peak at generation 15, we next examined the magnitude and among-population structure of aggression differences at the end of the experiment (Table S9; Figure 5). There was a significant main effect of selection treatment on aggression for both sexes (males: X^2^_1_ = 50.3, p < 0.0001; females: X^2^_1_ = 17.5, p < 0.0001; Table S9). This response was consistent across all populations, as the treatment by population interaction was not significant in initial models (males: X^2^_2_ = 0.23, p = 0.89; females: X^2^_2_ = 0.13, p = 0.94) and was therefore dropped from the final model. As predicted, we observed a significant positive slope for both males (slope = 1.52, X^2^_1_ = 31.5, p < 0.0001; Table S10) and females (slope = 1.1, X^2^_1_ = 13.3, p = 0.0003; Table S10). Post-hoc tests revealed that D individuals were significantly more aggressive than L and C treatment individuals, which did not differ from each other (Table S11; Figure 5).

**Figure 5:**
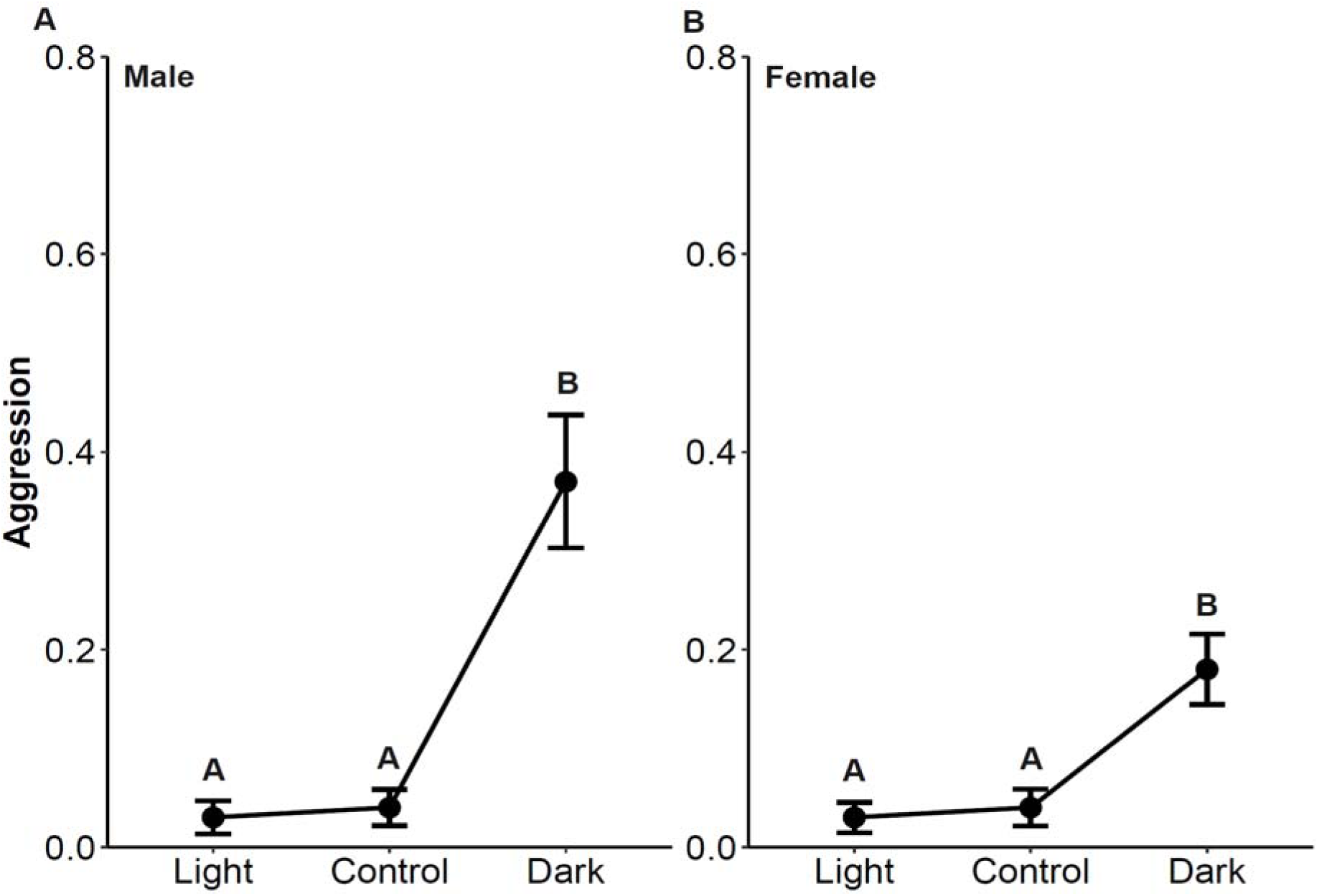
Aggression at generation 15 for A) Male analysis. B) Female analysis. The x-axis indicates the different treatments (L, C, and D). The y-axis represents the average aggression counts (number of aggressive acts). The letters above the data indicate significance, where groups with different letters are significantly different from each other.

### Correlated Evolution of Basal Activity Level

At generation 12, basal activity showed a significant correlated response to selection (Table S12; Figure 6). This response was consistent across all populations, as the treatment by population interaction in initial models was not significant (males: X^2^_2_ = 1.35, p =0.51; females: X^2^_2_ = 0.10, p =0.95) and was therefore dropped from the final models. As predicted, we observed a significant positive slope for both males (slope = 0.51, X^2^_1_ = 34.0, p < 0.0001; Table S13) and females (slope = 0.27; X^2^_1_ = 11.7, p = 0.0006; Table S13). Post-hoc tests showed that L individuals did not differ from C individuals (Table S14). In males, D individuals were significantly more active than both L and C individuals, which did not differ from each other (Figure 6A). In females, D individuals were significantly more active than L individuals, but not C individuals (Figure 6B).

**Figure 6:**
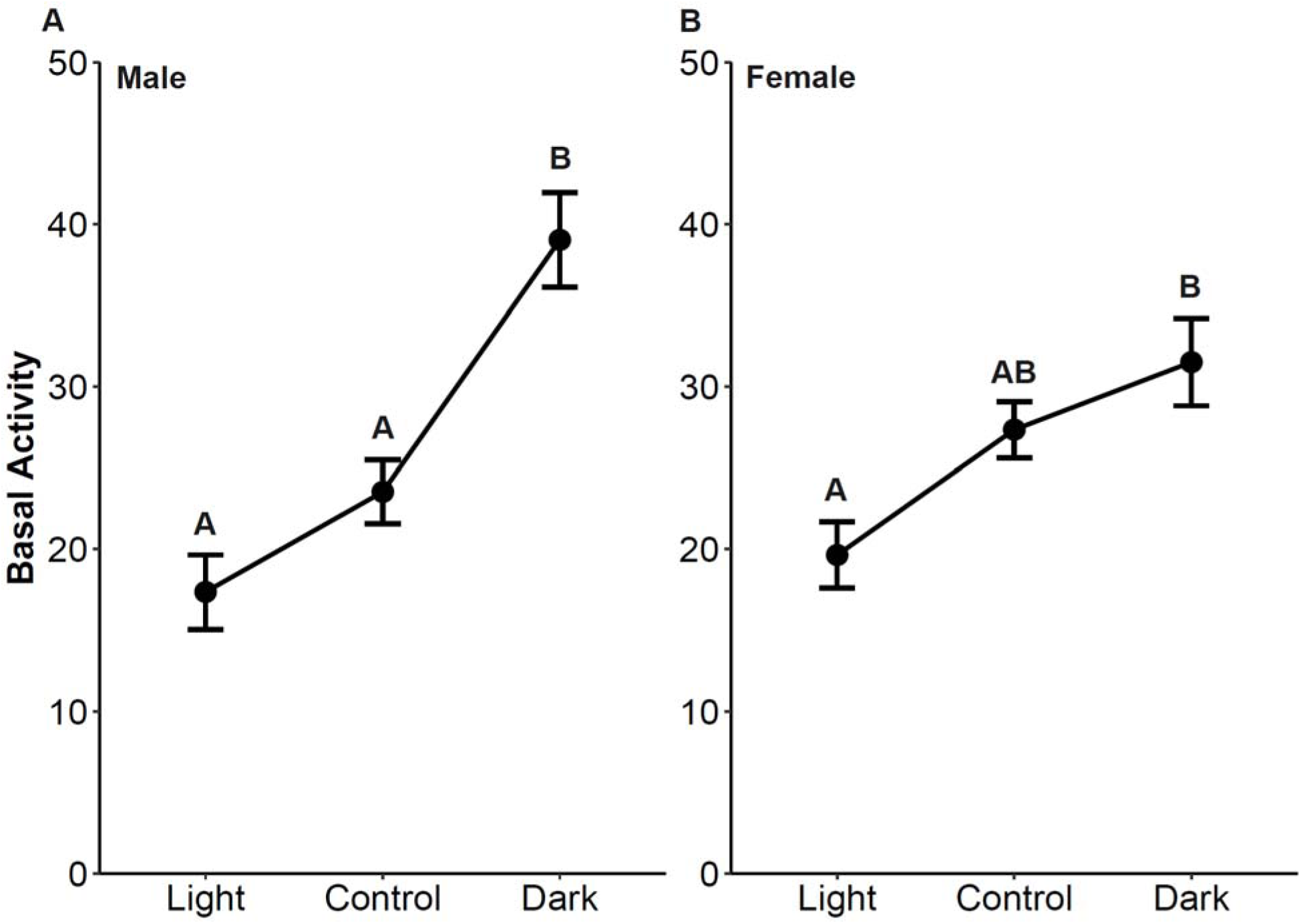
Basal activity level at generation 12 for A) Male analysis. B) Female analysis. The x-axis indicates the different treatments (L, C, and D). The y-axis represents the average basal activity level (number of lines crossed in one minute). The letters above the data indicate significance, where groups with different letters are significantly different from each other.

### Inconsistent Correlated Responses in Other Behaviors

In contrast to aggression and basal activity, the correlated responses for geotaxis, total activity, and sleep were inconsistent and unpredictable, with responses varying across species, populations, and sexes (see Supplemental Material; Tables S15-S23; Figures S12-S17). For example, D lines showed increased geotaxis in some populations but decreased geotaxis in others, with no consistent directional pattern across species or sexes. Similarly, total activity and sleep showed population-specific divergence with no consistent differences among color treatments.

### Phenotypic Correlations between Color and Behavior

To test whether color predicts aggressive potential at the individual level, we calculated within-population phenotypic correlations between color and aggression. Within-population correlations were significant but weak overall (rho = -0.035, rho² = 0.001, p = 0.003, n = 7,078; Table S24). The color-aggression correlation shifted significantly from near zero at generation 0 (rho = 0.011; Table S24) to negative by generation 15 (rho = -0.093; Table S24; pairwise Fisher z-test: p = 0.019; Table S25), indicating that darker individuals tended to be slightly more aggressive within populations by the end of the experiment. C lines showed no significant shift between generation 0 and generation 15 in any stratum (all p > 0.09; Table S25). Within-population correlations between color and basal activity were not significant (rho = -0.020, rho² < 0.001, p = 0.502; Table S24). Mean aggression and mean basal activity were positively correlated across line, population, and sex combinations (rho = 0.52, rho² = 0.27, p = 0.004, n = 24 groups; Table S26), suggesting that populations evolving higher aggression also tended to evolve higher basal activity. Full results, including species- and sex-specific breakdowns and geotaxis correlations, are reported in the Supplementary Material (Table S24-S26).

## Discussion

Our experiment demonstrates that artificial selection on cuticle color in *D. melanogaster* and *D. simulans* is accompanied by consistent correlated changes in some, but not all, behaviors. Selection for darker coloration led to a consistent increase in aggression at generation 15 and basal activity in both sexes of both species. In contrast, geotaxis, sleep, and total activity evolved unpredictably, with responses varying among species, populations, and sexes.

The consistent evolution of heightened aggression by generation 15 in D flies across both species suggests a genetic association between melanin-based color and aggressive behavior. However, the strength of this association is modest. Absolute aggression levels in our assay were low, and trajectories through time were noisy, which limits how strongly we can infer a deterministic link between color and aggression. Nonetheless, the fact that D lines consistently tended to be more aggressive than C and L lines in both species, despite low baseline aggression levels and demographic noise, is consistent with a conserved color-aggression association. Consistent with this, the within-population color-aggression correlation strengthened significantly in D lines from generation 0 to generation 15, while C lines showed no significant shift over the same period, arguing against a general temporal trend as the explanation. Within D lines, males were consistently more aggressive than females, suggesting that sex specific expression modulates the magnitude of the response while preserving the direction of the association between color and aggression. Similar context dependence in the evolution of male aggression has also been documented when aggression itself is the direct target of selection in our *D. mel*-2 base population (Girardeau et al. 2025). In that study, aggression was selected using three different social contexts. These contexts generated distinct evolutionary trajectories: the rate and pattern of change in total aggression differed markedly among treatments and were often noisy rather than smoothly directional. This suggests that treatment-specific responses are a general feature of this trait rather than unique to our selection experiment.

Our data revealing an association with cuticle color and aggression aligns with predictions from the melanin-pleiotropy hypothesis (Takahashi 2013; San-Jose and Roulin 2018), which posits a shared biochemical basis for the traits, and is also compatible with scenarios in which color could function as a status-related signal, with melanin-based coloration as the signal and aggressive propensity as the underlying quality (Rohwer 1975; Ruckman et al. 2024). However, given the low and variable aggression levels in our experiment, our data should be viewed as suggestive rather than definitive evidence for these mechanisms. At present, they indicate that darker cuticle coloration is associated with a greater propensity to engage in aggressive acts under our assay conditions, but they do not by themselves establish that melanin-based traits are universally reliable “badges of status” or that pleiotropy is the sole cause of the observed covariance. Within-population phenotypic correlations between color and aggression were statistically significant but weak (rho = -0.035, rho² = 0.001), indicating that color explained less than 0.1% of individual variation in aggression within populations. This weak phenotypic association is difficult to reconcile with the badge of status as the primary mechanism, which requires that color reliably predicts individual aggressive potential, allowing conspecifics to assess fighting ability from coloration. A weak phenotypic correlation does not, however, preclude a meaningful genetic association. Phenotypic correlations reflect both genetic and environmental sources of covariance, and when heritability of the behavioral trait is low, even a substantial genetic correlation will produce only a weak phenotypic signal (Falconer and Mackay 1996).

Correlational selection, in which dark color and aggression are jointly favored regardless of a shared genetic basis, could also maintain this association in natural populations (Lande 1980; Chebib and Guillaume 2021). However, our experimental design imposed selection exclusively on cuticle color, so correlational selection on aggression could not have operated. Any correlated response in aggression is most likely to reflect a genetic link between the traits, rather than incidental natural selection acting only on the D line. The consistency of the correlated response across *D. melanogaster* and *D. simulans* is more compatible with pleiotropy than linkage disequilibrium. These species have been evolving independently for millions of years, making conservation of specific chromosomal linkage associations unlikely (Falconer and Mackay 1996; Gaut and Long 2003; Slatkin 2008). Under linkage disequilibrium, the specific allelic associations underlying the color-aggression correlation would be expected to differ between lineages after millions of years of independent evolution, mutation, and recombination. Pleiotropy, in contrast, predicts a consistent directional association regardless of species-specific history. Our data, however, does not allow us to distinguish between pleiotropy and linkage disequilibrium, and a formal distinction between these mechanisms will require additional genetic approaches.

The correlated response in aggression was notably asymmetric in both species: D lines became more aggressive, whereas L lines did not become less aggressive than C lines. The lack of response in L lines is consistent with another experiment using the same assay in *D. mel*-2 (Girardeau et al. 2025). A likely explanation is that the base populations exhibited very low aggression under our assay conditions, such that further reductions in the underlying tendency to act aggressively would be difficult to resolve with our behavioral measure. In other words, we may be closer to a floor than a midpoint in aggression, which would naturally make upward responses easier to detect than downward ones. In natural populations, aggression carries real costs, including injury, energy expenditure, and predation risk, which balance its benefits and likely maintain aggression at an intermediate level (Georgiev et al. 2013). In the lab, both costs and benefits are reduced: there are no predators, food is provided ad libitum, and there is no territory to defend. The ecological context that motivates territorial aggression in wild populations is therefore absent (Kravitz and Fernandez 2015). Since our populations were recently derived from wild-caught individuals, this reduction in aggression is more likely to reflect a plastic response to lab conditions than an evolutionary adaptation. The assay itself may not capture the full range of aggressive propensity, and a higher-resolution assay or longer observation periods might reveal more of the variation that underlies the response we detected.

In contrast to the linear divergence in color, the correlated response in aggression over time was non-linear, peaking at generation 15 after a dip at generation 10. Environmental artifacts (e.g., humidity or temperature fluctuations) are one possible explanation, but all lines were maintained and assayed under tightly controlled conditions. Such inconsistent trajectories are often observed for correlated traits (Falconer and Mackay 1996), whose evolution is mediated by the interplay between indirect selection, genetic drift in finite populations (e.g., Yoo 1980; Weber 1996), and the potential for the underlying genetic covariance (the G-matrix) itself to change over time (Steppan et al. 2002; Arnold et al. 2008). One possible future direction is to continue selection for additional generations to reduce noise in the correlated responses. Even with this noisy background, the repeated tendency for D lines in both species to show higher aggression than light and control lines suggests a real association between pigmentation and aggression rather than a purely transient effect.

The consistent correlated evolutionary response in basal activity in D flies may reflect a shared genetic basis between color and activity, though the nature of this association is not straightforward. Our data are consistent with previous work that identified *Dopa decarboxylase (Ddc)*, a gene essential for both dopamine synthesis and melanin production, as a key locus associated with locomotor behavior in *D. melanogaster* (Jordan et al. 2006, 2007). The finding that many genes associated with locomotor behavior are also linked to aggression and neural development further suggests potential widespread genetic integration among these traits (Jordan et al. 2007). Our finding that D males were more active than D females in both species, and that D lines consistently showed elevated basal activity, are consistent with genetic connections between pigmentation and locomotor behavior being expressed in a sex specific but directionally conserved way across taxa. Despite this clear correlated evolutionary response, within-population phenotypic correlations between color and basal activity were not significant (rho = - 0.020, p = 0.502). However, mean aggression and mean basal activity were positively correlated across line, population, and sex combinations (rho = 0.52, rho² = 0.27, p = 0.004), suggesting that populations evolving higher aggression also tended to evolve higher basal activity. This raises the possibility that basal activity evolved as a correlated response to the color-aggression genetic association rather than through a direct genetic link between color and activity.

In contrast, we did not observe a consistent correlated response in total activity, which was measured using the DAM system. This result is consistent with the findings of Dierick and Greenspan (2006), who selected directly for high aggression and used an older, but similar, DAM monitoring system to measure general activity. They also found no correlated increase in total activity. This suggests that the genetic covariance between color and activity is likely context-dependent, manifesting strongly in exploratory or basal activity but not in overall activity levels measured in a confined tube environment.

The stochastic evolution of geotaxis, sleep, and total activity suggests that these traits exhibit little to no conserved genetic covariance with cuticle color, although we cannot rule out the possibility that, with continued selection across more generations, a modest association could emerge. While our gradual selection design was intended to allow recombination to break down transient, non-functional linkages (Kofler and Schlötterer 2014; Vlachos and Kofler 2019), the low effective population size, particularly in the final five generations, amplifies the impact of stochastic forces. These forces include genetic drift, but also stochasticity in the decay of any initial linkage disequilibrium. Together, these processes are expected to cause random, population-specific divergence in phenotypes not under strong, consistent indirect selection. Geotaxis, sleep, and total activity in our experiment showed exactly this pattern of idiosyncratic, population specific divergence with no consistent differences among color treatments. Their trajectories likely reflect drift or transient linkage rather than a stable conserved genetic association with pigmentation.

A productive direction for future research would be to determine the specific genetic mechanisms underlying these divergent outcomes. A primary goal is to determine if the consistent association between color, aggression, and basal activity reflects pleiotropy or linkage disequilibrium. Evolve-and-resequence (E&R) studies could identify candidate genes, and subsequent functional analyses, such as RNAi knockdowns, could then confirm their effects on both color and behavior, clarifying the extent to which such correlated responses constrain or facilitate evolution. A valuable intermediate step would be to estimate genetic correlations between color and behavior using a formal quantitative genetic breeding design. A significant genetic correlation consistent across species would strengthen the case for a shared genetic basis and help prioritize which trait combinations to target in functional studies. Beyond pairwise relationships, it will be important to characterize the multivariate genetic basis of correlated trait evolution, including whether single loci influence suites of behavioral, morphological, and physiological traits, and whether similar genetic modules recur across taxa. Large-scale mapping and association studies in multiple species, combined with comparative analyses of genetic covariance matrices, could reveal whether the color-behavior-physiology integration we observe reflects a broader, conserved genetic architecture.

## References

1. Altmann, J. 1974. Observational study of behavior: sampling methods. Behaviour 49:227–266. Brill.

2. Anderson, B. B., A. Scott, and R. Dukas. 2016. Social behavior and activity are decoupled in larval and adult fruit flies. Behav. Ecol. 27:820–828.

3. Anholt, R. R., and T. F. Mackay. 2015. Dissecting the genetic architecture of behavior in *Drosophila melanogaster*. Curr. Opin. Behav. Sci. 2:1–7.

4. Arnold, S. J., R. Bürger, P. A. Hohenlohe, B. C. Ajie, and A. G. Jones. 2008. Understanding the Evolution and Stability of the G-Matrix. Evol. Int. J. Org. Evol. 62:2451–2461.

5. Ashburner, M., and J. Roote. 2007. Maintenance of a Drosophila Laboratory: General Procedures. Cold Spring Harb. Protoc. 2007:pdb.ip35. Cold Spring Harbor Laboratory Press.

6. Austin, C. J., and A. J. Moehring. 2013. Optimal temperature range of a plastic species, *Drosophila simulans*. J. Anim. Ecol. 82:663–672.

7. Barghi, N., R. Tobler, V. Nolte, and C. Schlötterer. 2017. Drosophila simulans: A Species with Improved Resolution in Evolve and Resequence Studies. G3 GenesGenomesGenetics 7:2337–2343.

8. Bastide, H., D. Ogereau, C. Montchamp-Moreau, and P. R. Gérard. 2022. The fate of a suppressed X-linked meiotic driver: experimental evolution in Drosophila simulans. Chromosome Res. 30:141–150.

9. Bath, E., D. Edmunds, J. Norman, C. Atkins, L. Harper, W. G. Rostant, T. Chapman, S. Wigby, and J. C. Perry. 2021. Sex ratio and the evolution of aggression in fruit flies. Proc. R. Soc. B Biol. Sci. 288:20203053. Royal Society.

10. Baxter, C. M., and R. Dukas. 2017. Life history of aggression: effects of age and sexual experience on male aggression towards males and females. Anim. Behav. 123:11–20.

11. Blows, M., and B. Walsh. 2009. Spherical Cows Grazing in Flatland: Constraints to Selection and Adaptation. Pp. 83–101 *in* J. van der Werf, H.-U. Graser, R. Frankham, and C. Gondro, eds. Adaptation and Fitness in Animal Populations: Evolutionary and Breeding Perspectives on Genetic Resource Management. Springer Netherlands, Dordrecht.

12. Burke, M. K., J. P. Dunham, P. Shahrestani, K. R. Thornton, M. R. Rose, and A. D. Long. 2010. Genome-wide analysis of a long-term evolution experiment with Drosophila. Nature 467:587–590. Nature Publishing Group.

13. Burnet, B., L. Burnet, K. Connolly, and N. Williamson. 1988. A genetic analysis of locomotor activity in Drosophila melanogaster. Heredity 61:111–119. Nature Publishing Group.

14. Capy, P., and P. Gibert. 2004. Drosophila melanogaster, Drosophila simulans: so similar yet so different. Pp. 5–16 *in* P. Capy, P. Gibert, and I. Boussy, eds. Drosophila melanogaster, Drosophila simulans: So Similar, So Different. Springer Netherlands, Dordrecht.

15. Chebib, J., and F. Guillaume. 2021. Pleiotropy or linkage? Their relative contributions to the genetic correlation of quantitative traits and detection by multitrait GWA studies. Genetics 219:iyab159. Oxford University Press.

16. Chen, S., A. Y. Lee, N. M. Bowens, R. Huber, and E. A. Kravitz. 2002. Fighting fruit flies: A model system for the study of aggression. Proc. Natl. Acad. Sci. 99:5664–5668. National Academy of Sciences.

17. Cheverud, J. M. 1988. A Comparison of Genetic and Phenotypic Correlations. Evolution 42:958–968.

18. Cheverud, J. M. 1996. Developmental Integration and the Evolution of Pleiotropy. Am. Zool. 36:44–50. Oxford University Press.

19. Connallon, T., and M. D. Hall. 2018. Genetic constraints on adaptation: a theoretical primer for the genomics era. Ann. N. Y. Acad. Sci. 1422:65–87. Wiley-Blackwell.

20. David, J. R., P. Capy, V. Payant, and S. Tsakas. 1985. Thoracic trident pigmentation in Drosophila melanogaster: Differentiation of geographical populations. Génétique Sélection Évolution 17:211.

21. Dierick, H. A., and R. J. Greenspan. 2006. Molecular analysis of flies selected for aggressive behavior. Nat. Genet. 38:1023–1031. Nature Publishing Group.

22. Ducrest, A.-L., L. Keller, and A. Roulin. 2008. Pleiotropy in the melanocortin system, coloration and behavioural syndromes. Trends Ecol. Evol. 23:502–510.

23. Falconer, D. S., and T. F. C. Mackay. 1996. Introduction to quantitative genetics. Longmans Green, Harlow, Essex, UK.

24. Fernandez, M. P., S. Trannoy, and S. J. Certel. 2023. Fighting Flies: Quantifying and Analyzing Drosophila Aggression. Cold Spring Harb. Protoc. 2023:pdb.top107985. Cold Spring Harbor Laboratory Press.

25. Gaut, B. S., and A. D. Long. 2003. The Lowdown on Linkage Disequilibrium. Plant Cell 15:1502–1506.

26. Girardeau, A. R., G. E. Enochs, and J. B. Saltz. 2025. Evolutionary feedbacks for Drosophila aggression revealed through experimental evolution. Proc. Natl. Acad. Sci. 122:e2419068122. Proceedings of the National Academy of Sciences.

27. Hansen, T. F., and C. Pélabon. 2021. Evolvability: A Quantitative-Genetics Perspective. Annu. Rev. Ecol. Evol. Syst., doi: 10.1146/annurev-ecolsys-011121-021241.

28. Harbison, S. T., L. J. McCoy, and T. F. Mackay. 2013. Genome-wide association study of sleep in Drosophila melanogaster. BMC Genomics 14:281.

29. Hirsch, J. 1959. Studies in experimental behavior genetics. II. Individual differences in geotaxis as a function of chromosome variations in synthesized Drosophila populations. J. Comp. Physiol. Psychol. 52:304–308.

30. Hoffmann, A. A. 1987. A laboratory study of male territoriality in the sibling species *Drosophila melanogaster* and *D. simulans*. Anim. Behav. 35:807–818.

31. Hoffmann, A. A. 1991. Heritable Variation for Territorial Success in Field-Collected Drosophila melanogaster. Am. Nat. 138:668–679. The University of Chicago Press.

32. Houle, D. 1991. Genetic Covariance of Fitness Correlates: What Genetic Correlations Are Made of and Why It Matters. Evolution 45:630–648.

33. Houle, D., J. Mezey, P. Galpern, and A. Carter. 2003. Automated measurement of Drosophila wings. BMC Evol. Biol. 3:25.

34. Hutchins, M., T. Douglas, L. Pollack, and J. B. Saltz. 2024. Genetic Variation in Male Aggression Is Influenced by Genotype of Prior Social Partners in Drosophila melanogaster. Am. Nat. 203:551–561. The University of Chicago Press.

35. Jordan, K. W., M. A. Carbone, A. Yamamoto, T. J. Morgan, and T. F. Mackay. 2007. Quantitative genomics of locomotor behavior in Drosophila melanogaster. Genome Biol. 8:R172.

36. Jordan, K. W., T. J. Morgan, and T. F. C. Mackay. 2006. Quantitative Trait Loci for Locomotor Behavior in *Drosophila melanogaster*. Genetics 174:271–284.

37. Keene, A. C., E. R. Duboué, D. M. McDonald, M. Dus, G. S. B. Suh, S. Waddell, and J. Blau. 2010. Clock and cycle Limit Starvation-Induced Sleep Loss in Drosophila. Curr. Biol. 20:1209–1215. Elsevier.

38. Kelly, J. K., and K. A. Hughes. 2019. Pervasive Linked Selection and Intermediate-Frequency Alleles Are Implicated in an Evolve-and-Resequencing Experiment of *Drosophila simulans*. Genetics 211:943–961.

39. Koch, E. L., C. Rocabert, C. Beeravolu Reddy, and F. Guillaume. 2025. Gene expression evolution is predicted by stronger indirect selection at more pleiotropic genes. Evol. Lett. 9:719–730.

40. Kofler, R., and C. Schlötterer. 2014. A Guide for the Design of Evolve and Resequencing Studies. Mol. Biol. Evol. 31:474–483.

41. Kraft, B., V. A. Lemakos, J. Travis, and K. A. Hughes. 2018. Pervasive indirect genetic effects on behavioral development in polymorphic eastern mosquitofish. Behav. Ecol. 29:289–300.

42. Kravitz, E. A., and M. de la P. Fernandez. 2015. Aggression in Drosophila. Behav. Neurosci. 129:549–563. American Psychological Association, US.

43. Kravitz, E. A., and R. Huber. 2003. Aggression in invertebrates. Curr. Opin. Neurobiol. 13:736–743.

44. Krstevska, B., and A. A. Hoffmann. 1994. The effects of acclimation and rearing conditions on the response of tropical and temperate populations of *Drosophila melanogaster* and *D. simulans* to a temperature gradient (Diptera: Drosophilidae). J. Insect Behav. 7:279–288.

45. Lande, R. 1979. Quantitative Genetic Analysis of Multivariate Evolution, Applied to Brain: Body Size Allometry. Evolution 33:402–416. [Society for the Study of Evolution, Wiley].

46. Lande, R. 1980. The genetic covariance between characters maintained by pleiotropic mutations. Genetics 94:203–215. Oxford University Press.

47. Lank, D. B., C. M. Smith, O. Hanotte, T. Burke, and F. Cooke. 1995. Genetic polymorphism for alternative mating behaviour in lekking male ruff *Philomachus pugnax*. Nature 378:59–62. Nature Publishing Group.

48. Leinonen, T., J. M. Cano, and J. Merilä. 2011. Genetics of body shape and armour variation in threespine sticklebacks. J. Evol. Biol. 24:206–218.

49. Lenth, R. 2025. emmeans: Estimated Marginal Means, aka Least-Squares Means.

50. Mallard, F., B. Afonso, and H. Teotónio. 2023. Selection and the direction of phenotypic evolution. eLife 12:e80993. eLife Sciences Publications, Ltd.

51. Martin, J.-R. 2003. Locomotor activity: a complex behavioural trait to unravel. Behav. Processes 64:145–160.

52. Massey, J. H., and P. J. Wittkopp. 2016. The Genetic Basis of Pigmentation Differences Within and Between Drosophila Species. Curr. Top. Dev. Biol. 119:27–61.

53. McGraw, K. J., J. Dale, and E. A. Mackillop. 2003. Social environment during molt and the expression of melanin-based plumage pigmentation in male house sparrows (*Passer domesticus*). Behav. Ecol. Sociobiol. 53:116–122.

54. Murphey, R. M., and C. F. Hall. 1969. Some correlates of negative geotaxis in *Drosophila melanogaster*. Anim. Behav. 17:181–185.

55. Obbard, D. J., J. Maclennan, K.-W. Kim, A. Rambaut, P. M. O’Grady, and F. M. Jiggins. 2012. Estimating Divergence Dates and Substitution Rates in the Drosophila Phylogeny. Mol. Biol. Evol. 29:3459–3473.

56. Pfeiffenberger, C., B. C. Lear, K. P. Keegan, and R. Allada. 2010a. Locomotor Activity Level Monitoring Using the Drosophila Activity Monitoring (DAM) System. Cold Spring Harb. Protoc. 2010:pdb.prot5518. Cold Spring Harbor Laboratory Press.

57. Pfeiffenberger, C., B. C. Lear, K. P. Keegan, and R. Allada. 2010b. Processing Sleep Data Created with the Drosophila Activity Monitoring (DAM) System. Cold Spring Harb. Protoc. 2010:pdb.prot5520. Cold Spring Harbor Laboratory Press.

58. Podos, J. 2001. Correlated evolution of morphology and vocal signal structure in Darwin’s finches. Nature 409:185–188. Nature Publishing Group.

59. R Core Team. 2025. R: A Language and Environment for Statistical Computing. R Foundation for Statistical Computing, Vienna, Austria.

60. Rajpurohit, S., L. M. Peterson, A. J. Orr, A. J. Marlon, and A. G. Gibbs. 2016. An Experimental Evolution Test of the Relationship between Melanism and Desiccation Survival in Insects. PLOS ONE 11:e0163414. Public Library of Science.

61. Remolina, S. C., P. L. Chang, J. Leips, S. V. Nuzhdin, and K. A. Hughes. 2012. GENOMIC BASIS OF AGING AND LIFE-HISTORY EVOLUTION IN DROSOPHILA MELANOGASTER. Evolution 66:3390–3403.

62. Rohwer, S. 1975. The Social Significance of Avian Winter Plumage Variability. Evolution 29:593–610. [Society for the Study of Evolution, Wiley].

63. Roulin, A., and A.-L. Ducrest. 2011. Association between melanism, physiology and behaviour: A role for the melanocortin system. Eur. J. Pharmacol. 660:226–233.

64. Ruckman, S. N., E. A. Humphrey, L. Muzzey, I. Prantalou, M. Pleasants, and K. A. Hughes. 2024. Assessing the Association Between Animal Color and Behavior: A Meta-Analysis of Experimental Studies. Ecol. Evol. 14:e70655.

65. Sacchi, R., A. J. Coladonato, M. Battaiola, C. Pasquariello, S. Buratti, C. Matellini, M. Mangiacotti, S. Scali, and M. A. L. Zuffi. 2021. Subjective resource value affects aggressive behavior independently of resource-holding-potential and color morphs in male common wall lizard. J. Ethol. 39:179–189.

66. San-Jose, L. M., and A. Roulin. 2018. Toward Understanding the Repeated Occurrence of Associations between Melanin-Based Coloration and Multiple Phenotypes. Am. Nat. 192:111–130. The University of Chicago Press.

67. Scali, S., M. Mangiacotti, R. Sacchi, A. J. Coladonato, M. Falaschi, L. Saviano, M. G. Rampoldi, M. Crozi, C. Perotti, F. Zucca, E. Gozzo, and M. A. L. Zuffi. 2021. Close encounters of the three morphs: Does color affect aggression in a polymorphic lizard? Aggress. Behav. 47:430–438.

68. Schneider, C. A., W. S. Rasband, and K. W. Eliceiri. 2012. NIH Image to ImageJ: 25 years of image analysis. Nat. Methods 9:671–675. Nature Publishing Group.

69. Sinervo, B., and C. M. Lively. 1996. The rock–paper–scissors game and the evolution of alternative male strategies. Nature 380:240–243. Nature Publishing Group.

70. Slatkin, M. 2008. Linkage disequilibrium — understanding the evolutionary past and mapping the medical future. Nat. Rev. Genet. 9:477–485. Nature Publishing Group.

71. Sokolowski, M. B. 2001. Drosophila: Genetics meets behaviour. Nat. Rev. Genet. 2:879–890. Nature Publishing Group.

72. Steppan, S. J., P. C. Phillips, and D. Houle. 2002. Comparative quantitative genetics: evolution of the G matrix. Trends Ecol. Evol. 17:320–327. Elsevier.

73. Suh, J., and F. R. Jackson. 2007. Drosophila Ebony Activity Is Required in Glia for the Circadian Regulation of Locomotor Activity. Neuron 55:435–447. Elsevier.

74. Takahashi, A. 2013. Pigmentation and behavior: potential association through pleiotropic genes in *Drosophila*. Genes Genet. Syst. 88:165–174.

75. Taylor, M. J., and R. I. Tuxworth. 2019. Continuous tracking of startled Drosophila as an alternative to the negative geotaxis climbing assay. J. Neurogenet. 33:190–198. Taylor & Francis.

76. Therneau, T. M. 2024. coxme: Mixed Effects Cox Models.

77. Tibbetts, E. A., and J. Dale. 2004. A socially enforced signal of quality in a paper wasp. Nature 432:218–222. Nature Publishing Group.

78. Toma, D. P., K. P. White, J. Hirsch, and R. J. Greenspan. 2002. Identification of genes involved in Drosophila melanogaster geotaxis, a complex behavioral trait. Nat. Genet. 31:349–353. Nature Publishing Group.

79. Turner, T. L., P. M. Miller, and V. A. Cochrane. 2013. Combining Genome-Wide Methods to Investigate the Genetic Complexity of Courtship Song Variation in Drosophila melanogaster. Mol. Biol. Evol. 30:2113–2120.

80. Turner, T. L., A. D. Stewart, A. T. Fields, W. R. Rice, and A. M. Tarone. 2011. Population-Based Resequencing of Experimentally Evolved Populations Reveals the Genetic Basis of Body Size Variation in Drosophila melanogaster. PLOS Genet. 7:e1001336. Public Library of Science.

81. Ueda, A., and Y. Kidokoro. 2002. Aggressive behaviours of female Drosophila melanogaster are influenced by their social experience and food resources. Physiol. Entomol. 27:21–28.

82. Vlachos, C., and R. Kofler. 2019. Optimizing the Power to Identify the Genetic Basis of Complex Traits with Evolve and Resequence Studies. Mol. Biol. Evol. 36:2890–2905.

83. Wagner, G. P., and J. Zhang. 2011. The pleiotropic structure of the genotype–phenotype map: the evolvability of complex organisms. Nat. Rev. Genet. 12:204–213. Nature Publishing Group.

84. Walsh, B., and M. W. Blows. 2009. Abundant Genetic Variation + Strong Selection = Multivariate Genetic Constraints: A Geometric View of Adaptation. Annu. Rev. Ecol. Evol. Syst. 40:41–59.

85. Walter, G. M. 2023. Experimental Evidence That Phenotypic Evolution but Not Plasticity Occurs along Genetic Lines of Least Resistance in Homogeneous Environments. Am. Nat. 201:E70–E89. The University of Chicago Press.

86. Weber, K. E. 1996. Large Genetic Change at Small Fatness Cost in Large Populations of Drosophila melanogaster Selected for Wind Tunnel Flight: Rethinking Fitness Surfaces. Genetics 144:205–213.

87. Wittkopp, P. J., and P. Beldade. 2009. Development and evolution of insect pigmentation: Genetic mechanisms and the potential consequences of pleiotropy. Semin. Cell Dev. Biol. 20:65–71.

88. Wittkopp, P. J., S. B. Carroll, and A. Kopp. 2003. Evolution in black and white: genetic control of pigment patterns in Drosophila. Trends Genet. 19:495–504.

89. Wittkopp, P. J., G. Smith-Winberry, L. L. Arnold, E. M. Thompson, A. M. Cooley, D. C. Yuan, Q. Song, and B. F. McAllister. 2011. Local adaptation for body color in Drosophila americana. Heredity 106:592–602. Nature Publishing Group.

90. Yoo, B. H. 1980. Long-term selection for a quantitative character in large replicate populations of Drosophila melanogaster: 1. Response to selection. Genet. Res. 35:1–17.

91. Zhong, L., Z. Yang, H. Tang, Y. Xu, X. Liu, and J. Shen. 2022. Differential analysis of negative geotaxis climbing trajectories in Drosophila under different conditions. Arch. Insect Biochem. Physiol. 111:e21922.

