## Supplemental Text for "Behavior evolves as a correlated response to selection on cuticle color in *Drosophila melanogaster* and *D. simulans*"

**Supplemental Methods**

***Source populations***

We used *D. melanogaster* derived from two different sources. Population 1 (*D. mel-*1) descended from flies originally caught in Archibald Orchard (43.99816567881053, -78.70830481010981), Clarington, Canada, in the summer of 2021 by J. Sztepanacz (personal communication with J. Sztepanacz). A laboratory population was established from 600-1000 mated wild-caught females and subsequently maintained at a large population size (800 individuals). Population 2 (*D. mel-*2) descended from flies originally collected in Coral Gables, FL, USA, by B. de Bivort. The collection methods are published in Akhund-Zade et al. 2020. The laboratory population was established from one hundred mated wild-caught females, which were kept at a large population size (250 individuals).

We established two *D. simulans* populations from wild flies collected between February and May 2023 in Tallahassee, FL, USA (coordinates: 30.46, -84.22). Population 1 (*D. sim-*1) was established from wild-caught flies collected between February and March of 2023. Population 2 (*D. sim-*2) was established from wild-caught flies collected between April and May 2023. In *D. melanogaster*, hundreds of polymorphisms undergo shifts in allele frequency and oscillate over seasons (Bergland et al. 2014). Consequently, although these two populations were established from flies collected from the exact geographical location, they likely had different initial allele frequencies because of temporal variation in the wild population and sampling effects. These flies were collected by sweep netting across a compost pile every other day for 30 minutes after sunrise (40 days). After collecting for the day, the individuals were taken into the lab, and all females were isolated in vials with the standard media. Visual species identification of females is challenging to perform in these species, so we assessed their male offspring for species identification (Coyne 1983). Females were allowed to lay eggs for 48 hours before being removed. After the offspring emerged at 12 days, the vials were sexed, and the males were used to determine the species of the individuals. If the vial only contained *D. simulans* males, then the offspring were released into a population cage to mix with the offspring of other females. Population 1 was created from the offspring of 265 mated females. Population 2 was created from the offspring of 235 mated females.

***Color Selection by Eye***

To perform selection, we visually compared the dorsal thorax between the macrochaeta of the individuals by eye in sets of 50 by anesthetizing them with CO_2_ and then categorizing them into four groups based on the color and appearance of their thorax, as described in David et al. (1985). The four categories included: 0) no trident; 1) faint trident; 2) trident clearly marked; 3) dark full trident on the fly (Figure 1 in David et al. (1985). Individuals were sorted into either category 0 (not having a trident) or category 1 (having a trident), and then all flies that had a trident were further sorted into categories 1 through 3 based on how light or dark the trident was. SNR was the only person who conducted the color selection.

To validate the visual binning method used for cuticle coloration, we assessed both its agreement with an objective color measurement and its repeatability over time. We photographed the dorsal thorax of 100 individuals and used ImageJ to extract mean grayscale intensity values from the standardized region between macrochaeta landmarks (described in full detail below). These quantitative color measurements served as an objective baseline to compare against the visual bin assignments (0–3 categories). We tested agreement between methods using a general linear model with grayscale value as the response variable and visual bin as a fixed effect. We found a significant difference between the bins (Table S27; Figure S1). To determine which bins were significantly different from one another, we used a pairwise t-test with the Bonferroni correction. To do this, we used the emmeans package, which reports the least squares means and conducts the pairwise t-tests (Lenth 2025). We used the emmeans function, with the fixed effect of bin, and the “cov.reduce” command equaling false to separate the bins rather than averaging them (Lenth 2025). Then we used the pairs function with the Bonferroni correction to print the paired comparisons. We found that the bins were significantly different from one another in the expected directions (Table S28; Figure S1). This analysis indicates that the visual binning reliably reflects variation in actual color. Second, to test repeatability, the same observer scored 100 individuals twice at different time points. To test the repeatability of visual binning methods, we calculated the intraclass correlation coefficient (ICC) using a two-way mixed-effects model for absolute agreement to quantify consistency between the two scoring sessions using the R package irr (Gamer et al. 2019). The ICC value was high (ICC = 0.933; F_99,99.1_ = 29.2, p < 0.001), indicating strong repeatability of the visual binning method over time (Figure S2). These validations demonstrate that the binning method is an accurate and reliable measure of cuticle coloration for selection for the labor and time-intensive selection procedure.

***Color Quantification of Individuals Used in Behavioral Assays***

To quantify color more precisely, we photographed all individuals used in the behavioral tests (described below) within 3 days of behavioral trials. To photograph the individuals, we used the WINGMACHINE (Houle et al. 2003). In brief, this apparatus uses a vacuum to hold an individual’s wings in place. We anesthetized the individual with CO_2_ and then used the vacuum to hold the fly in position to take a photo of the dorsal thorax of the fly. Each fly was photographed with a black and white color standard (WhiBal G7 WB7-PC) and a 2 mm scale within the field of view.

To quantify the color of the dorsal thorax, we used ImageJ to find the mean grey scale value (Schneider et al. 2012). First, we converted the image to 8-bit from RGB color (Figure S3a). Then, we calibrated the image to the black and white color standard. To do this, we measured the black and white portion of the color standard below the fly. Using the calibrate option in ImageJ, we used the “straight line” function and gave the black and white values known values of 0 and 255, respectively. Then, using the freeform tool in ImageJ, we used the bristle attachment sites as landmarks to measure the average color of the dorsal thorax over the entire area (Figure S3b). Two individuals analyzed each photo, and grey scale values were averaged between scorers (average standard deviation = 6.536).

**Behavioral Assays**

***Aggression:*** The arena consisted of a 35 mm petri dish wrapped with plastic wrap. The arena floor was filled with standard media, and we placed a dot of active yeast paste in the center of the arena (Girardeau et al. 2025). Following a five-minute acclimation period, we employed an instantaneous scan sampling approach (Altmann 1974), where every five minutes for one hour, we recorded the occurrence of fencing, lunging, boxing, and tussling (Chen et al. 2002). Fencing occurs when two flies extend one foreleg each to strike or push the opponent. Lunging occurs when a fly rapidly rears up on its hind legs and snaps down onto the opponent with its forelegs. Boxing consists of both flies rearing up and exchanging repeated foreleg strikes while maintaining an upright posture. Tussling involves the flies locking together with their legs and often rolling or pushing each other across the substrate while in close contact. These behaviors are seen in both sexes; however, boxing and tussling are rare in females (Chen et al. 2002; Ueda and Kidokoro 2002). At the end of the trial, all focal individuals were housed in their arenas until they were photographed (described above). We tested 100 males and 100 females from each treatment per population per species for generations 0, 5, and 10. At generation 15, we only tested 30 males and 30 females from each treatment per population per species because of logistical reasons associated with testing multiple behaviors toward the end of the experiment for each population.

***Basal Activity Level:*** The open field arena consisted of a 60 mm petri dish with a 1x1 cm grid on the bottom. A Logitech Brio 101 (960-001589) HD webcam at 1080p was positioned 8 inches above the petri dish. Basal activity trials were only conducted at generation 12. We tested 50 males and 50 females from each treatment per population per species.

***Geotaxis Level:*** The geotaxis arena consisted of a jumbo clear plastic straw (Comfy package 10 mm diameter straws) taped on one end and plugged with cotton at the other. The straws were marked 1 cm from the bottom to designate the “starting line” and marked 3 cm from the bottom to designate the “finish line”. Geotaxis trials were conducted at generation 13. We tested 50 males and 50 females from each treatment per population per species.

***Total Activity Level and Sleep:*** To gain a more detailed understanding of the differences in activity level changes over a longer time period, we used the DAM system. The DAM system detects activity by monitoring infrared beam crossings of individually housed flies. To determine the activity level of the flies, we used the *Drosophila* Sleep Counting Macro (Pfeiffenberger et al. 2010a,b). In brief, this macro identifies 5-minute bouts and counts the number of beam breaks during that period. To load flies into the apparatus, flies were briefly anesthetized with CO_2_ and loaded individually into DAM tubes with standard media at one end. Flies were then acclimated for at least 24 hours before the start of behavioral trials. After the acclimation period, total activity was measured for 72 hours. We measured activity as total activity. Total activity is measured as the total number of beams crossed per day. We also measured total sleep as the time that the flies were inactive. The individuals needed to be inactive for at least 5 minutes to be considered asleep before the system would start counting sleep time. All measurements were averaged over the three-day experimental period. We tested 62 males and 62 females from each treatment per population and species. These experiments were performed in a humidified incubator at 25ºC and 65% humidity on a 12-hour light/dark cycle (Percival Scientific). Placement of the DAM apparatus within the incubator was randomized with respect to treatment. However, each DAM apparatus contained only one treatment group, and within each apparatus, females were always loaded before males.

***Statistical analyses – Direct and Correlated Responses to Selection on Thorax Color***

To test for responses to selection at the final generations of measurement, we used a consistent generalized linear model (GLM) framework. Critically, all models were analyzed separately for males and females because the residual distributions differ between the sexes. For all models, we visually inspected the residuals to ensure that model assumptions (e.g., normality, homoscedasticity, goodness of fit) were met. The full model for each trait related the dependent variable (e.g., grayscale value, aggression counts) to the following fixed effects: species, treatment, population (nested within species), and all two-way interactions. To test our hypothesis that selection treatments would diverge linearly, we treated the predictor “treatment” as a continuous variable with values -1, 0, and 1, corresponding to L, C, and D treatments, respectively. This convention allowed us to test for both the direction and statistical significance of the evolutionary response to selection. We employed a maximum likelihood approach and generated F-test or Wald Chi-square (X²) statistics to evaluate the significance of each main and interaction effect. If interaction terms were not significant (p > 0.05), they were dropped from the model to test the main effects.

If a significant main effect or interaction involving “treatment” was found, we conducted post-hoc tests to determine which pairs of treatments differed significantly using the emmeans package in R, which reports the least squares means and conducts pairwise t-tests (Lenth 2025). If we found a significant interaction between treatment and population, we then performed a simple slopes analysis to understand how the response to selection varied among populations. This post-hoc test estimates the slope of the evolutionary response for each population individually and Wald Chi-square tests to determine significance. These tests were conducted using the emtrends function in the emmeans package (Lenth 2025). All statistical analyses were conducted using R version 4.5.0 (R Core Team 2025).

***Color Analysis:*** To test for a direct response to selection at generation 15, we analyzed the mean grayscale values using a general linear model assuming a normal distribution.

***Aggression Analysis:*** To determine how selection affected aggressive behavior at generation 15, we analyzed the counts of aggressive acts using a generalized linear model assuming a Poisson distribution and using a log link function. Since generalized linear models with Poisson distributions do not estimate residual variance, we used asymptotic z-tests with infinite degrees of freedom to conduct the post-hoc pairwise comparisons.

***Activity Level and Sleep Analyses:*** For basal activity at generation 12, we analyzed line-crossing counts using a generalized linear model with a negative binomial distribution. For activity at generation 16, we analyzed the average total sleep and total activity using a general linear model assuming a normal distribution.

***Geotaxis Analysis:*** For the time-to-event geotaxis data, we used a Cox proportional hazards mixed-effects model, implemented in the coxme package in R (Therneau 2024), with a random effect of observer. This model estimates the risk of completing the behavioral task over time, while accounting for both fixed effects and random effects. Each geotaxis yielded two observations: 1) the time that the fly took to cross the second line, and 2) whether they completed the trial in the allotted time or not. If flies took longer than one minute to cross the second line, they were censored and coded as 60 seconds in the trial and 0 for the censoring value. Of 1,218 total trials, 60 (4.9%) were censored.

***Statistical Analyses – Dynamics of Selection Responses Over 15 Generations***

To examine temporal patterns in direct and indirect responses to selection, we used statistical models similar to those described above with the addition of generation and the square of generation as predictors. We included interactions between generation and treatment to assess whether evolutionary trajectories differed between selection treatments. For thorax color, we used a general linear model assuming a normal distribution; for the aggression data, we assumed a Poisson distribution with a log link function.

**Supplemental Results**

***Correlated Evolution of Geotaxis***

At generation 13, the correlated response in geotaxis was inconsistent, varying unpredictably across populations for both sexes (Figures S12 and S13). This was reflected in a significant treatment by population interaction (males: X^2^_2_ = 23.5, p < 0.0001; females: X^2^_2_ = 36.7, p < 0.0001; Table S15). The evolutionary trajectories differed among groups and were often contradictory (Table S16). For example, among males, there was a significant response in *D. mel*-2 and *D. sim*-1, but non-significant responses in others (Table S16). Among females, the response was even more varied: the slope was positive in *D. mel*-2 but significantly negative (indicating a slower response) in *D. mel*-1 and *D. sim*-2 (Table S16). Post-hoc tests confirmed this lack of a consistent pattern (Table S17). For instance, in *D. mel*-1 males, both D and L selected lines evolved to be faster than controls (Table S17; Figure S12), while in *D. mel*-2 females, D flies evolved to be significantly slower than L and C flies (Table S17; Figure S13). Many other pairwise comparisons were not significant, reinforcing that geotaxis did not evolve predictably with color.

***Correlated Evolution of Total Activity Level***

At generation 16, the correlated response in total activity was complex and inconsistent in direction, differing between species in males and among populations in females (Figures S14 and S15). In males, a significant treatment by species interaction (F_1.699_ = 17.7, p < 0.0001; Table S18) indicated that the two species responded differently. As predicted, the response for *D. melanogaster* was significantly positive (slope = 108.9, X^2^_1_ = 17.6, p < 0.0001; Table S19), with post-hoc tests showing D males were more active than L males (Table S20). In contrast, *D. simulans* showed no significant response to selection (Table S20).

The response in females was even more varied, reflected in a significant three-way interaction between treatment, population, and species (F_1.719_ = 6.01, p = 0.0023; Table S18). The two *D. melanogaster* populations evolved in opposite directions: slope was significantly positive in *D. mel*-1 but significantly negative in *D. mel*-2 (46.8 vs. -48.0, respectively; Table S19). The only significant pairwise difference was in *D. mel-2*, where C females were less active than both D and L females (Table S20). No other post-hoc tests were significant (Table S20), reinforcing the conclusion that total activity did not evolve predictably.

***Correlated Evolution of Sleep***

At generation 16, the correlated response for sleep was inconsistent and complex in direction, particularly in males (Figures S16 and S17). In males, a significant treatment by population interaction (F_2.697_ = 11.2, p < 0.0001; Table S21) confirmed that the response to selection was highly variable. The evolutionary trajectories were contradictory: the slope was significantly negative in *D. mel*-2 (indicating less sleep in dark-selected lines, as predicted), but significantly positive in *D. sim*-1 indicating more sleep (Table S22). Post-hoc tests reflected this complexity; for example, D males in *D. mel*-2 slept significantly less than C, whereas D males in *D. sim*-1 slept significantly more than L males (Table S23; Figure S16). In females, the overall linear slope for the treatment effect on sleep was not significant (slope = 2.7, X^2^_1_ = 0.09, p = 0.76; Table S22). However, post-hoc tests revealed a significant, non-linear response to selection (Table S23; Figure S17). Specifically, both D and L females slept significantly less than C females. The D and L treatments, however, were not significantly different from each other (Table S23; Figure S17).

***Phenotypic Correlations between Color and Behavior***

Within-population color-aggression correlations were significant in *D. melanogaster* (rho = -0.063, rho² = 0.004, p < 0.001; Table S24) but not *D. simulans* (rho = -0.008, p = 0.631; Table S24). Correlations were significant in males (rho = -0.050, rho² = 0.003, p = 0.004; Table S24) but not females (rho = -0.022, p = 0.194; Table S24). By line, the D line showed a significant negative correlation (rho = -0.082, rho² = 0.007, p = 0.001; Table S24), while C and L lines did not (p = 0.086 and p = 0.080, respectively; Table S24). Within-population correlations between color and basal activity were not significant in any species, sex, or line breakdown (all p > 0.05; Table S24).

Within-population color-geotaxis correlations were significant in *D. simulans* (rho = -0.204, rho² = 0.042, p < 0.001; Table S24) but not *D. melanogaster* (rho = 0.013, p = 0.746; Table S24). Correlations were significant in males (rho = -0.173, rho² = 0.030, p < 0.001; Table S24) but not females (rho = -0.002, p = 0.955; Table S24). The C line showed a significant negative correlation (rho = -0.192, rho² = 0.037, p < 0.001; Table S24), while D and L lines did not (p = 0.408 and p = 0.518, respectively; Table S24).

Pairwise Fisher z-tests comparing generation 0 to generation 15 revealed significant shifts in color-aggression correlations within the D line overall (rho = 0.011 to rho = -0.168, p = 0.010; Table S25), driven by *D. simulans* (rho = 0.056 to rho = -0.203, p = 0.009; Table S25) and particularly *D. simulans* females (rho = 0.081 to rho = -0.327, p = 0.003; Table S25). Dark line females overall also shifted significantly (rho = 0.029 to rho = -0.176, p = 0.037; Table S25). In the L line, *D. simulans* showed a significant shift (rho = 0.056 to rho = -0.141, p = 0.046; Table S25), driven by females (rho = 0.081 to rho = -0.224, p = 0.027; Table S25).

Correlations between mean aggression and mean basal activity were significant overall when aggression was averaged across generations 5, 10, and 15 (rho = 0.52, rho² = 0.27, p = 0.004; Table S26). This association was strongest in the D line (rho = 0.857, rho² = 0.734, p < 0.001; Table S26). By population, *D. mel*-2 (rho = 0.829, p = 0.003; Table S26), *D. sim*-1 (rho = 0.743, p = 0.027; Table S26), and *D. sim*-2 (rho = 0.714, p = 0.041; Table S26) all showed significant positive correlations. *D. mel*-1 showed a positive but non-significant trend (rho = 0.657, p = 0.081; Table S26).

**References**

Akhund-Zade, J., D. Yoon, A. Bangerter, N. Polizos, M. Campbell, A. Soloshenko, T. Zhang, E. Wice, A. Albright, A. Narayanan, P. Schmidt, J. Saltz, J. Ayroles, M. Klein, A. Bergland, and B. de Bivort. 2020. Wild flies hedge their thermal preference bets in response to seasonal fluctuations. bioRxiv.

Altmann, J. 1974. Observational study of behavior: sampling methods. Behaviour 49:227–266. Brill.

Bergland, A. O., E. L. Behrman, K. R. O’Brien, P. S. Schmidt, and D. A. Petrov. 2014. Genomic Evidence of Rapid and Stable Adaptive Oscillations over Seasonal Time Scales in Drosophila. PLOS Genet. 10:e1004775. Public Library of Science.

Chen, S., A. Y. Lee, N. M. Bowens, R. Huber, and E. A. Kravitz. 2002. Fighting fruit flies: A model system for the study of aggression. Proc. Natl. Acad. Sci. 99:5664–5668. National Academy of Sciences.

Coyne, J. A. 1983. Genetic Basis of Differences in Genital Morphology Among Three Sibling Species of Drosophila. Evolution 37:1101–1118. [Society for the Study of Evolution, Wiley].

David, J. R., P. Capy, V. Payant, and S. Tsakas. 1985. Thoracic trident pigmentation in Drosophila melanogaster: Differentiation of geographical populations. Génétique Sélection Évolution 17:211.

Gamer, M., J. Lemon, and I. F. P. Singh. 2019. irr: Various Coefficients of Interrater Reliability and Agreement.

Girardeau, A. R., G. E. Enochs, and J. B. Saltz. 2025. Evolutionary feedbacks for Drosophila aggression revealed through experimental evolution. Proc. Natl. Acad. Sci. 122:e2419068122. Proceedings of the National Academy of Sciences.

Houle, D., J. Mezey, P. Galpern, and A. Carter. 2003. Automated measurement of Drosophila wings. BMC Evol. Biol. 3:25.

Lenth, R. 2025. emmeans: Estimated Marginal Means, aka Least-Squares Means.

Pfeiffenberger, C., B. C. Lear, K. P. Keegan, and R. Allada. 2010a. Locomotor Activity Level Monitoring Using the Drosophila Activity Monitoring (DAM) System. Cold Spring Harb. Protoc. 2010:pdb.prot5518. Cold Spring Harbor Laboratory Press.

Pfeiffenberger, C., B. C. Lear, K. P. Keegan, and R. Allada. 2010b. Processing Sleep Data Created with the Drosophila Activity Monitoring (DAM) System. Cold Spring Harb. Protoc. 2010:pdb.prot5520. Cold Spring Harbor Laboratory Press.

R Core Team. 2025. R: A Language and Environment for Statistical Computing. R Foundation for Statistical Computing, Vienna, Austria.

Schneider, C. A., W. S. Rasband, and K. W. Eliceiri. 2012. NIH Image to ImageJ: 25 years of image analysis. Nat. Methods 9:671–675. Nature Publishing Group.

Therneau, T. M. 2024. coxme: Mixed Effects Cox Models.

Ueda, A., and Y. Kidokoro. 2002. Aggressive behaviours of female Drosophila melanogaster are influenced by their social experience and food resources. Physiol. Entomol. 27:21–28.
