## Supplemental Tables and Figures for "Behavior evolves as a correlated response to selection on cuticle color in *Drosophila melanogaster* and *D. simulans*"

Table S1: Results from generalized linear model on the rate of thorax color evolution during the selection experiment. Italics represent significant differences.

| **Males:** | | | | |
| --- | --- | --- | --- | --- |
| Effect | F Values | df-num | df-dem | p-value |
| *Treatment* | *22.95* | *1* | *2324* | *1.8e-6* |
| *Species* | *18.19* | *1* | *2324* | *2.1e-5* |
| *Generation* | *17.13* | *1* | *2324* | *3.6e-5* |
| *Generation^2^* | *31.43* | *1* | *2324* | *2.3e-8* |
| *Population nested in Species* | *24.92* | *2* | *2324* | *2.0e-11* |
| Treatment x Species | 0.14 | 1 | 2324 | 0.71 |
| *Treatment x Generation* | *15.32* | *1* | *2324* | *9.2e-5* |
| *Treatment x Generation^2^* | *42.73* | *1* | *2324* | *7.7e-11* |
| Species x Generation | 0.80 | 1 | 2324 | 0.37 |
| Species x Generation^2^ | 1.21 | 1 | 2324 | 0.27 |
| *Treatment x Population nested in Species* | *12.07* | *2* | *2324* | *6.1e-6* |
| *Generation x Population nested in Species* | *13.51* | *2* | *2324* | *1.5e-6* |
| *Generation^2^ x Population nested in Species* | *8.08* | *2* | *2324* | *0.0003* |
| **Females:** | | | | |
| Treatment | 2.54 | 1 | 2567 | 0.11 |
| *Species* | *10.15* | *1* | *2567* | *0.0015* |
| Generation | 2.23 | 1 | 2567 | 0.14 |
| Generation^2^ | 2.72 | 1 | 2567 | 0.10 |
| *Population nested in Species* | *3.40* | *2* | *2567* | *0.033* |
| Treatment x Species | 0.014 | 1 | 2567 | 0.91 |
| *Treatment x Generation* | *11.55* | *1* | *2567* | *0.0007* |
| Treatment x Generation^2^ | 0.74 | 1 | 2567 | 0.39 |
| *Species x Generation* | *12.60* | *1* | *2567* | *0.0004* |
| *Species x Generation^2^* | *8.53* | *1* | *2567* | *0.0035* |
| Treatment x Population nested in Species | 0.22 | 2 | 2567 | 0.80 |
| *Generation x Population nested in Species* | *8.0* | *2* | *2567* | *0.0003* |
| *Generation^2^ x Population nested in Species* | *11.03* | *2* | *2567* | *1.7e-5* |

Table S2: Results of pairwise t-tests of color data over the course of the experiment. Italics represent significant differences.

| **Males:** | | | | | | | | |
| --- | --- | --- | --- | --- | --- | --- | --- | --- |
| Species | Pop | Gen | Contrast | estimate | SE | t-ratio | df | p-value |
| *D. mel* | *1* | *5* | *C-D* | *17.86* | *2.06* | *8.68* | *2318* | *< 0.0001* |
| D. mel | 1 | 5 | C-L | 1.36 | 2.06 | 0.66 | *2318* | 1.0 |
| *D. mel* | *1* | *5* | *D-L* | *-16.50* | *2.07* | *-7.96* | *2318* | *< 0.0001* |
| *D. mel* | *2* | *5* | *C-D* | *10.99* | *2.36* | *4.65* | *2318* | *0.0019* |
| D. mel | 2 | 5 | C-L | -4.82 | 2.44 | -1.98 | 2318 | 0.99 |
| *D. mel* | *2* | *5* | *D-L* | *-15.81* | *2.55* | *-6.20* | *2318* | *< 0.0001* |
| *D. sim* | *1* | *5* | *C-D* | *14.02* | *2.20* | *6.38* | *2318* | *< 0.0001* |
| D. sim | 1 | 5 | C-L | -3.97 | 2.16 | -1.84 | 2318 | 1.0 |
| *D. sim* | *1* | *5* | *D-L* | *-17.98* | *2.14* | *-8.39* | *2318* | *< 0.0001* |
| *D. sim* | *2* | *5* | *C-D* | *-12.36* | *2.21* | *-5.60* | 2318 | *< 0.0001* |
| *D. sim* | *2* | *5* | *C-L* | *-17.43* | *2.43* | *-7.17* | 2318 | *< 0.0001* |
| D. sim | 2 | 5 | D-L | -5.07 | 2.34 | -2.16 | 2318 | 0.97 |
| *D. mel* | *1* | *10* | *C-D* | *25.68* | *2.05* | *12.50* | 2318 | *< 0.0001* |
| D. mel | 1 | 10 | C-L | 0.24 | 2.05 | 0.12 | 2318 | 1.0 |
| *D. mel* | *1* | *10* | *D-L* | *-25.44* | *2.06* | *-12.38* | 2318 | *< 0.0001* |
| *D. mel* | *2* | *10* | *C-D* | *18.80* | *2.22* | *8.46* | 2318 | *< 0.0001* |
| D. mel | 2 | 10 | C-L | -5.95 | 2.24 | -2.65 | 2318 | 0.76 |
| *D. mel* | *2* | *10* | *D-L* | *-24.74* | *2.17* | *-11.38* | 2318 | *< 0.0001* |
| *D. sim* | *1* | *10* | *C-D* | *21.83* | *2.38* | *9.17* | 2318 | *< 0.0001* |
| D. sim | 1 | 10 | C-L | -5.09 | 2.22 | -2.30 | 2318 | 0.94 |
| *D. sim* | *1* | *10* | *D-L* | *-26.92* | *2.32* | *-11.58* | 2318 | *< 0.0001* |
| D. sim | 2 | 10 | C-D | -4.54 | 2.08 | -2.18 | 2318 | 0.97 |
| *D. sim* | *2* | *10* | *C-L* | *-18.55* | *2.15* | *-8.612* | 2318 | *< 0.0001* |
| *D. sim* | *2* | *10* | *D-L* | *-14.01* | *2.13* | *-6.59* | 2318 | *< 0.0001* |
| *D. mel* | *1* | *15* | *C-D* | *38.51* | *2.87* | *13.42* | 2318 | *< 0.0001* |
| *D. mel* | *1* | *15* | *C-L* | *-23.05* | *2.86* | *-8.05* | 2318 | *< 0.0001* |
| *D. mel* | *1* | *15* | *D-L* | *-61.55* | *2.88* | *-21.36* | 2318 | *< 0.0001* |
| *D. mel* | *2* | *15* | *C-D* | *31.63* | *2..95* | *10.73* | 2318 | *< 0.0001* |
| *D. mel* | *2* | *15* | *C-L* | *-29.23* | *2.97* | *-9.84* | 2318 | *< 0.0001* |
| *D. mel* | *2* | *15* | *D-L* | *-60.85* | *2.98* | *-20.44* | 2318 | *< 0.0001* |
| *D. sim* | *1* | *15* | *C-D* | *34.66* | *2.95* | *11.75* | 2318 | *< 0.0001* |
| *D. sim* | *1* | *15* | *C-L* | *-28.37* | *2.91* | *-9.76* | 2318 | *< 0.0001* |
| *D. sim* | *1* | *15* | *D-L* | *-63.03* | *2.94* | *-21.44* | 2318 | *< 0.0001* |
| D. sim | 2 | 15 | C-D | 8.29 | 2.89 | 2.87 | 2318 | 0.58 |
| *D. sim* | *2* | *15* | *C-L* | *-41.83* | *2.94* | *-14.21* | 2318 | *< 0.0001* |
| *D. sim* | *2* | *15* | *D-L* | *-50.12* | *2.93* | *-17.13* | 2318 | *< 0.0001* |
| **Females:** | | | | | | | | |
| Species | Pop | Gen | Contrast | estimate | SE | t-ratio | df | p-value |
| *D. mel* | *1* | *5* | *C-D* | *16.37* | *1.94* | *8.45* | *2561* | *< 0.0001* |
| D. mel | 1 | 5 | C-L | 3.82 | 1.93 | 1.98 | 2561 | 0.99 |
| *D. mel* | *1* | *5* | *D-L* | *-12.56* | *1.84* | *-6.83* | *2561* | *< 0.0001* |
| *D. mel* | *2* | *5* | *C-D* | *20.72* | *1.85* | *11.22* | *2561* | *< 0.0001* |
| *D. mel* | *2* | *5* | *C-L* | *7.39* | *1.85* | *4.00* | *2561* | *0.028* |
| *D. mel* | *2* | *5* | *D-L* | *-13.33* | *1.85* | *-7.21* | *2561* | *< 0.0001* |
| *D. sim* | *1* | *5* | *C-D* | *9.19* | *1.86* | *4.93* | *2561* | *0.0005* |
| D. sim | 1 | 5 | C-L | -3.51 | 1.87 | -1.88 | 2561 | 0.99 |
| *D. sim* | *1* | *5* | *D-L* | *-12.69* | *1.87* | *-6.78* | *2561* | *< 0.0001* |
| *D. sim* | *2* | *5* | *C-D* | *14.37* | *1.92* | *7.47* | *2561* | *< 0.0001* |
| D. sim | 2 | 5 | C-L | 0.71 | 1.95 | 0.36 | 2561 | 1.0 |
| *D. sim* | *2* | *5* | *D-L* | *-13.67* | *1.99* | *-6.86* | *2561* | *< 0.0001* |
| *D. mel* | *1* | *10* | *C-D* | *27.3* | *1.89* | *14.77* | *2561* | *< 0.0001* |
| D. mel | 1 | 10 | C-L | -3.70 | 1.86 | -1.99 | 2561 | 0.99 |
| *D. mel* | *1* | *10* | *D-L* | *-31.64* | *1.86* | *-17.02* | *2561* | *< 0.0001* |
| *D. mel* | *2* | *10* | *C-D* | *32.27* | *1.83* | *17.61* | *2561* | *< 0.0001* |
| D. mel | 2 | 10 | C-L | -0.133 | 1.84 | -0.072 | 2561 | 1.0 |
| *D. mel* | *2* | *10* | *D-L* | *-32.41* | *1.84* | *-17.61* | *2561* | *< 0.0001* |
| *D. sim* | *1* | *10* | *C-D* | *20.74* | *1.88* | *11.04* | *2561* | *< 0.0001* |
| *D. sim* | *1* | *10* | *C-L* | *-11.03* | *1.84* | *-5.98* | *2561* | *< 0.0001* |
| *D. sim* | *1* | *10* | *D-L* | *-31.77* | *1.87* | *-16.97* | *2561* | *< 0.0001* |
| *D. sim* | *2* | *10* | *C-D* | *25.93* | *1.85* | *13.98* | *2561* | *< 0.0001* |
| D. sim | 2 | 10 | C-L | -6.81 | 1.86 | -3.66 | 2561 | 0.090 |
| *D. sim* | *2* | *10* | *D-L* | *-32.75* | *1.88* | *-17.43* | *2561* | *< 0.0001* |
| *D. mel* | *1* | *15* | *C-D* | *28.09* | *2.62* | *10.72* | *2561* | *< 0.0001* |
| *D. mel* | *1* | *15* | *C-L* | *-18.95* | *2.63* | *-7.19* | *2561* | *< 0.0001* |
| *D. mel* | *1* | *15* | *D-L* | *-47.04* | *2.62* | *-17.95* | *2561* | *< 0.0001* |
| *D. mel* | *2* | *15* | *C-D* | *32.43* | *2.61* | *12.43* | *2561* | *< 0.0001* |
| *D. mel* | *2* | *15* | *C-L* | *-15.37* | *2.67* | *-5.77* | *2561* | *< 0.0001* |
| *D. mel* | *2* | *15* | *D-L* | *-47.81* | *2.66* | *-17.95* | *2561* | *< 0.0001* |
| *D. sim* | *1* | *15* | *C-D* | *20.90* | *2.62* | *7.99* | *2561* | *< 0.0001* |
| *D. sim* | *1* | *15* | *C-L* | *-26.27* | *2.63* | *-9.98* | *2561* | *< 0.0001* |
| *D. sim* | *1* | *15* | *D-L* | *-47.18* | *2.64* | *-17.89* | *2561* | *< 0.0001* |
| *D. sim* | *2* | *15* | *C-D* | *26.09* | *2.62* | *9.97* | *2561* | *< 0.0001* |
| *D. sim* | *2* | *15* | *C-L* | *-22.06* | *2.64* | *8.36* | *2561* | *< 0.0001* |
| *D. sim* | *2* | *15* | *D-L* | *-48.15* | *2.65* | *-18.15* | *2561* | *< 0.0001* |

Table S3: Results from generalized linear model on the color data at generation 15. Italics represent significant differences.

| **Males:** | | | | |
| --- | --- | --- | --- | --- |
| Effect | F Values | df-num | df-dem | p-value |
| *Treatment* | *460.42* | *1* | *345* | *<2.2e-16* |
| *Species* | *46.51* | *1* | *345* | *4.1e-11* |
| Population nested in Species | 1.66 | 2 | 345 | 0.19 |
| *Treatment x Species* | *55.56* | *1* | *345* | *7.4e-13* |
| *Treatment x Population nested in Species* | *12.43* | *2* | *345* | *6.1e-6* |
| **Females:** | | | | |
| *Treatment* | *221.99* | *1* | *337* | *< 2.2e-16* |
| *Species* | *13.79* | *1* | *337* | *0.0002* |
| *Population nested in Species* | *13.59* | *2* | *337* | *2.1e-6* |
| Treatment x Species | 1.12 | 1 | 337 | 0.29 |
| *Treatment x Population nested in Species* | *5.20* | *2* | *337* | *0.006* |

Table S4: Results of analysis of slope for each population for the color data at generation 15. Italics represent significant differences.

| **Males:** | | | | | | | | |
| --- | --- | --- | --- | --- | --- | --- | --- | --- |
| Effect | Slope | SE | | X^2^ Value | df | | p-value | |
| *D. mel – 1* | *-42.6* | *1.99* | | *460.0* | *1* | | *< 0.0001* | |
| *D. mel – 2* | *-29.0* | *1.95* | | *221.0* | *1* | | *< 0.0001* | |
| *D. sim - 1* | *-21.9* | *1.95* | | *125.0* | *1* | | *< 0.0001* | |
| *D. sim - 2* | *-24.5* | *1.95* | | *158.0* | *1* | | *< 0.0001* | |
| **Females:** | | | | | | | | |
| *D. mel – 1* | *-27.9* | | *1.87* | *222.0* | | *1* | | *< 0.0001* |
| *D. mel – 2* | *-19.4* | | *2.04* | *90.5* | | *1* | | *< 0.0001* |
| *D. sim - 1* | *-25.1* | | *1.90* | *173.4* | | *1* | | *< 0.0001* |
| *D. sim - 2* | *-22.3* | | *1.89* | *139.5* | | *1* | | *< 0.0001* |

Table S5: Results of pairwise t-tests of color data at generation 15. Italics represent significant differences.

| **Males:** | | | | | | | |
| --- | --- | --- | --- | --- | --- | --- | --- |
| Species | Population | Contrast | estimate | SE | t-ratio | df | p-value |
| *D. mel* | *1* | *C-D* | *50.3* | *3.90* | *12.88* | *341* | *< 0.0001* |
| *D. mel* | *1* | *C-L* | *-35.1* | *3.87* | *-9.08* | *341* | *< 0.0001* |
| *D. mel* | *1* | *D-L* | *-85.4* | *3.93* | *-21.70* | *341* | *< 0.0001* |
| *D. mel* | *2* | *C-D* | *23.0* | *3.87* | *5.95* | *341* | *< 0.0001* |
| *D. mel* | *2* | *C-L* | *-35.1* | *3.90* | *-9.0* | *341* | *< 0.0001* |
| *D. mel* | *2* | *D-L* | *-58.1* | *3.87* | *-15.02* | *341* | *< 0.0001* |
| *D. sim* | *1* | *C-D* | *20.9* | *3.87* | *5.40* | *341* | *< 0.0001* |
| *D. sim* | *1* | *C-L* | *-22.8* | *3.83* | *-5.96* | *341* | *< 0.0001* |
| *D. sim* | *1* | *D-L* | *-43.7* | *3.87* | *-11.31* | *341* | *< 0.0001* |
| *D. sim* | *2* | *C-D* | *19.2* | *3.83* | *5.01* | *341* | *0.0001* |
| *D. sim* | *2* | *C-L* | *-29.9* | *3.87* | *-7.73* | *341* | *< 0.0001* |
| *D. sim* | *2* | *D-L* | *-49.1* | *3.87* | *-12.70* | *341* | *< 0.0001* |
| **Females:** | | | | | | | |
| Species | Population | Contrast | estimate | SE | t-ratio | df | p-value |
| *D. mel* | *1* | *C-D* | *33.7* | *3.74* | *9.02* | *333* | *< 0.0001* |
| *D. mel* | *1* | *C-L* | *-22.1* | *3.74* | *-5.92* | *333* | *< 0.0001* |
| *D. mel* | *1* | *D-L* | *-55.8* | *3.74* | *-14.94* | *333* | *< 0.0001* |
| *D. mel* | *2* | *C-D* | *19.4* | *3.83* | *5.06* | *333* | *< 0.0001* |
| *D. mel* | *2* | *C-L* | *-19.5* | *4.12* | *-4.73* | *333* | *0.0002* |
| *D. mel* | *2* | *D-L* | *-38.9* | *4.09* | *-9.51* | *333* | *< 0.0001* |
| *D. sim* | *1* | *C-D* | *30.2* | *3.80* | *7.94* | *333* | *< 0.0001* |
| *D. sim* | *1* | *C-L* | *-20.0* | *3.80* | *-5.26* | *333* | *< 0.0001* |
| *D. sim* | *1* | *D-L* | *-50.2* | *3.80* | *-13.20* | *333* | *< 0.0001* |
| *D. sim* | *2* | *C-D* | *23.9* | *3.77* | *6.34* | *333* | *< 0.0001* |
| *D. sim* | *2* | *C-L* | *-20.8* | *3.74* | *-5.56* | *333* | *< 0.0001* |
| *D. sim* | *2* | *D-L* | *-44.6* | *3.77* | *-11.85* | *333* | *< 0.0001* |

Table S6: Table of mean ± sd for color, aggression, basal activity at gen 12, geotaxis, total activity at gen 16, and sleep for each selection treatment per population and species at the end of the experiment.

| **Males:** | | | | | | |  |
| --- | --- | --- | --- | --- | --- | --- | --- |
| Pop | Trt | color | aggression | log(basal act) | log(geotaxis) | Total act | Sleep |
| D. mel-1 | D | 20.82 ± 7.82 | 0.50 ± 1.11 | 3.59 ± 1.40 | 1.61 ± 0.68 | 903.05 ± 128.57 | 848.67 ± 233.59 |
| D. mel-1 | C | 71.07 ± 15.68 | 0.07 ± 0.25 | 2.74 ± 1.59 | 2.52 ± 1.08 | 843.68 ± 117.97 | 818.44 ± 192.28 |
| D. mel-1 | L | 106.2 ± 17.74 | 0.03 ± 0.19 | 2.30 ± 1.85 | 1.92 ± 0.67 | 681.87 ± 157.74 | 872.20 ± 178.28 |
| D. mel-2 | D | 36.47 ± 14.19 | 0.40 ± 0.62 | 4.02 ± 0.90 | 2.11 ± 0.73 | 893.53 ± 157.92 | 665.16 ± 187.52 |
| D. mel-2 | C | 59.46 ± 12.28 | 0.03 ± 0.19 | 3.22 ± 1.36 | 2.10 ± 0.81 | 706.14 ± 114.74 | 923.86 ± 204.87 |
| D. mel-2 | L | 94.54 ± 19.53 | 0.03 ± 0.19 | 2.82 ± 1.40 | 2.72 ± 1.13 | 675.30 ± 118.04 | 899.89 ± 170.62 |
| D. sim-1 | D | 29.15 ± 10.44 | 0.14 ± 0.35 | 1.76 ± 1.55 | 2.21 ± 0.64 | 362.27 ± 135.49 | 1065.50 ± 168.06 |
| D. sim-1 | C | 50.04 ± 8.73 | 0.0 ± 0.0 | 1.09 ± 1.32 | 2.70 ± 0.78 | 317.85 ± 168.42 | 1117.64 ± 162.96 |
| D. sim-1 | L | 72.88 ± 14.82 | 0.03 ± 0.18 | 0.92 ± 1.24 | 2.64 ± 0.80 | 567.53 ± 167.62 | 914.11 ± 203.51 |
| D. sim-2 | D | 24.76 ± 15.81 | 0.43 ± 0.63 | 2.17 ± 1.45 | 2.49 ± 1.50 | 379.73 ± 177.89 | 1085.18 ± 150.03 |
| D. sim-2 | C | 43.96 ± 19.11 | 0.07 ± 0.25 | 1.13 ± 1.36 | 2.66 ± 1.50 | 371.18 ± 217.15 | 1122.04 ± 135.54 |
| D. sim-2 | L | 73.86 ± 16.25 | 0.03 ± 0.19 | 0.80 ± 1.07 | 2.97 ± 1.49 | 417.85 ± 176.28 | 1026.59 ± 155.58 |
| **Females:** | | | | | | |  |
| Pop | Trt | color | aggression | log(basal act) | log(geotaxis) | Total act | Sleep |
| D. mel-1 | D | 16.83 ± 8.42 | 0.20 ± 0.41 | 3.91 ± 1.0 | 2.13 ± 0.58 | 646.34 ± 330.77 | 793.91 ± 206.77 |
| D. mel-1 | C | 50.54 ± 18.26 | 0.03 ± 0.18 | 3.47 ± 0.97 | 2.63 ± 0.90 | 638.20 ± 430.38 | 851.51 ± 150.92 |
| D. mel-1 | L | 72.64 ± 13.00 | 0.03 ± 0.18 | 3.37 ± 1.05 | 2.73 ± 1.22 | 552.79 ± 258.27 | 812.91 ± 207.16 |
| D. mel-2 | D | 25.17 ± 8.86 | 0.17 ± 0.38 | 2.99 ± 1.52 | 2.89 ± 0.79 | 519.21 ± 222.42 | 796.67 ± 217.83 |
| D. mel-2 | C | 44.55 ± 22.70 | 0.04 ± 0.19 | 2.91 ± 1.62 | 2.27 ± 0.43 | 478.90 ± 232.38 | 915.66 ± 176.25 |
| D. mel-2 | L | 64.05 ± 21.02 | 0.05 ± 0.21 | 2.23 ± 1.77 | 2.30 ± 0.53 | 610.00 ± 293.17 | 789.17 ± 189.73 |
| D. sim-1 | D | 27.99 ± 8.57 | 0.10 ± 0.31 | 1.51 ± 1.51 | 2.71 ± 0.68 | 257.96 ± 145.63 | 1036.78 ± 221.96 |
| D. sim-1 | C | 58.16 ± 10.20 | 0.07 ± 0.26 | 1.15 ± 1.43 | 2.39 ± 0.77 | 293.95 ± 126.79 | 993.99 ± 198.30 |
| D. sim-1 | L | 78.15 ± 13.29 | 0.0 ± 0.0 | 1.04 ± 1.31 | 3.08 ± 1.10 | 325.44 ± 205.15 | 953.76 ± 230.44 |
| D. sim-2 | D | 20.70 ± 16.86 | 0.24 ± 0.44 | 1.39 ± 1.50 | 2.71 ± 1.52 | 476.80 ± 216.05 | 844.64 ± 173.48 |
| D. sim-2 | C | 44.58 ± 8.93 | 0.03 ± 0.18 | 1.31 ± 1.34 | 2.35 ± 1.58 | 403.42 ± 206.53 | 961.48 ± 175.07 |
| D. sim-2 | L | 65.35 ± 16.07 | 0.03 ± 0.28 | 1.13 ± 1.17 | 2.79 ± 1.03 | 426.72 ± 163.93 | 842.16 ± 168.50 |

Table S7: Results from generalized linear model of aggression evolution over the course of the experiment. Italics represent significant differences.

| **Males:** | | | |
| --- | --- | --- | --- |
| Effect | X^2^ Values | df | p-value |
| Treatment | 0.73 | 1 | 0.39 |
| Species | 2.03 | 1 | 0.15 |
| *Generation* | *12.07* | *1* | *0.0003* |
| *Population nested in Species* | 34.59 | 2 | 3.1e-8 |
| Treatment x Species | 0.925 | 1 | 0.34 |
| *Treatment x Generation* | *4.62* | *1* | *0.031* |
| Species x Generation | 0.054 | 1 | 0.82 |
| *Treatment x Population nested in Species* | *6.75* | *2* | *0.034* |
| *Generation x Population nested in Species* | *25.28* | *2* | *3.2e-6* |
| **Females:** | | | |
| *Treatment* | *8.90* | *1* | *0.0028* |
| *Species* | *16.42* | *1* | *5.1e-5* |
| *Generation* | *5.46* | *1* | *0.019* |
| *Population nested in Species* | *17.40* | *2* | *0.0002* |
| Treatment x Species | 1.20 | 1 | 0.28 |
| Treatment x Generation | 0.078 | 1 | 0.78 |
| *Species x Generation* | *3.26* | *1* | *0.071* |
| Treatment x Population nested in Species | 0.52 | 2 | 0.77 |
| *Generation x Population nested in Species* | *8.37* | *2* | *0.015* |

Table S8: Results of asymptotic z-tests of aggression data over the course of the experiment. Estimates are on the log scale. Italics represent significant differences.

| **Males:** | | | | | | | | |
| --- | --- | --- | --- | --- | --- | --- | --- | --- |
| Species | Pop | Gen | Contrast | estimate | SE | z-ratio | df | p-value |
| D. mel | 1 | 5 | C-D | -0.024 | 0.22 | -0.11 | Inf | 1.0 |
| D. mel | 1 | 5 | C-L | 1.21 | 0.33 | 2.68 | Inf | 0.085 |
| D. mel | 1 | 5 | D-L | 1.23 | 0.32 | 3.83 | Inf | 0.051 |
| D. mel | 2 | 5 | C-D | -1.09 | 0.39 | -2.78 | Inf | 0.65 |
| D. mel | 2 | 5 | C-L | 1.21 | 0.72 | 1.68 | Inf | 1.0 |
| D. mel | 2 | 5 | D-L | 2.30 | 0.66 | 3.47 | Inf | 0.16 |
| D. sim | 1 | 5 | C-D | 0.34 | 0.33 | 1.03 | Inf | 1.0 |
| D. sim | 1 | 5 | C-L | 1.12 | 0.42 | 2.64 | Inf | 0.76 |
| D. sim | 1 | 5 | D-L | 0.78 | 0.43 | 1.81 | Inf | 1.0 |
| D. sim | 2 | 5 | C-D | 0.38 | 0.43 | 0.88 | Inf | 1.0 |
| D. sim | 2 | 5 | C-L | 1,07 | 0.62 | 1.74 | Inf | 1.0 |
| D. sim | 2 | 5 | D-L | 0.69 | 0.58 | 1.19 | Inf | 1.0 |
| D. mel | 1 | 10 | C-D | -0.72 | 0.23 | -3.11 | Inf | 0.38 |
| D. mel | 1 | 10 | C-L | 0.83 | 0.34 | 2.44 | Inf | 0.89 |
| *D. mel* | *1* | *10* | *D-L* | *1.55* | *0.32* | *4.92* | *Inf* | *0.0005* |
| *D. mel* | *2* | *10* | *C-D* | *-1.79* | *0.38* | *-4.66* | *Inf* | *0.0017* |
| D. mel | 2 | 10 | C-L | 0.83 | 0.67 | 1.23 | Inf | 1.0 |
| *D. mel* | *2* | *10* | *D-L* | *2.62* | *0.60* | *4.38* | *Inf* | *0.0062* |
| D. sim | 1 | 10 | C-D | -0.35 | *0.35* | -1.01 | Inf | 1.0 |
| D. sim | 1 | 10 | C-L | 0.74 | 0.45 | 1.66 | Inf | 1.0 |
| D. sim | 1 | 10 | D-L | 1.09 | *0.44* | 2.51 | Inf | 0.85 |
| D. sim | 2 | 10 | C-D | -0.32 | 0.34 | -0.93 | Inf | 1.0 |
| D. sim | 2 | 10 | C-L | 0.69 | 0.47 | 1.47 | Inf | 1.0 |
| D. sim | 2 | 10 | D-L | 1.01 | 0.44 | 2.30 | Inf | 0.94 |
| D. mel | 1 | 15 | C-D | -1.42 | 0.39 | -3.59 | Inf | 0.11 |
| D. mel | 1 | 15 | C-L | 0.45 | 0.58 | 0.78 | Inf | 1.0 |
| D. mel | 1 | 15 | D-L | 1.87 | 0.53 | 3.55 | Inf | 0.13 |
| *D. mel* | *2* | *15* | *C-D* | *-2.48* | *0.49* | *-5.09* | *Inf* | *0.0002* |
| D. mel | 2 | 15 | C-L | 0.45 | 0.78 | 0.58 | Inf | 1.0 |
| *D. mel* | *2* | *15* | *D-L* | *2.93* | *0.68* | *4.34* | *Inf* | *0.0073* |
| D. sim | 1 | 15 | C-D | -1.05 | 0.48 | -2.18 | Inf | 0.97 |
| D. sim | 1 | 15 | C-L | 0.36 | 0.66 | 0.55 | Inf | 1.0 |
| D. sim | 1 | 15 | D-L | 1.41 | 0.62 | 2.29 | Inf | 0.94 |
| D. sim | 2 | 15 | C-D | -1.01 | 0.39 | -2.62 | Inf | 0.78 |
| D. sim | 2 | 15 | C-L | 0.32 | 0.53 | 0.60 | Inf | 1.0 |
| D. sim | 2 | 15 | D-L | 1.33 | 0.48 | 2.78 | Inf | 0.66 |
| **Females:** | | | | | | | | |
| Species | Pop | Gen | Contrast | estimate | SE | z-ratio | df | p-value |
| D. mel | 1 | 5 | C-D | -0.83 | 0.31 | -2.72 | Inf | 0.71 |
| D. mel | 1 | 5 | C-L | 0.78 | 0.44 | 1.78 | Inf | 1.0 |
| *D. mel* | *1* | *5* | *D-L* | *1.60* | *0.38* | *4.21* | *Inf* | *0.012* |
| D. mel | 2 | 5 | C-D | -0.43 | 0.43 | -0.99 | Inf | 1.0 |
| D. mel | 2 | 5 | C-L | 1.73 | 0.83 | 2.08 | Inf | 0.98 |
| D. mel | 2 | 5 | D-L | 2.16 | 0.80 | 2.69 | Inf | 0.73 |
| D. sim | 1 | 5 | C-D | -1.53 | 0.60 | -2.54 | Inf | 0.83 |
| D. sim | 1 | 5 | C-L | 0.60 | 0.93 | 0.65 | Inf | 1.0 |
| D. sim | 1 | 5 | D-L | 2.13 | 0.81 | 2.63 | Inf | 0.77 |
| D. sim | 2 | 5 | C-D | -1.29 | 0.56 | -2.31 | Inf | 0.94 |
| D. sim | 2 | 5 | C-L | 0.21 | 0.80 | 0.27 | Inf | 1.0 |
| D. sim | 2 | 5 | D-L | 1.50 | 0.70 | 2.14 | Inf | 0.98 |
| D. mel | 1 | 10 | C-D | -0.70 | 0.28 | -2.48 | Inf | 0.87 |
| D. mel | 1 | 10 | C-L | 0.90 | 0.42 | 2.12 | Inf | 0.98 |
| *D. mel* | *1* | *10* | *D-L* | *1.60* | *0.39* | *4.12* | *Inf* | *0.017* |
| D. mel | 2 | 10 | C-D | -0.30 | 0.37 | -0.83 | Inf | 1.0 |
| D. mel | 2 | 10 | C-L | 1.85 | 0.77 | 2.419 | Inf | 0.90 |
| D. mel | 2 | 10 | D-L | 2.15 | 0.75 | 2.87 | Inf | 0.58 |
| D. sim | 1 | 10 | C-D | -1.40 | 0.56 | -2.50 | Inf | 0.86 |
| D. sim | 1 | 10 | C-L | 0.72 | 0.87 | 0.83 | Inf | 1.0 |
| D. sim | 1 | 10 | D-L | 2.12 | 0.75 | 2.81 | Inf | 0.63 |
| D. sim | 2 | 10 | C-D | -1.17 | 0.47 | -2.47 | Inf | 0.87 |
| D. sim | 2 | 10 | C-L | 0.33 | 0.65 | 0.51 | Inf | 1.0 |
| D. sim | 2 | 10 | D-L | 1.50 | 0.56 | 2.70 | Inf | 0.72 |
| D. mel | 1 | 15 | C-D | -0.58 | 0.46 | -1.25 | Inf | 1.0 |
| D. mel | 1 | 15 | C-L | 1.01 | 0.72 | 1.40 | Inf | 1.0 |
| D. mel | 1 | 15 | D-L | 1.59 | 0.67 | 2.36 | Inf | 0.92 |
| D. mel | 2 | 15 | C-D | -0.18 | 0.48 | -0.37 | Inf | 1.0 |
| D. mel | 2 | 15 | C-L | 1.97 | 0.91 | 2.15 | Inf | 0.98 |
| D. mel | 2 | 15 | D-L | 2.15 | 0.88 | 2.44 | Inf | 0.89 |
| D. sim | 1 | 15 | C-D | -1.28 | 0.64 | -1.98 | Inf | 0.99 |
| D. sim | 1 | 15 | C-L | 0.84 | 1.0 | 0.84 | Inf | 1.0 |
| D. sim | 1 | 15 | D-L | 2.11 | 0.88 | 2.39 | Inf | 0.91 |
| D. sim | 2 | 15 | C-D | -1.04 | 0.53 | -1.97 | Inf | 0.99 |
| D. sim | 2 | 15 | C-L | 0.45 | 0.74 | 0.61 | Inf | 1.0 |
| D. sim | 2 | 15 | D-L | 1.49 | 0.65 | 2.30 | Inf | 0.94 |

Table S9: Results from generalized linear model with Poisson distribution of aggression data at generation 15. Italics represent significant differences.

| **Males:** | | | |
| --- | --- | --- | --- |
| Effect | X^2^ Value | df | p-value |
| *Treatment* | *50.30* | *1* | *1.3e-12* |
| *Species* | *7.27* | *1* | *0.0070* |
| *Population nested in Species* | *6.237* | *2* | *0.044* |
| **Females:** | | | |
| *Treatment* | *17.47* | *1* | *2.9e-5* |
| Species | 1.11 | 1 | 0.44 |
| Population nested in Species | 21.31 | 2 | 0.57 |

Table S10: Results of analysis of slope for each sex for the aggression data at generation 15. Italics represent significant differences.

| **Males:** | | | | | |
| --- | --- | --- | --- | --- | --- |
| Effect | Slope | SE | X^2^ Value | df | p-value |
| *Treatment* | *1.52* | *0.37* | *31.5* | *1* | *< 0.0001* |
| **Females:** | | | | | |
| Effect | Slope | SE | X^2^ Value | df | p-value |
| *Treatment* | *1.1* | *0.3* | *13.3* | *1* | *0.0003* |

Table S11: Results of asymptotic z-tests of aggression data at generation 15. Estimates are on the log scale. Italics represent significant differences.

| **Males:** | | | | | |
| --- | --- | --- | --- | --- | --- |
| Contrast | estimate | SE | z-ratio | df | p-value |
| *C – D* | *-2.17* | *0.47* | *-4.59* | *Inf* | *< 0.0001* |
| C – L | 0.20 | 0.67 | 0.30 | Inf | 0.95 |
| *D – L* | *2.37* | *0.53* | *4.53* | *Inf* | *< 0.0001* |
| **Females:** | | | | | |
| Contrast | estimate | SE | z-ratio | df | p-value |
| *C – D* | *-1.44* | *0.50* | *-2.89* | *Inf* | *0.011* |
| C – L | 0.46 | 0.73 | 0.63 | Inf | 0.81 |
| *D – L* | *1.90* | *0.62* | *3.07* | *Inf* | *0.0061* |

Table S12: Results from generalized linear models with negative binomial distribution of basal activity data at generation 12. Italics represent significant differences.

| **Males:** | | | |
| --- | --- | --- | --- |
| Effect | X^2^ Value | df | p-value |
| *Treatment* | *71.07* | *1* | *< 2.2e-16* |
| *Species* | *35.62* | *1* | *2.4e-9* |
| Population nested in Species | 1.38 | 2 | 0.50 |
| **Females:** | | | |
| *Treatment* | *109.77* | *1* | *< 2.2e-16* |
| *Species* | *11.73* | *1* | *0.0006* |
| Population nested in Species | 3.97 | 2 | 0.14 |

Table S13: Results of analysis of slope for each population for the basal activity data at generation 12. Italics represent significant differences.

| **Males:** | | | | | |
| --- | --- | --- | --- | --- | --- |
| Effect | Slope | SE | X^2^ Value | df | p-value |
| *Treatment* | *0.51* | *0.087* | *34.0* | *1* | *< 0.0001* |
| **Females:** | | | | | |
| Effect | Slope | SE | X^2^ Value | df | p-value |
| *Treatment* | *0.27* | *0.080* | *11.7* | *1* | *0.0006* |

Table S14: Results of asymptotic z-tests of basal activity data at generation 12. Estimates are on the log scale. Italics represent significant differences.

| **Males:** | | | | | | | | | |
| --- | --- | --- | --- | --- | --- | --- | --- | --- | --- |
| Contrast | Estimate | SE | | z-ratio | | df | | | p-value |
| *C – D* | *-0.71* | *0.16* | | *-4.44* | | *Inf* | | | *< 0.001* |
| C – L | 0.31 | 0.16 | | 1.94 | | Inf | | | 0.13 |
| *D – L* | *1.02* | *0.17* | | *5.86* | | *Inf* | | | *< 0.001* |
| **Females:** | | | | | | | | | |
| Contrast | estimate | | SE | | z-ratio | | df | p-value | |
| C – D | -0.29 | | 0.14 | | -1.99 | | Inf | 0.11 | |
| C – L | 0.261 | | 0.14 | | 1.81 | | Inf | 0.17 | |
| *D – L* | *0.547* | | *0.16* | | *3.43* | | *Inf* | *0.002* | |

Table S15: Results from survival analysis for geotaxis data at generation 13. Italics represent significant differences.

| **Males:** | | | |
| --- | --- | --- | --- |
| Effect | Chi-square | Degrees of Freedom | p-value |
| Treatment | 3.75 | 1 | 0.053 |
| Species | 0.213 | 1 | 0.64 |
| *Population nested in Species* | *42.29* | *2* | *6.6e-10* |
| *Treatment x Species* | *10.02* | 1 | *0.002* |
| *Treatment x Population nested in Species* | *23.53* | *2* | *7.8e-6* |
| **Females:** | | | |
| *Treatment* | *10.11* | *1* | *0.0015* |
| *Species* | *13.77* | *1* | *0.0002* |
| *Population nested in Species* | *33.32* | *2* | *5.8e-8* |
| Treatment x Species | 1.66 | 1 | 0.20 |
| *Treatment x Population nested in Species* | *36.64* | *2* | *1.1e-8* |

Table S16: Results of analysis of slope for each population for the geotaxis data at generation 13. Italics represent significant differences.

| **Males:** | | | | | | | | |
| --- | --- | --- | --- | --- | --- | --- | --- | --- |
| Effect | Slope | SE | | X^2^ Value | df | | p-value | |
| D. mel – 1 | -0.26 | 0.14 | | 3.75 | 1 | | 0.053 | |
| *D. mel – 2* | *0.47* | *0.12* | | *15.28* | *1* | | *0.0001* | |
| *D. sim - 1* | *0.32* | *0.12* | | *7.74* | *1* | | *0.0054* | |
| D. sim - 2 | -0.16 | 0.13 | | 1.48 | 1 | | 0.22 | |
| **Females:** | | | | | | | | |
| *D. mel – 1* | *0.36* | | *0.11* | *10.11* | | *1* | | *0.0015* |
| *D. mel – 2* | *-0.43* | | *0.11* | *15.51* | | *1* | | *0.0001* |
| D. sim - 1 | 0.15 | | 0.11 | 1.78 | | 1 | | 0.18 |
| *D. sim - 2* | *-0.33* | | *0.14* | *5.45* | | *1* | | *0.020* |

Table S17: Results of asymptotic z-tests correction of geotaxis data at generation 13. Estimates are on the log scale. Italics represent significant differences.

| **Males:** | | | | | | | |
| --- | --- | --- | --- | --- | --- | --- | --- |
| Species | Population | Contrast | estimate | SE | z-ratio | df | p-value |
| *D. mel* | *1* | *C-D* | *-0.71* | *0.19* | *-3.66* | *Inf* | *0.0135* |
| *D. mel* | *1* | *C-L* | *-1.11* | *0.19* | *-6.00* | *Inf* | *< 0.001* |
| D. mel | 1 | D-L | -0.40 | 0.21 | -1.90 | Inf | 0.76 |
| D. mel | 2 | C-D | -0.43 | 0.20 | -2.16 | Inf | 0.58 |
| D. mel | 2 | C-L | 0.44 | 0.20 | 2.21 | Inf | 0.54 |
| *D. mel* | *2* | *D-L* | *0.87* | *0.23* | *3.80* | *Inf* | *0.0080* |
| D. sim | 1 | C-D | -0.54 | 0.20 | -2.73 | Inf | 0.21 |
| D. sim | 1 | C-L | 0.080 | 0.20 | 0.40 | Inf | 1.0 |
| D. sim | 1 | D-L | 0.61 | 0.22 | 2.86 | Inf | 0.16 |
| D. sim | 2 | C-D | 0.24 | 0.26 | 0.91 | Inf | 1.0 |
| D. sim | 2 | C-L | -0.06 | 0.24 | -0.23 | Inf | 1.0 |
| D. sim | 2 | D-L | -0.29 | 0.26 | -1.14 | Inf | 1.0 |
| **Females:** | | | | | | | |
| Species | Population | Contrast | estimate | SE | z-ratio | df | p-value |
| D. mel | 1 | C-D | -0.19 | 0.23 | -0.84 | Inf | 1.0 |
| D. mel | 1 | C-L | 0.45 | 0.19 | 2.32 | Inf | 0.46 |
| D. mel | 1 | D-L | 0.64 | 0.24 | 2.67 | Inf | 0.24 |
| *D. mel* | *2* | *C-D* | *0.83* | *0.22* | *3.81* | *Inf* | *0.0076* |
| D. mel | 2 | C-L | -0.088 | 0.21 | -0.42 | Inf | 1.0 |
| *D. mel* | *2* | *D-L* | *-0.92* | *0.23* | *-3.93* | *Inf* | *0.0048* |
| D. sim | 1 | C-D | -0.040 | 0.20 | -0.21 | Inf | 1.0 |
| D. sim | 1 | C-L | 0.34 | 0.20 | 1.71 | Inf | 0.86 |
| D. sim | 1 | D-L | 0.38 | 0.23 | 1.69 | Inf | 0.87 |
| D. sim | 2 | C-D | -0.10 | 0.26 | -0.41 | Inf | 1.0 |
| D. sim | 2 | C-L | -0.82 | 0.27 | -3.01 | Inf | 0.11 |
| D. sim | 2 | D-L | -0.72 | 0.27 | -2.67 | Inf | 0.24 |

Table S18: Results from generalized linear model on the total activity data at generation 16. Italics represent significant differences.

| **Males:** | | | | |
| --- | --- | --- | --- | --- |
| Effect | F Values | df-num | df-dem | p-value |
| *Treatment* | *17.61* | *1* | *699* | *3.06e-5* |
| *Species* | *70.18* | *1* | *699* | *2.96e-16* |
| Population nested in Species | 0.78 | 2 | 699 | 0.46 |
| *Treatment x Species* | *17.70* | *1* | *699* | *2.92e-5* |
| **Females:** | | | | |
| *Treatment* | *4.28* | *1* | *719* | *0.039* |
| *Species* | *125.66* | *1* | *719* | *<2.2e-16* |
| *Population nested in Species* | *16.76* | *2* | *719* | *7.66e-8* |
| *Treatment x Species* | *5.26* | *1* | *719* | *0.022* |
| *Treatment x Population nested in Species* | *6.01* | *2* | *719* | *0.0023* |

Table S19: Results of analysis of slope for each population for the total activity data at generation 16. Italics represent significant differences.

| **Males:** | | | | | |
| --- | --- | --- | --- | --- | --- |
| Effect | Slope | SE | X^2^ Value | df | p-value |
| *D. mel* | *108.9* | *26.0* | *17.61* | *1* | *<0.0001* |
| D. sim | -52.7 | 28.3 | 3.46 | 1 | 0.063 |
| **Females:** | | | | | |
| Effect | Slope | SE | X^2^ Value | df | p-value |
| *D. mel-1* | *46.8* | *22.6* | *4.28* | *1* | *0.039* |
| *D. mel-2* | *-48.0* | *21.5* | *4.97* | *1* | *0.026* |
| D. sim-1 | -33.7 | 26.8 | 1.58 | 1 | 0.21 |
| D. sim-2 | 25.0 | 22.6 | 1.23 | 1 | 0.27 |

Table S20: Results of pairwise t-tests of total activity data at generation 16. Italics represent significant differences.

| **Males:** | | | | | | | |
| --- | --- | --- | --- | --- | --- | --- | --- |
| Species | Population | Contrast | estimate | SE | t-ratio | df | p-value |
| D. mel | 1 and 2 | C-D | -123.5 | 53.4 | -2.31 | 697 | 0.063 |
| D. mel | 1 and 2 | C-L | 94.8 | 52.2 | 1.82 | 697 | 0.21 |
| *D. mel* | *1 and 2* | *D-L* | *218.3* | *51.9* | *4.21* | *697* | *0.0001* |
| D. sim | 1 and 2 | C-D | -23.9 | 56.4 | -0.43 | 697 | 1.0 |
| D. sim | 1 and 2 | C-L | -130.6 | 57.0 | -2.29 | 697 | 0.067 |
| D. sim | 1 and 2 | D-L | -106.7 | 56.6 | -1.88 | 697 | 0.18 |
| **Females:** | | | | | | | |
| Species | Population | Contrast | estimate | SE | t-ratio | df | p-value |
| D. mel | 1 | C-D | -8.14 | 45.3 | -0.180 | 715 | 1.0 |
| D. mel | 1 | C-L | 85.4 | 45.3 | 1.885 | 715 | 0.18 |
| D. mel | 1 | D-L | 93.6 | 45.1 | 2.073 | 715 | 0.12 |
| D. mel | 2 | C-D | -40.3 | 45.3 | -0.89 | 715 | 1.0 |
| *D. mel* | *2* | *C-L* | *-131.1* | *43.2* | *-3.032* | *715* | *0.0076* |
| D. mel | 2 | D-L | -90.8 | 43.1 | -2.109 | 715 | 0.11 |
| D. sim | 1 | C-D | 36.0 | 53.3 | 0.676 | 715 | 1.0 |
| D. sim | 1 | C-L | -31.5 | 52.4 | -0.601 | 715 | 1.0 |
| D. sim | 1 | D-L | -67.5 | 53.5 | -1.260 | 715 | 0.62 |
| D. sim | 2 | C-D | -73.4 | 45.1 | -1.626 | 715 | 0.31 |
| D. sim | 2 | C-L | -23.3 | 45.1 | -0.516 | 715 | 1.0 |
| D. sim | 2 | D-L | 50.1 | 45.1 | 1.110 | 715 | 0.80 |

Table S21: Results from generalized linear model on the total sleep data at generation 16. Italics represent significant differences.

| **Males:** | | | | |
| --- | --- | --- | --- | --- |
| Effect | F Values | df-num | df-dem | p-value |
| Treatment | 0.51 | 1 | 697 | 0.48 |
| *Species* | *78.84* | *1* | *697* | *<2.2e-16* |
| Population nested in Species | 2.86 | 2 | 697 | 0.058 |
| *Treatment x Species* | *11.05* | *1* | *697* | *0.0009* |
| *Treatment x Population nested in Species* | *11.22* | *2* | *697* | *1.59e-5* |
| **Females:** | | | | |
| Treatment | 0.0895 | 1 | 722 | 0.76 |
| *Species* | *63.12* | *1* | *722* | *7.5e-15* |
| *Population nested in Species* | *13.01* | *2* | *722* | *2.83e-6* |

Table S22: Results of analysis of slope for each population for the total sleep data at generation 16. Italics represent significant differences.

| **Males:** | | | | | |
| --- | --- | --- | --- | --- | --- |
| Effect | Slope | SE | X^2^ Value | df | p-value |
| D. mel-1 | -12.0 | 16.8 | 0.51 | 1 | 0.48 |
| *D. mel-2* | *-113.9* | *15.9* | *51.4* | *1* | *<0.0001* |
| *D. sim-1* | *73.8* | *19.6* | *14.2* | *1* | *0.0002* |
| D. sim-2 | 29.1 | 16.5 | 3.13 | 1 | 0.077 |
| **Females:** | | | | | |
| Effect | Slope | SE | X^2^ Value | df | p-value |
| Treatment | 2.67 | 8.92 | 0.090 | 1 | 0.76 |

Table S23: Results of pairwise t-tests of total sleep data at generation 16. Italics represent significant differences.

| **Males:** | | | | | | | |
| --- | --- | --- | --- | --- | --- | --- | --- |
| Species | Population | Contrast | estimate | SE | t-ratio | df | p-value |
| D. mel | 1 | C-D | -30.2 | 32.7 | -0.92 | 693 | 1.0 |
| D. mel | 1 | C-L | -53.8 | 32.4 | -1.66 | 693 | 0.29 |
| D. mel | 1 | D-L | -23.5 | 32.6 | 0.72 | 693 | 1.0 |
| *D. mel* | *2* | *C-D* | *258.7* | *32.5* | *7.97* | *693* | *<0.0001* |
| D. mel | 2 | C-L | 24.0 | 31.3 | 0.767 | 693 | 1.0 |
| *D. mel* | *2* | *D-L* | *-234.7* | *30.8* | *-7.61* | *693* | *<0.0001* |
| D. sim | 1 | C-D | 52.1 | 37.3 | 1.40 | 693 | *0.49* |
| *D. sim* | *1* | *C-L* | *203.5* | *38.2* | *5.33* | *693* | *<0.0001* |
| *D. sim* | *1* | *D-L* | *151.4* | *38.0* | *4.0* | *693* | *0.0002* |
| D. sim | 2 | C-D | 36.9 | 32.1 | 1.15 | 693 | 0.75 |
| *D. sim* | *2* | *C-L* | *95.5* | *32.2* | *3.0* | *693* | *0.0094* |
| D. sim | 2 | D-L | 58.6 | 31.9 | 1.84 | 693 | 0.20 |

| **Females:** | | | | | |
| --- | --- | --- | --- | --- | --- |
| Contrast | estimate | SE | t-ratio | df | p-value |
| *C – D* | *60.47* | *17.9* | *3.38* | *721* | *0.0023* |
| *C – L* | *64.44* | *17.6* | *3.658* | *721* | *0.0008* |
| D – L | 3.97 | 17.7 | 0.225 | 721 | 0.002 |

Table S24: Spearman rank correlations between thorax color and behavior. Significant p-values (p < 0.05) are shown in italics.

| Behavior | Group | Species | Sex | Line | n | k | rho | 95% CI | p-value |
| --- | --- | --- | --- | --- | --- | --- | --- | --- | --- |
| Aggression | *Overall* | *Both* | *Both* | *All* | *7078* | *40* | *-0.035* | *(-0.059, -0.012)* | *0.003* |
|  | Species | *mel* | *Both* | *All* | *3531* | *20* | *-0.63* | *(-0.096, -0.03)* | *<0.001* |
|  |  | sim | Both | All | 3547 | 20 | -0.008 | (-0.041, 0.025) | 0.634 |
|  | Sex | *Both* | *Male* | *All* | *3416* | *20* | *-0.05* | *(-0.084, -0.016)* | *0.004* |
|  |  | Both | Female | All | 3662 | 20 | -0.022 | (-0.054, 0.011) | 0.194 |
|  | Line | Both | Both | C | 1649 | 12 | -0.043 | (-0.091, 0.006) | 0.086 |
|  |  | *Both* | *Both* | *D* | *1646* | *12* | *-0.082* | *(-0.13, -0.033)* | *0.001* |
|  |  | Both | Both | L | 1632 | 12 | -0.044 | (-0.093, 0.005) | 0.080 |
| Basal Activity | Overall | Both | Both | All | 1143 | 12 | -0.02 | (-0.079, 0.039) | 0.502 |
|  | Species | mel | Both | All | 612 | 6 | -0.036 | (-0.116, 0.044) | 0.38 |
|  |  | sim | Both | All | 531 | 6 | -0.002 | (-0.088, 0.085) | 0.967 |
|  | Sex | Both | Male | All | 537 | 6 | -0.004 | (-0.089, 0.082) | 0.933 |
|  |  | Both | Female | All | 606 | 6 | -0.035 | (-0.115, 0.046) | 0.398 |
|  | Line | Both | Both | C | 500 | 4 | -0.035 | (-0.124, 0.053) | 0.433 |
|  |  | Both | Both | D | 320 | 4 | -0.108 | (-0.217, 0.003) | 0.056 |
|  |  | Both | Both | L | 323 | 4 | 0.091 | (-0.02, 0.2) | 0.107 |
| Geotaxis | *Overall* | *Both* | *Both* | *All* | *1167* | *12* | *-0.084* | *(-0.142, -0.026)* | *0.005* |
|  | Species | mel | Both | All | 647 | 6 | -0.013 | (-0.065, 0.091) | 0.745 |
|  |  | *sim* | *Both* | *All* | *520* | *6* | *-0.204* | *(-0.286, -0.118)* | *<0.001* |
|  | Sex | *Both* | *Male* | *All* | *556* | *6* | *-0.173* | *(-0.254, -0.09)* | *<0.001* |
|  |  | Both | Female | All | 611 | 6 | -0.002 | (-0.083, 0.078) | 0.955 |
|  | Line | *Both* | *Both* | *C* | *517* | *4* | *-0.192* | *(-0.275, -0.107)* | *<0.001* |
|  |  | Both | Both | D | 318 | 4 | 0.047 | (-0.065, 0.158) | 0.408 |
|  |  | Both | Both | L | 332 | 4 | -0.036 | (-0.145, 0.073) | 0.518 |

Table S25: Results of pairwise Fisher z-tests comparing within-population color-aggression correlations between generation 0 and generation 15, broken down by line, species, and sex. Significant p-values (p < 0.05) are shown in italics.

| Line | Species | Sex | rho Gen 0 | rho Gen 15 | Z | p-value |
| --- | --- | --- | --- | --- | --- | --- |
| Control | Both | Both | 0.011 | -0.032 | 0.618 | 0.536 |
|  | mel | Both | -0.039 | -0.113 | 0.743 | 0.458 |
|  | sim | Both | 0.056 | 0.047 | 0.089 | 0.929 |
|  | Both | Males | -0.007 | -0.14 | 1.356 | 0.175 |
|  | Both | Females | 0.029 | 0.078 | -0.494 | 0.621 |
|  | mel | Males | -0.044 | -0.272 | 1.671 | 0.095 |
|  | mel | Females | -0.033 | 0.056 | -0.626 | 0.531 |
|  | sim | Males | 0.029 | -0.005 | 0.244 | 0.807 |
|  | sim | Females | 0.081 | 0.1 | -0.137 | 0.891 |
| Dark | *Both* | *Both* | *0.011* | *-0.168* | *2.564* | *0.01* |
|  | mel | Both | -0.039 | -0.133 | 0.948 | 0.343 |
|  | *sim* | *Both* | *0.056* | *-0.203* | *2.629* | *0.009* |
|  | Both | Males | -0.007 | -0.159 | 1.545 | 0.122 |
|  | *Both* | *Females* | *0.029* | *-0.176* | *2.08* | *0.037* |
|  | mel | Males | -0.044 | -0.244 | 1.446 | 0.148 |
|  | mel | Females | -0.033 | -0.02 | -0.092 | 0.927 |
|  | sim | Males | 0.029 | -0.074 | 0.735 | 0.462 |
|  | *sim* | *Females* | *0.081* | *-0.327* | *2.982* | *0.003* |
| Light | Both | Both | 0.011 | -0.079 | 1.262 | 0.207 |
|  | mel | Both | -0.039 | -0.011 | -0.265 | 0.791 |
|  | *sim* | *Both* | *0.056* | *-0.141* | *1.993* | *0.046* |
|  | Both | Males | -0.007 | -0.053 | 0.465 | 0.642 |
|  | Both | Females | 0.029 | -0.106 | 1.323 | 0.186 |
|  | mel | Males | -0.044 | -0.051 | 0.049 | 0.961 |
|  | mel | Females | -0.033 | 0.033 | -0.441 | 0.66 |
|  | sim | Males | 0.029 | -0.055 | 0.599 | 0.549 |
|  | *sim* | *Females* | *0.081* | *-0.224* | *2.209* | *0.027* |

Table S26: Group-level Spearman correlations between mean aggression (averaged across generations 5, 10, and 15) and mean basal activity across Line × Population × Sex combinations (n = 24 groups). rho² represents variance explained.

| Group | Aggression | rho | rho^2^ | p | n |
| --- | --- | --- | --- | --- | --- |
| Overall | Gen 15 | 0.265 | 0.07 | 0.197 | 24 |
|  | Average | 0.519 | 0.269 | 0.0044 | 24 |
| Control | Gen 15 | 0 | 0 | 1 | 8 |
|  | Average | -0.048 | 0.002 | 0.907 | 8 |
| Dark | Gen 15 | 0.429 | 0.184 | 0.2453 | 8 |
|  | Average | 0.857 | 0.734 | 0 | 8 |
| Light | Gen 15 | -0.155 | 0.024 | 0.7012 | 8 |
|  | Average | 0.179 | 0.032 | 0.6566 | 8 |
| *D. mel*-1 | Gen 15 | 0.457 | 0.209 | 0.304 | 6 |
|  | Average | 0.657 | 0.432 | 0.0812 | 6 |
| *D. mel*-2 | Gen 15 | 0.229 | 0.052 | 0.6387 | 6 |
|  | Average | 0.829 | 0.687 | 0.0031 | 6 |
| *D. sim*-1 | Gen 15 | 0.743 | 0.552 | 0.0265 | 6 |
|  | Average | 0.6 | 0.36 | 0.1336 | 6 |
| *D. sim*-2 | Gen 15 | 0.686 | 0.471 | 0.0595 | 6 |
|  | Average | 0.714 | 0.51 | 0.0412 | 6 |

Table S25: Results of binning photos at the start of the experiment. Italics indicate significant differences.

| Effect | F Values | df-num | df-dem | p-value |
| --- | --- | --- | --- | --- |
| *Bin* | *138.19* | *3* | *96* | *< 2.2e-16* |

Table S26: Results of post-hoc test of binning photos at the start of the experiment. Italics indicate significant differences.

| contrast | estimate | SE | t-ratio | df | p-value |
| --- | --- | --- | --- | --- | --- |
| *Bin 0 – Bin 1* | *19.1* | *4.17* | *4.59* | *96* | *0.0001* |
| *Bin 0 – Bin 2* | *38.5* | *2.92* | *13.19* | *96* | *< 0.0001* |
| *Bin 0 – Bin 3* | *56.1* | *2.84* | *19.77* | *96* | *< 0.0001* |
| *Bin 1 – Bin 2* | *19.4* | *3.98* | *4.87* | *96* | *< 0.0001* |
| *Bin 1 – Bin 3* | *37.0* | *3.92* | *9.44* | *96* | *< 0.0001* |
| *Bin 2 – Bin 3* | *17.6* | *2.55* | *6.92* | *96* | *< 0.0001* |

**Figures**

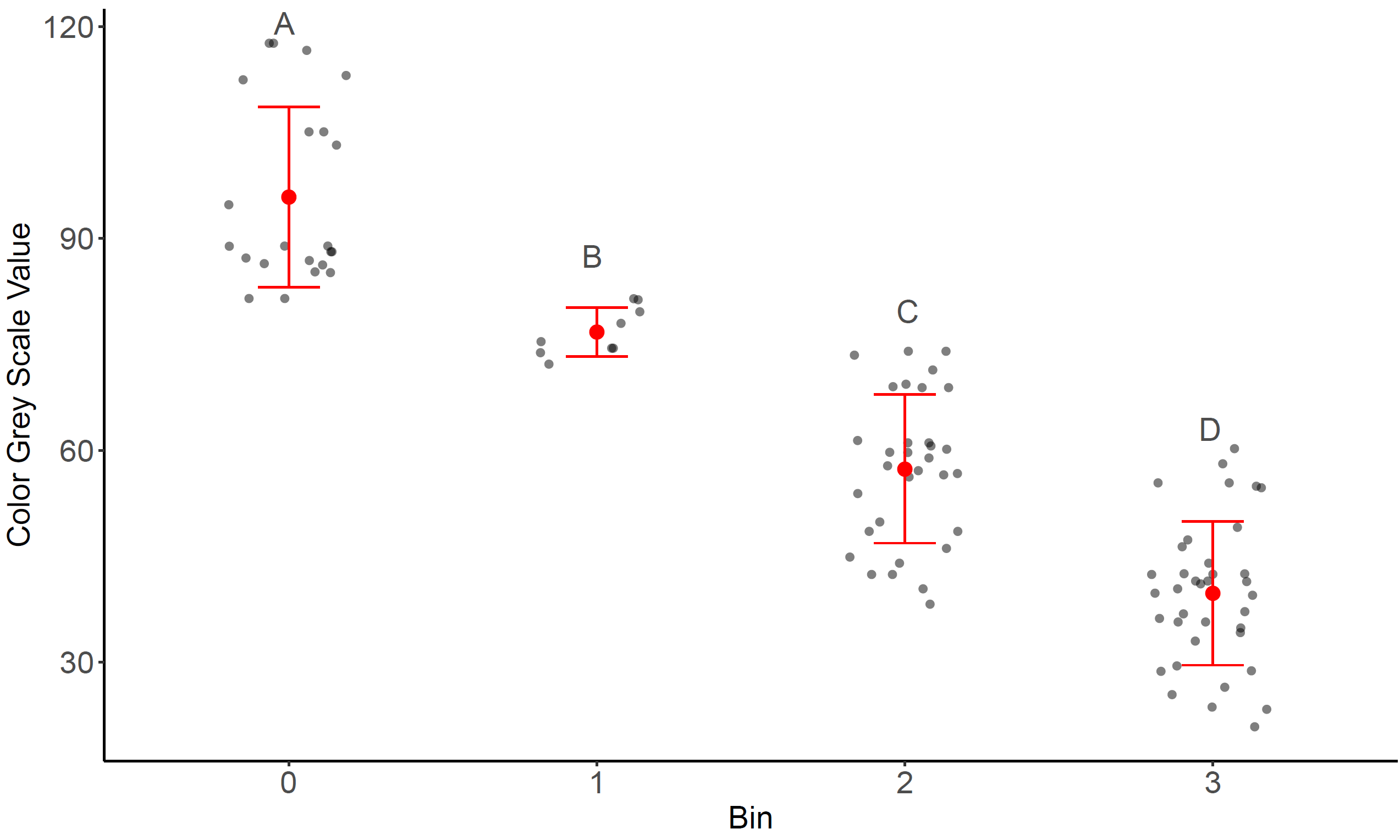

Figure S1: Plot of Binned Photos at Start of the Experiment. Letters indicate significance between bins.

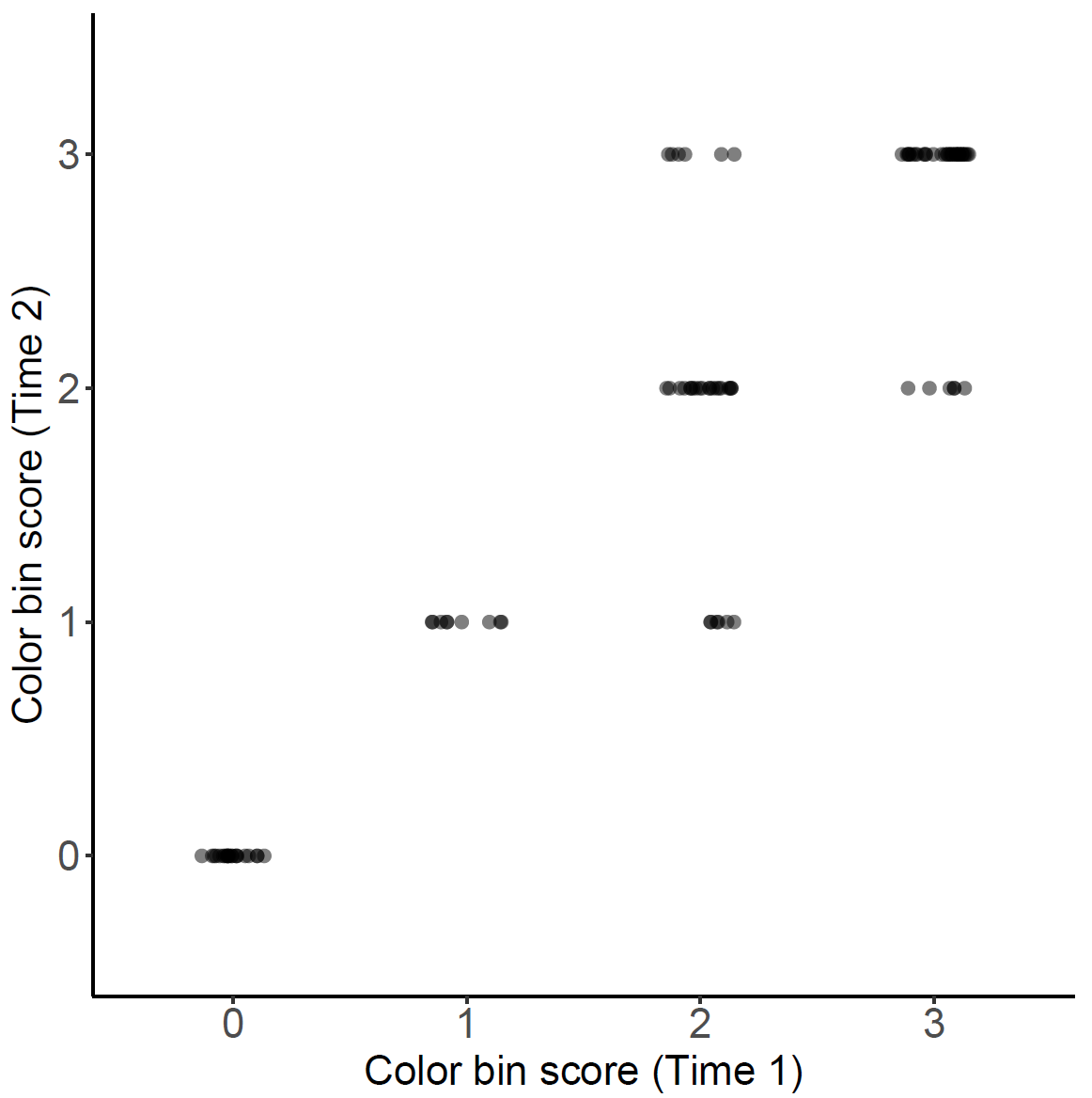

Figure S2: Repeatability of visual binning scores for dorsal thorax pigmentation. Each point represents a single fly scored twice by eye for trident pigmentation, using a categorical scale from 0 (no trident) to 3 (fully pigmented trident). The x-axis shows the score from the first scoring session (Time 1), and the y-axis shows the corresponding score from the second session (Time 2). Most points fall along the diagonal, indicating high consistency in visual binning across time.

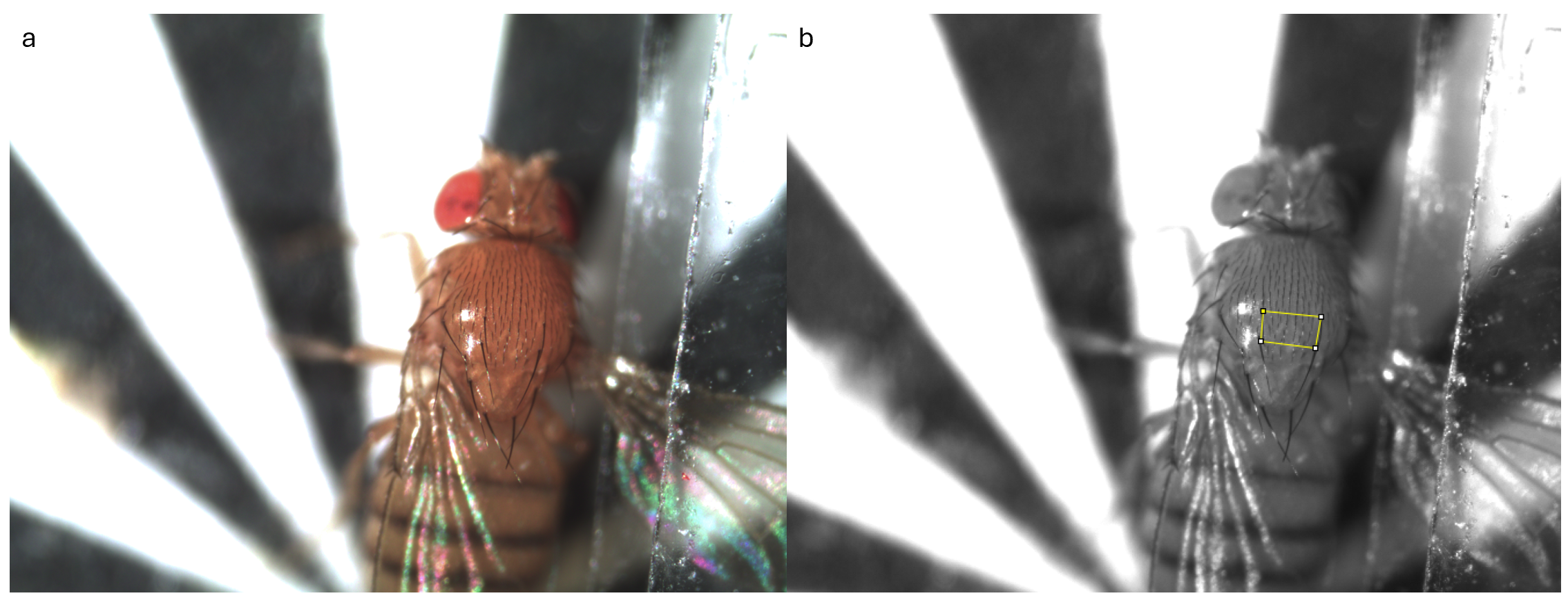

B

A

Figure S3: Example Photograph of Fly in WINGMACHINE. a) The original photo was taken using the WINGMACHINE. b) 8-bit converted photo with marks on bristle attachment sites to measure grey-scale value in ImageJ.

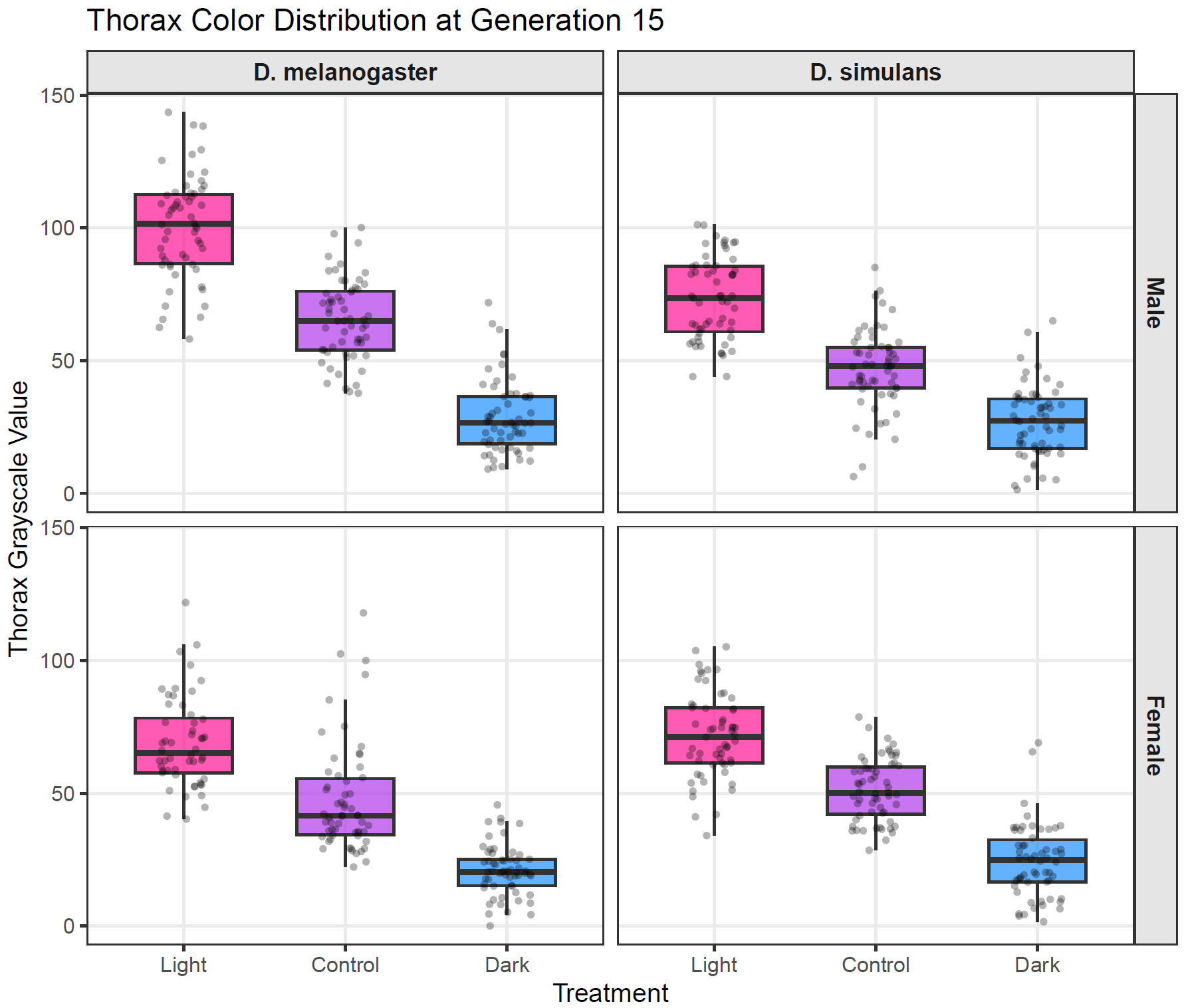

Figure S4: Thorax color distributions at generation 15. Boxplots show the distribution of thorax grayscale values for light (pink), control (purple), and dark (blue) selected individuals at generation 15. Raw data points are overlaid as jittered dots. Panels are separated by sex (rows) and species (columns). Lower grayscale values indicate darker pigmentation.

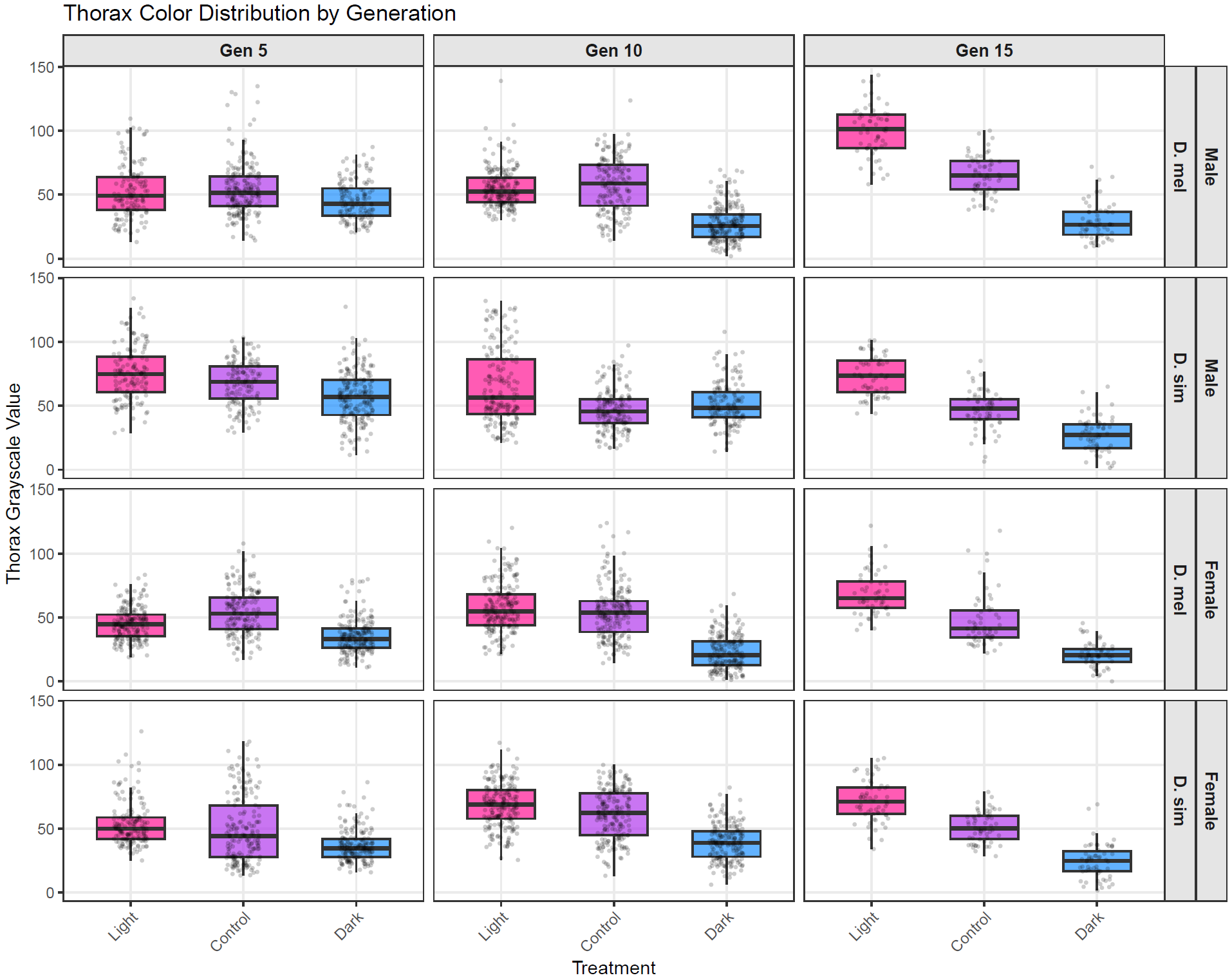

Figure S5: Thorax color distributions across selected generations. Boxplots show the distribution of thorax grayscale values for light (pink), control (purple), and dark (blue) selected individuals at generations 5, 10, and 15. Raw data points are overlaid as jittered dots. Panels are separated by sex and species. Lower grayscale values indicate darker pigmentation.

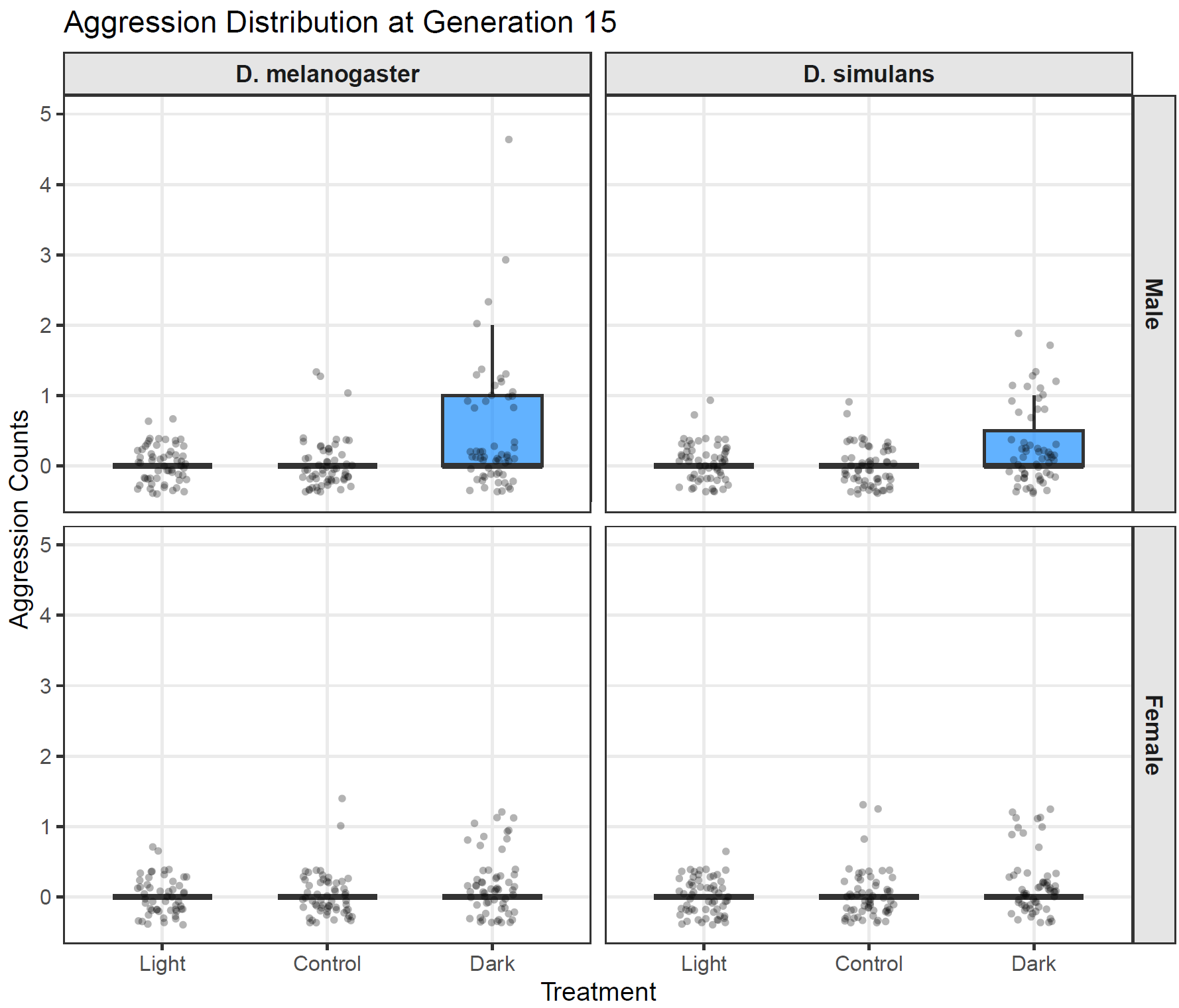

Figure S6: Aggression distributions at generation 15. Boxplots show the distribution of aggression counts for light (pink), control (purple), and dark (blue) selected individuals at generation 15. Raw data points are overlaid as jittered dots. Panels are separated by sex (rows) and species (columns).

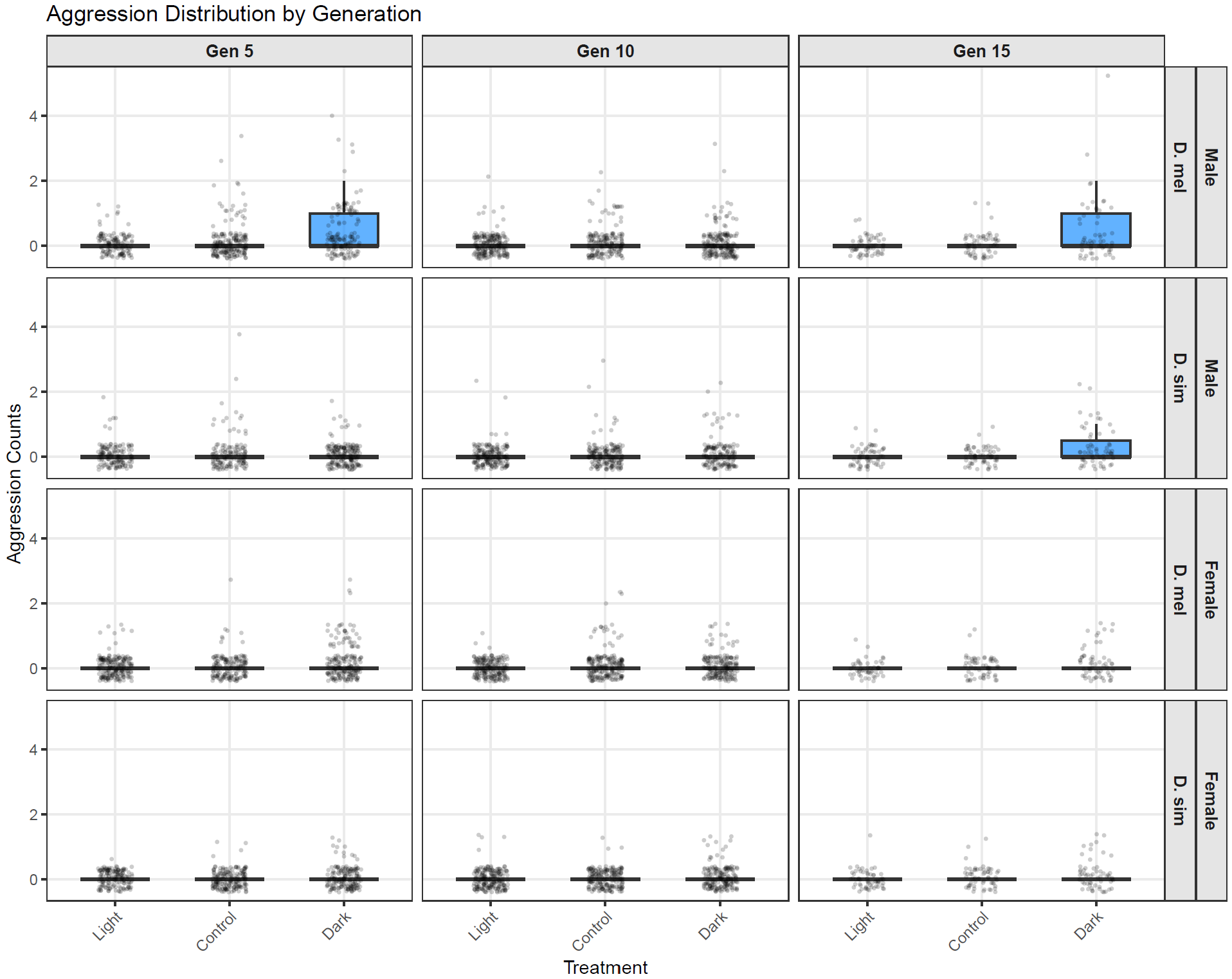

Figure S7: Aggression distributions across selected generations. Boxplots show the distribution of aggression counts for light (pink), control (purple), and dark (blue) selected individuals at generations 5, 10, and 15. Raw data points are overlaid as jittered dots. Panels are separated by sex and species.

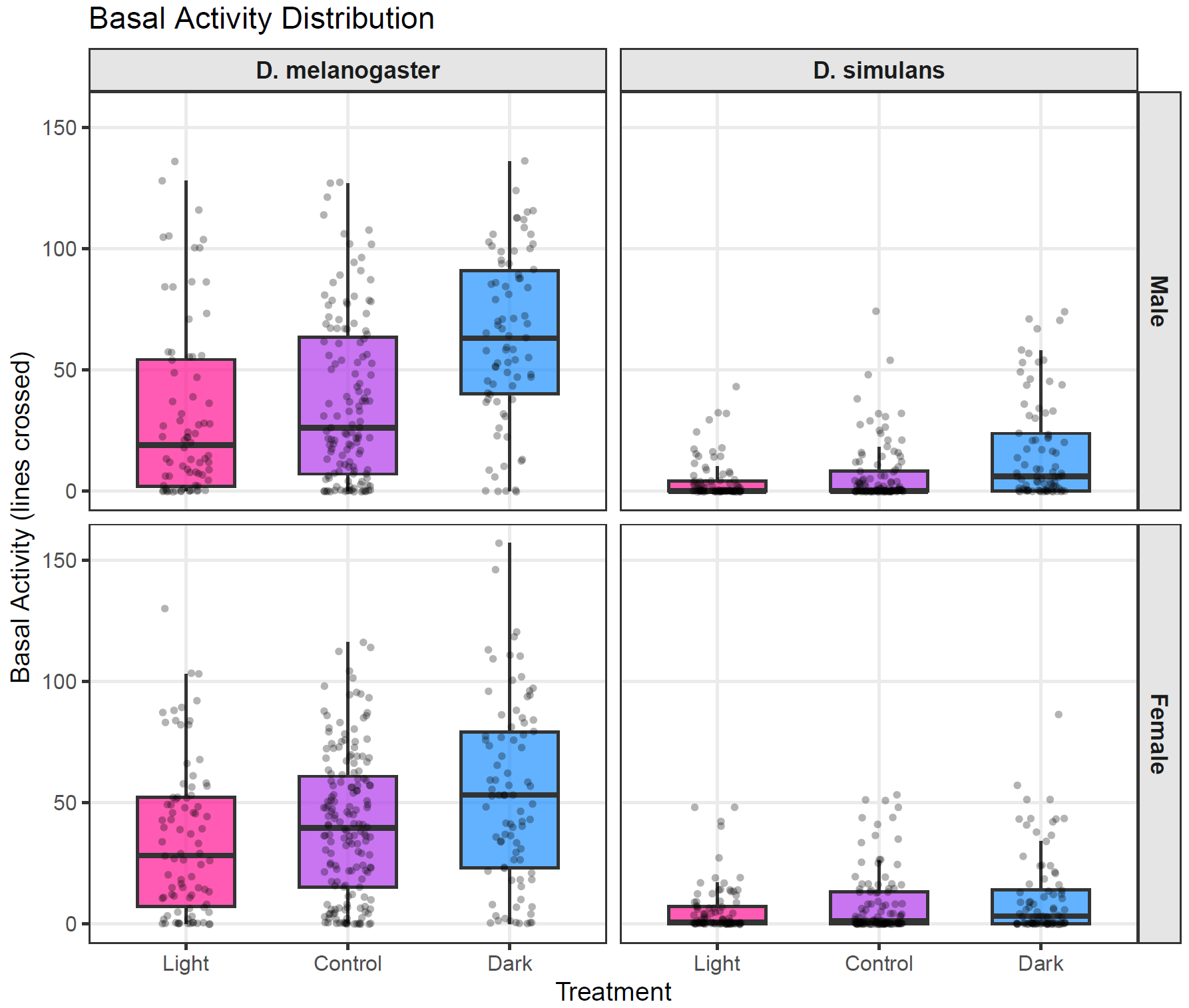

Figure S8: Basal activity distributions at generation 12. Boxplots show the distribution of basal activity (lines crossed in an open field arena) for light (pink), control (purple), and dark (blue) selected individuals at generation 12. Raw data points are overlaid as jittered dots. Panels are separated by sex (rows) and species (columns).

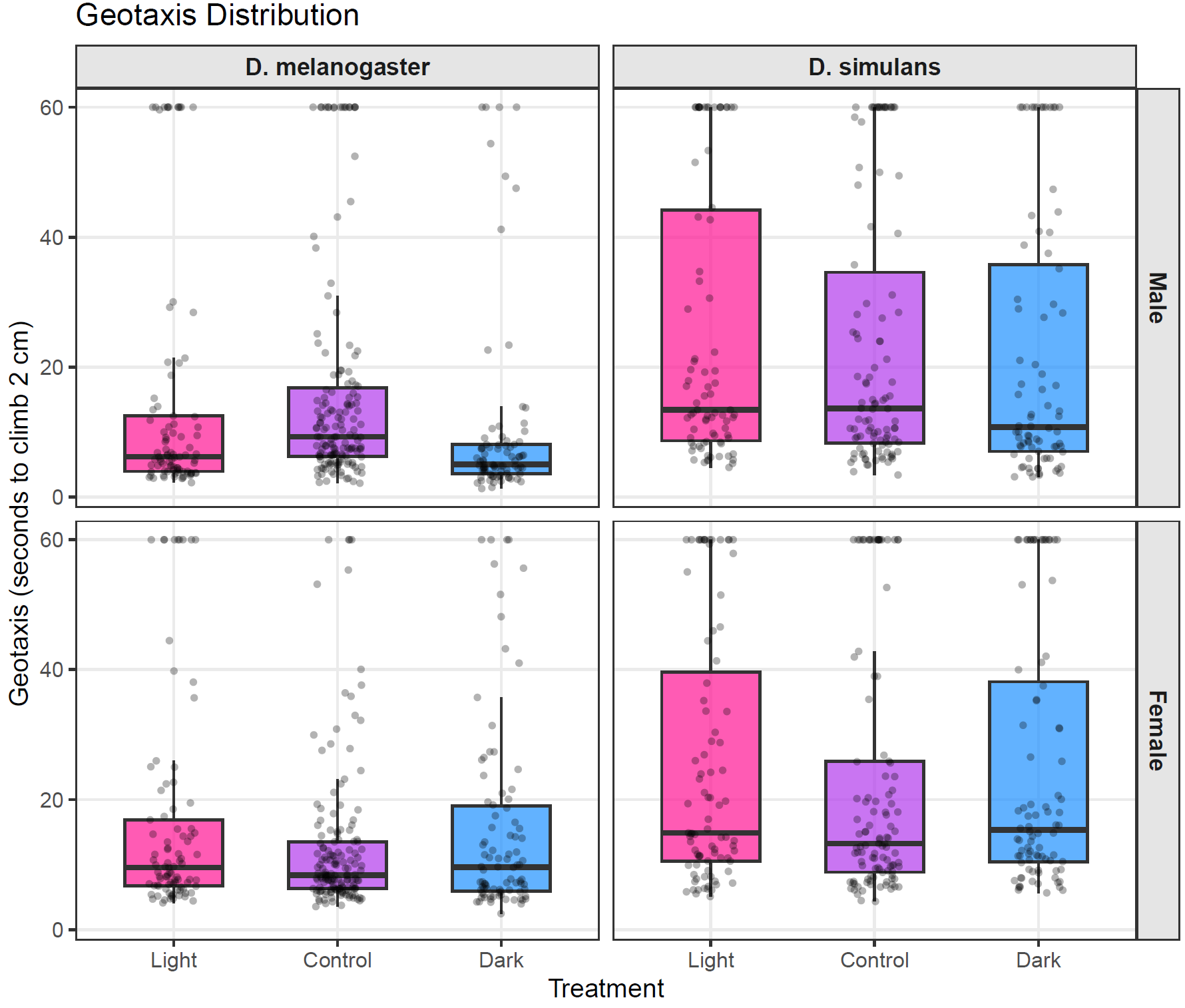

Figure S9: Geotaxis distributions at generation 13. Boxplots show the distribution of geotaxis latency (seconds to climb 2 cm) for light (pink), control (purple), and dark (blue) selected individuals at generation 13. Raw data points are overlaid as jittered dots. Panels are separated by sex (rows) and species (columns). Individuals that did not complete the task within 60 seconds are not shown.

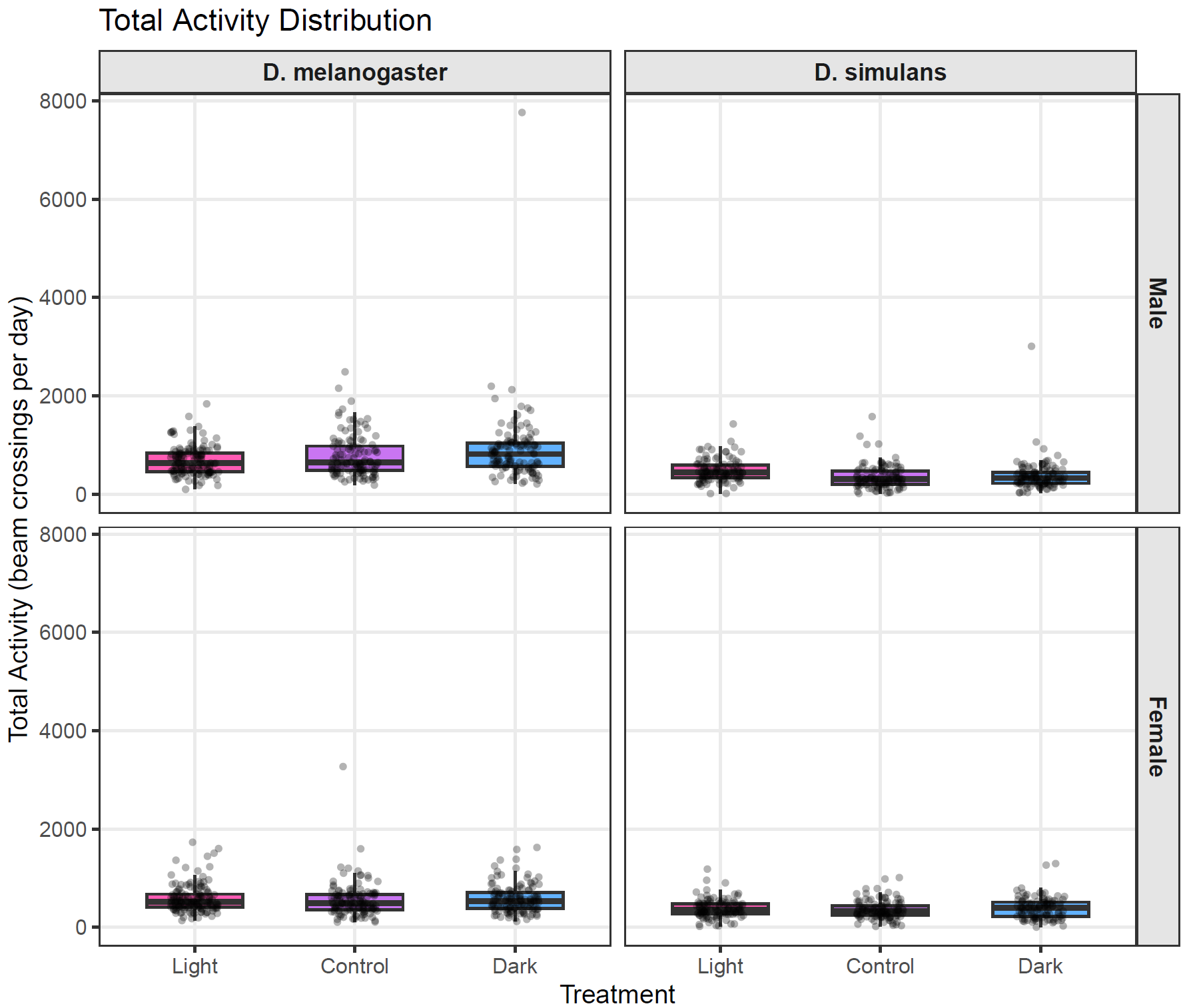

Figure S10: Total activity distributions at generation 16. Boxplots show the distribution of total daily activity (beam crossings per day) measured using the Drosophila Activity Monitor system for light (pink), control (purple), and dark (blue) selected individuals at generation 16. Raw data points are overlaid as jittered dots. Panels are separated by sex (rows) and species (columns).

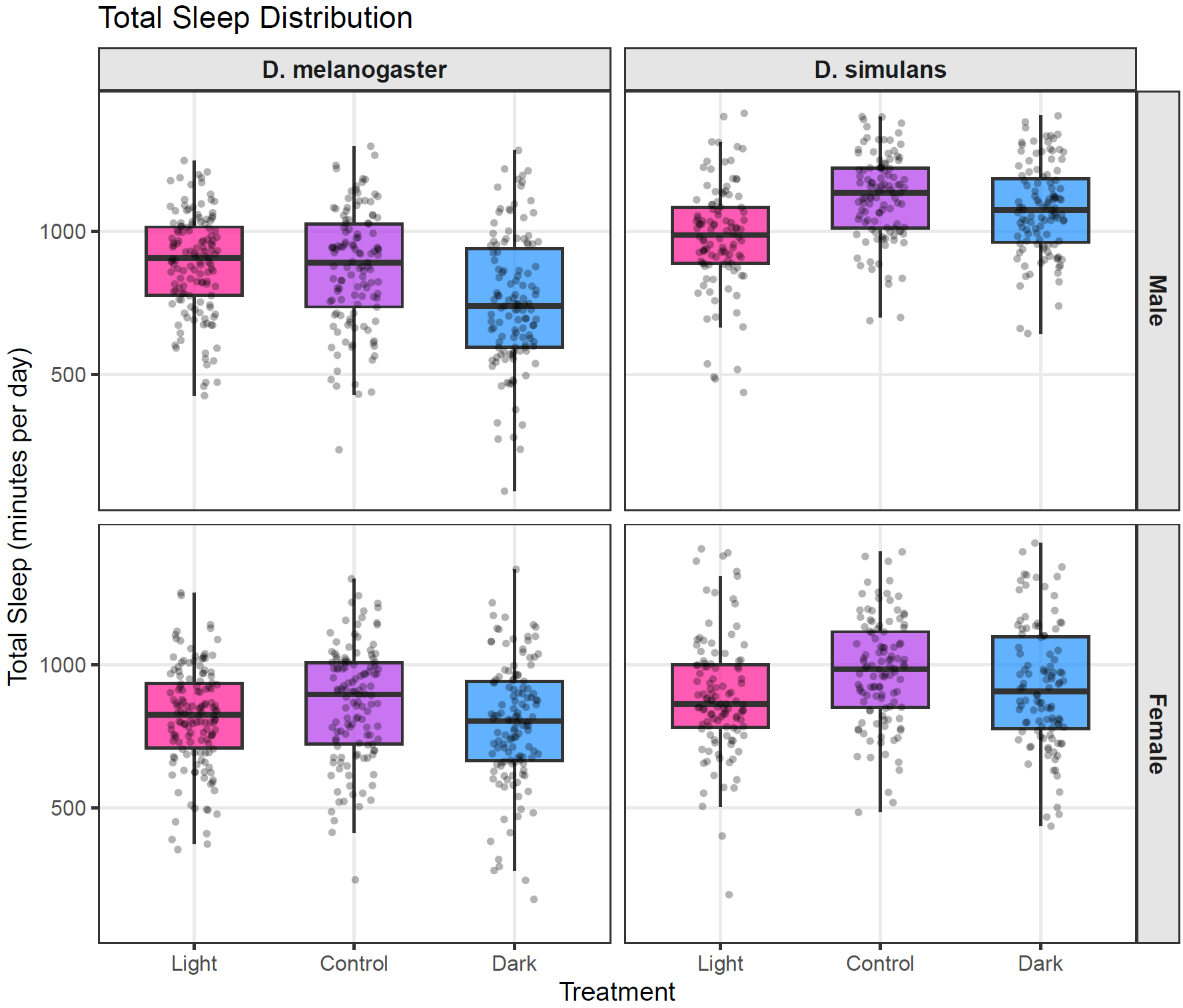

Figure S11: Sleep distributions at generation 16. Boxplots show the distribution of total daily sleep (minutes per day) measured using the Drosophila Activity Monitor system for light (pink), control (purple), and dark (blue) selected individuals at generation 16. Raw data points are overlaid as jittered dots. Panels are separated by sex (rows) and species (columns).

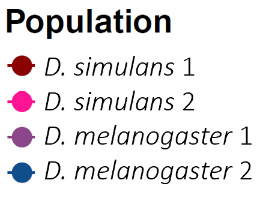

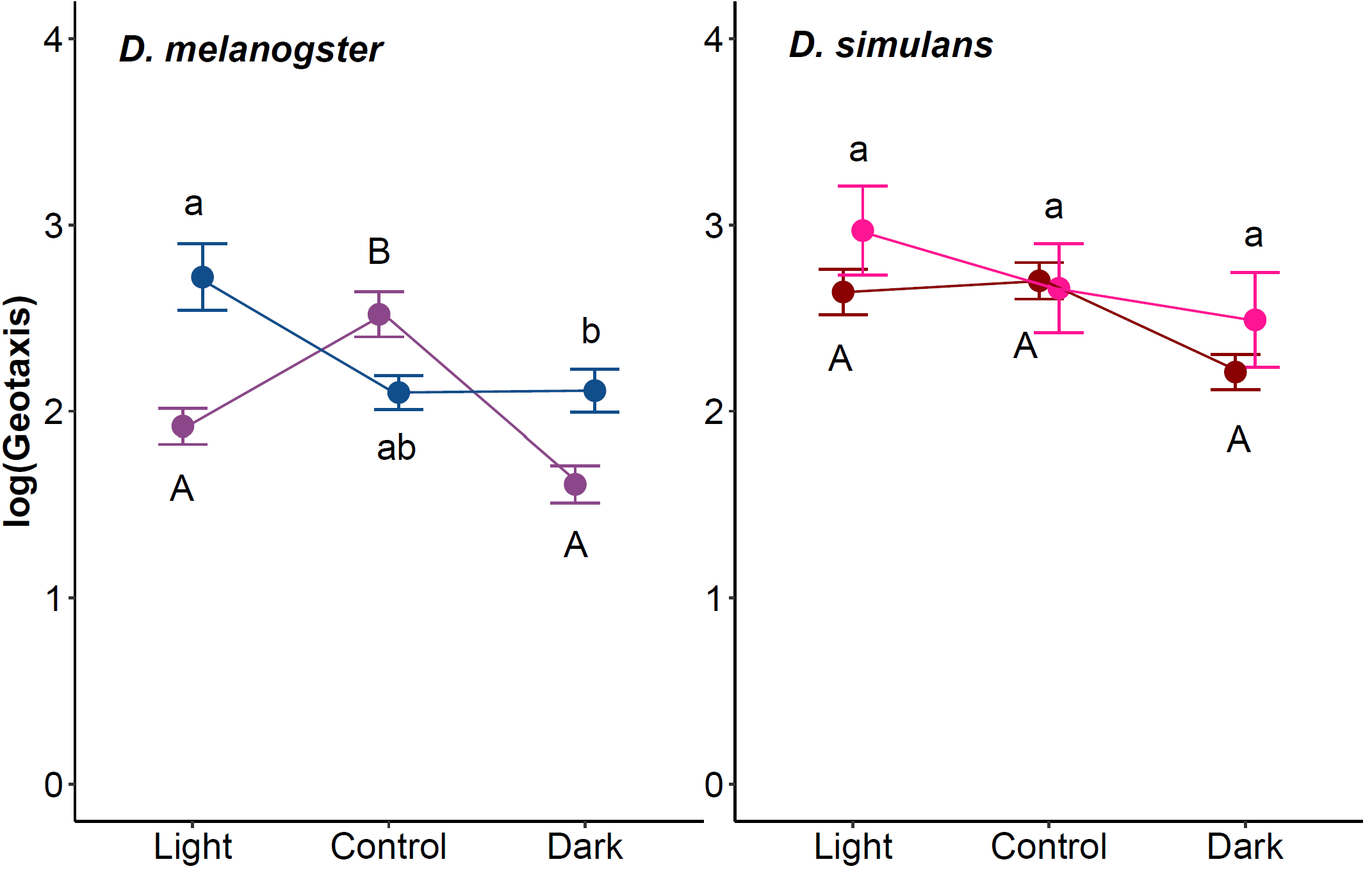

**A**

**B**

Figure S12: Male geotaxis at generation 13 for A) *D. melanogaster* and B) *D. simulans*. The x-axis indicates the different treatments (L, C, and D). The y-axis represents the log scale of the geotaxis times (time to cross second line). The letters above and below the data indicate significance, where groups with different letters are significantly different from each other. Uppercase letters indicate the significance of *D. mel*-1 and *D. sim*-1, and lowercase letters indicate the significance of *D. mel*-2 and *D. sim*-2 populations.

**A**

**B**

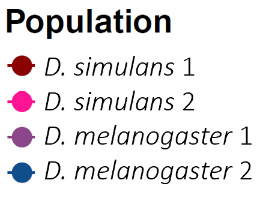

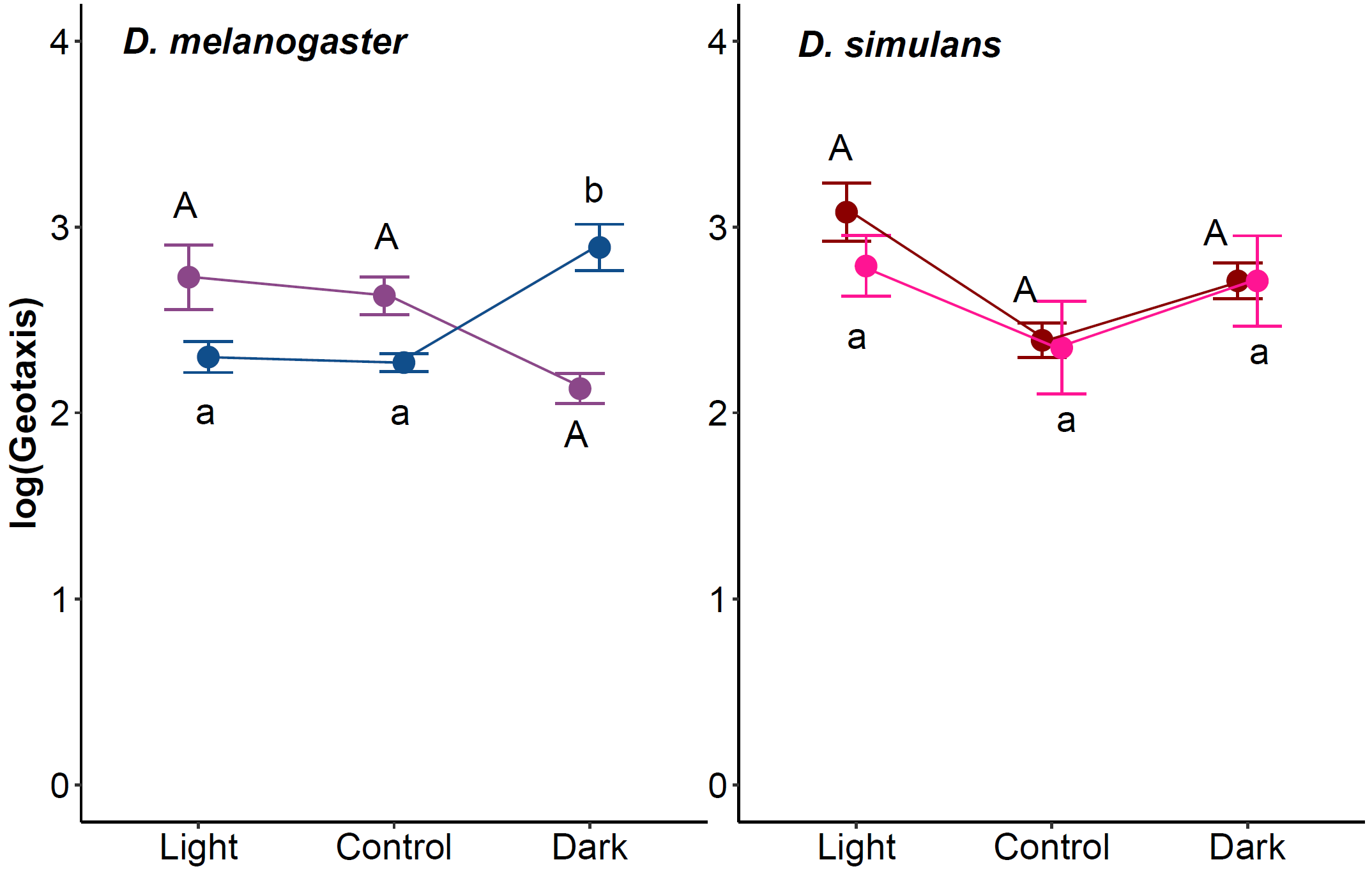

Figure S13: Female geotaxis at generation 13 for A) *D. melanogaster* and B) *D. simulans*. The x-axis indicates the different treatments (L, C, and D). The y-axis represents the log scale of the geotaxis times (time to cross second line). The letters above and below the data indicate significance, where groups with different letters are significantly different from each other. Uppercase letters indicate the significance of *D. mel*-1 and *D. sim*-1, and lowercase letters indicate the significance of *D. mel*-2 and *D. sim*-2 populations.

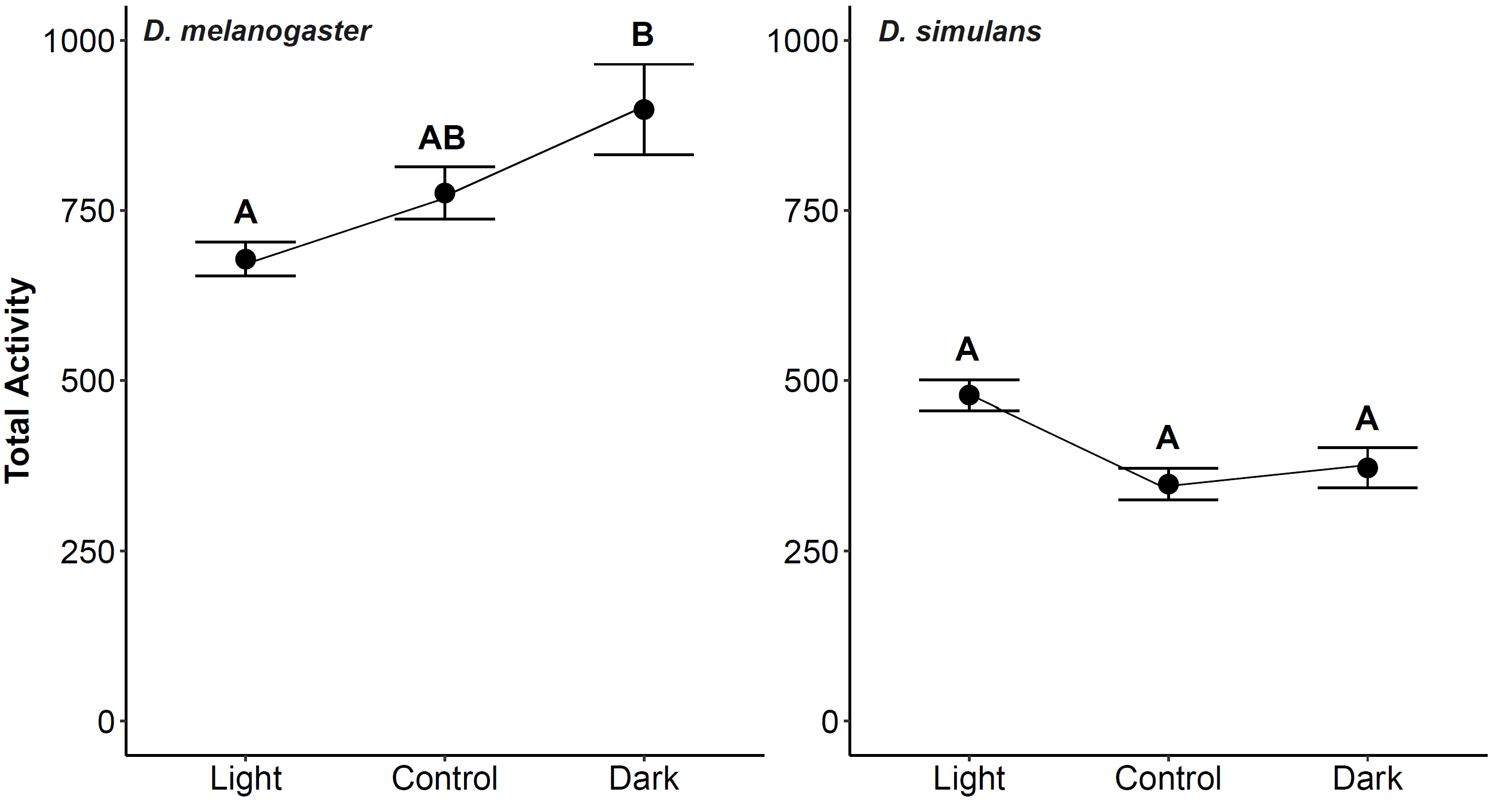

**B**

**A**

Figure S14: Male total activity level at generation 16 for A) *D. melanogaster* and B) *D. simulans*. The x-axis indicates the different treatments (L, C, and D). The y-axis represents the total activity level (average number of beams crossed over three days). The letters above the data indicate significance, where groups with different letters are significantly different from each other.

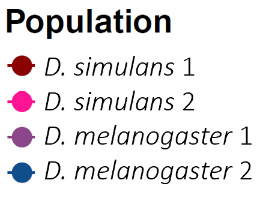

**B**

**A**

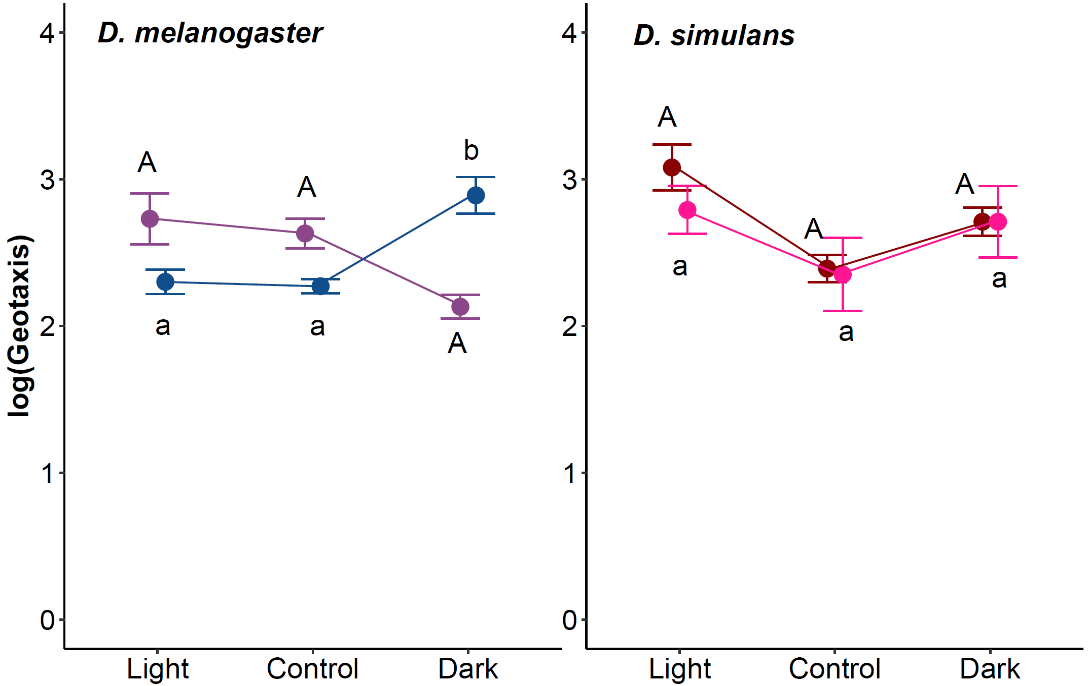

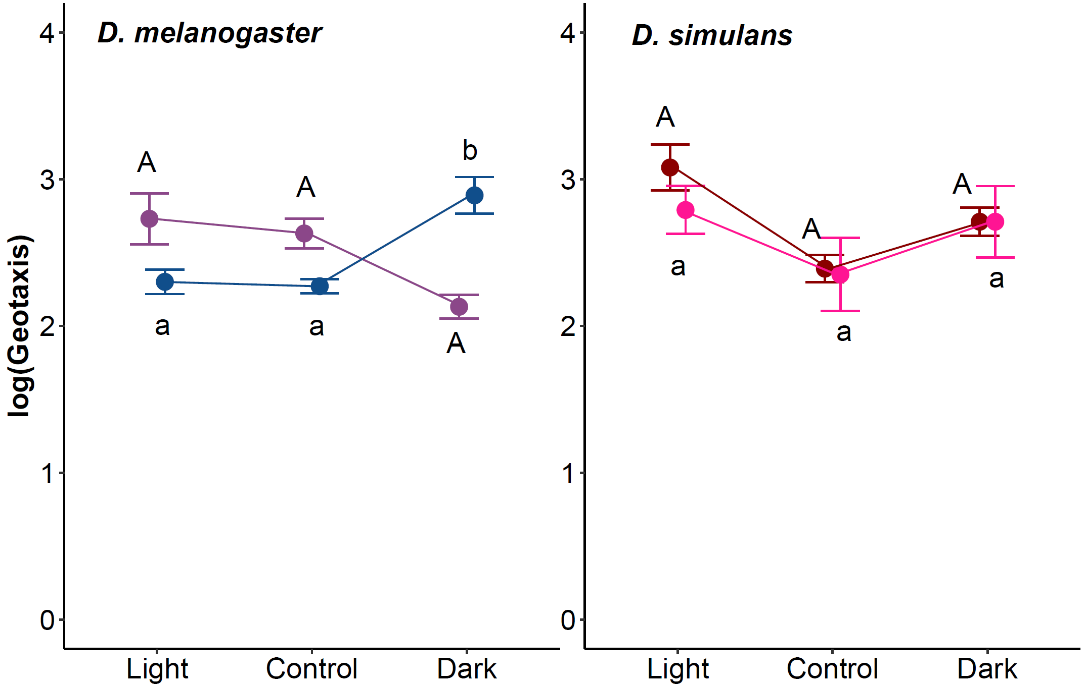

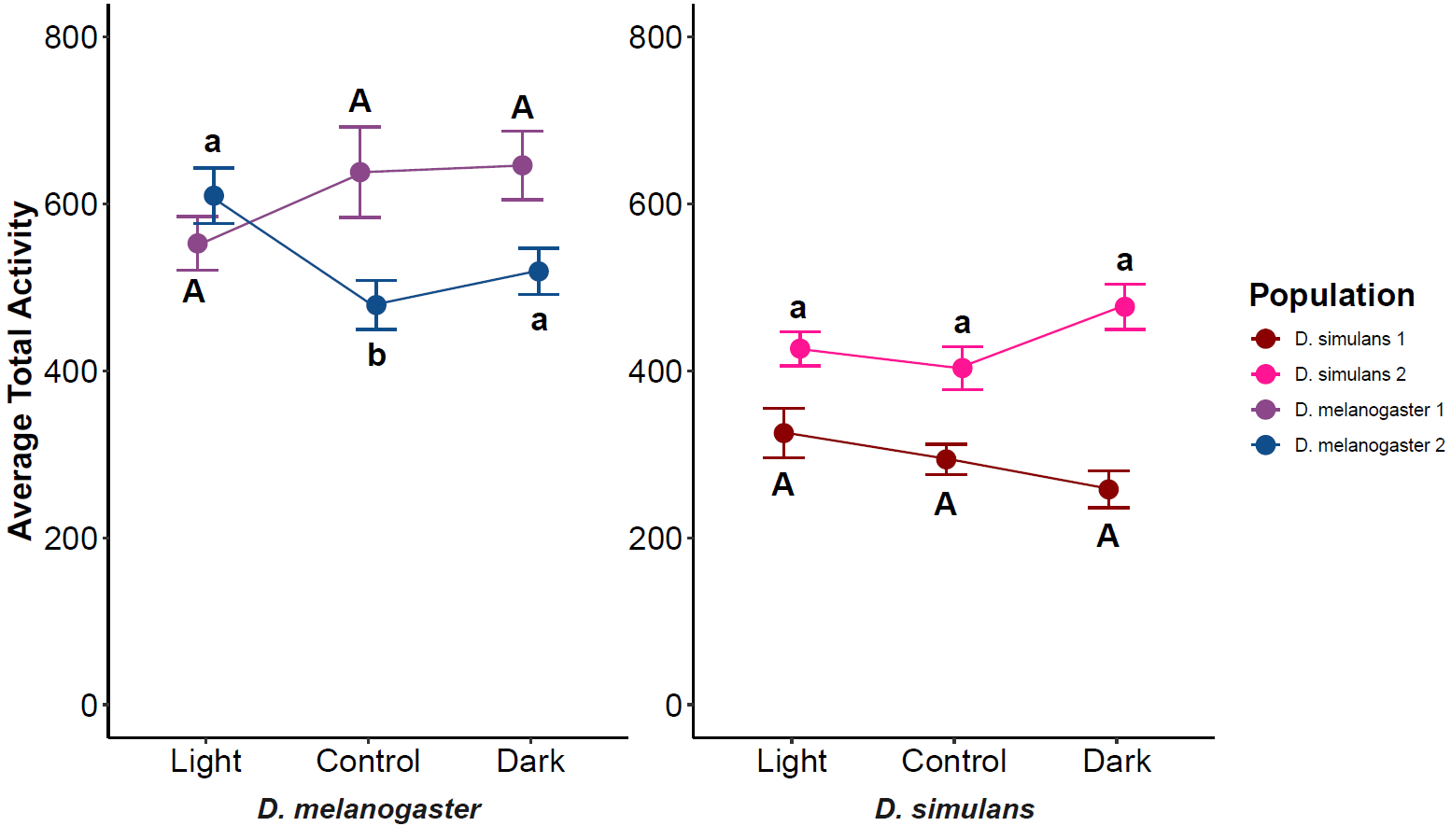

Figure S15: Female total activity level at generation 16 for A) *D. melanogaster* and B) *D. simulans*. The x-axis indicates the different treatments (L, C, and D). The y-axis represents the total activity level (average number of beams crossed over three days). The letters above and below the data indicate significance, where groups with different letters are significantly different from each other. Uppercase letters indicate the significance of *D. mel*-1 and *D. sim*-1, and lowercase letters indicate the significance of *D. mel*-2 and *D. sim*-2 populations.

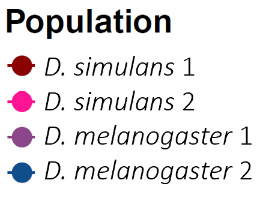

**A**

**B**

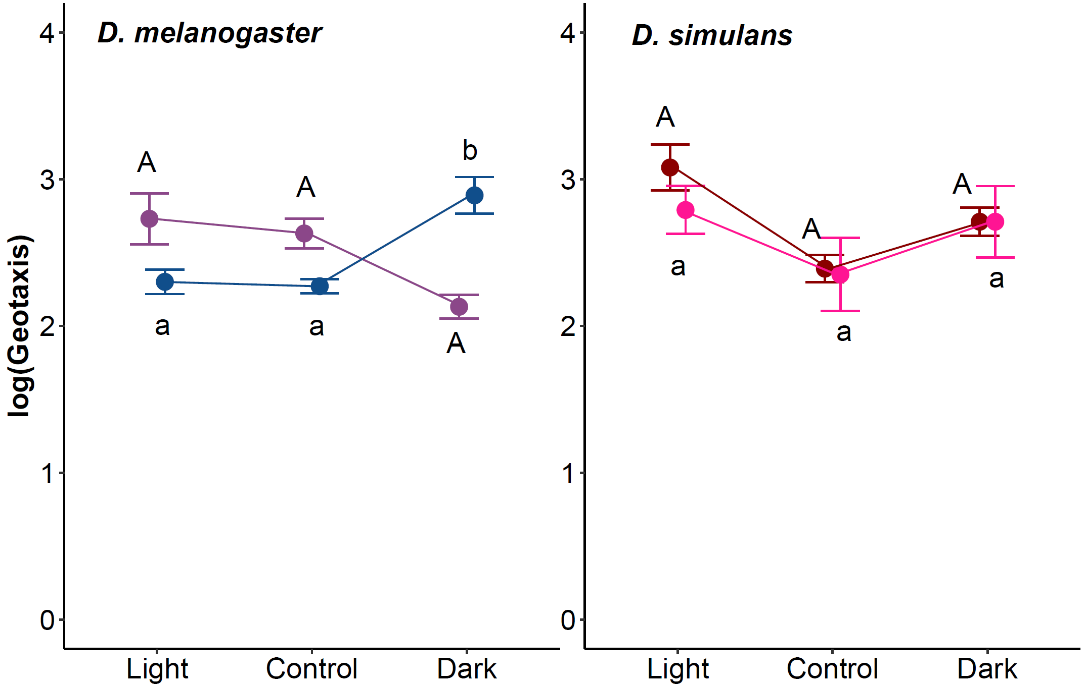

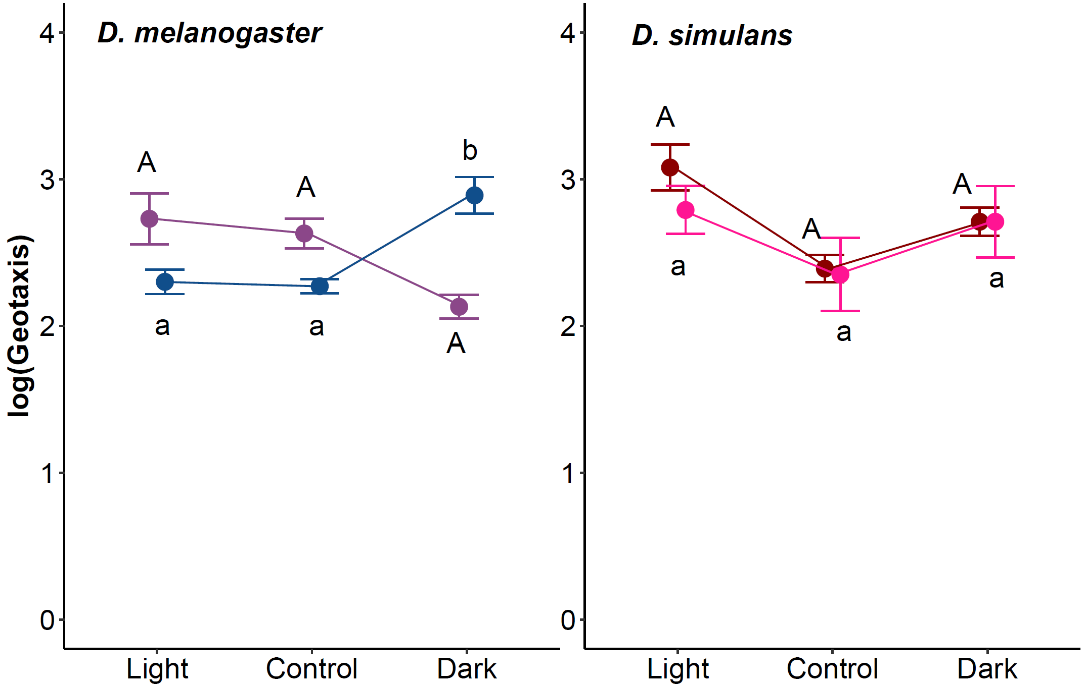

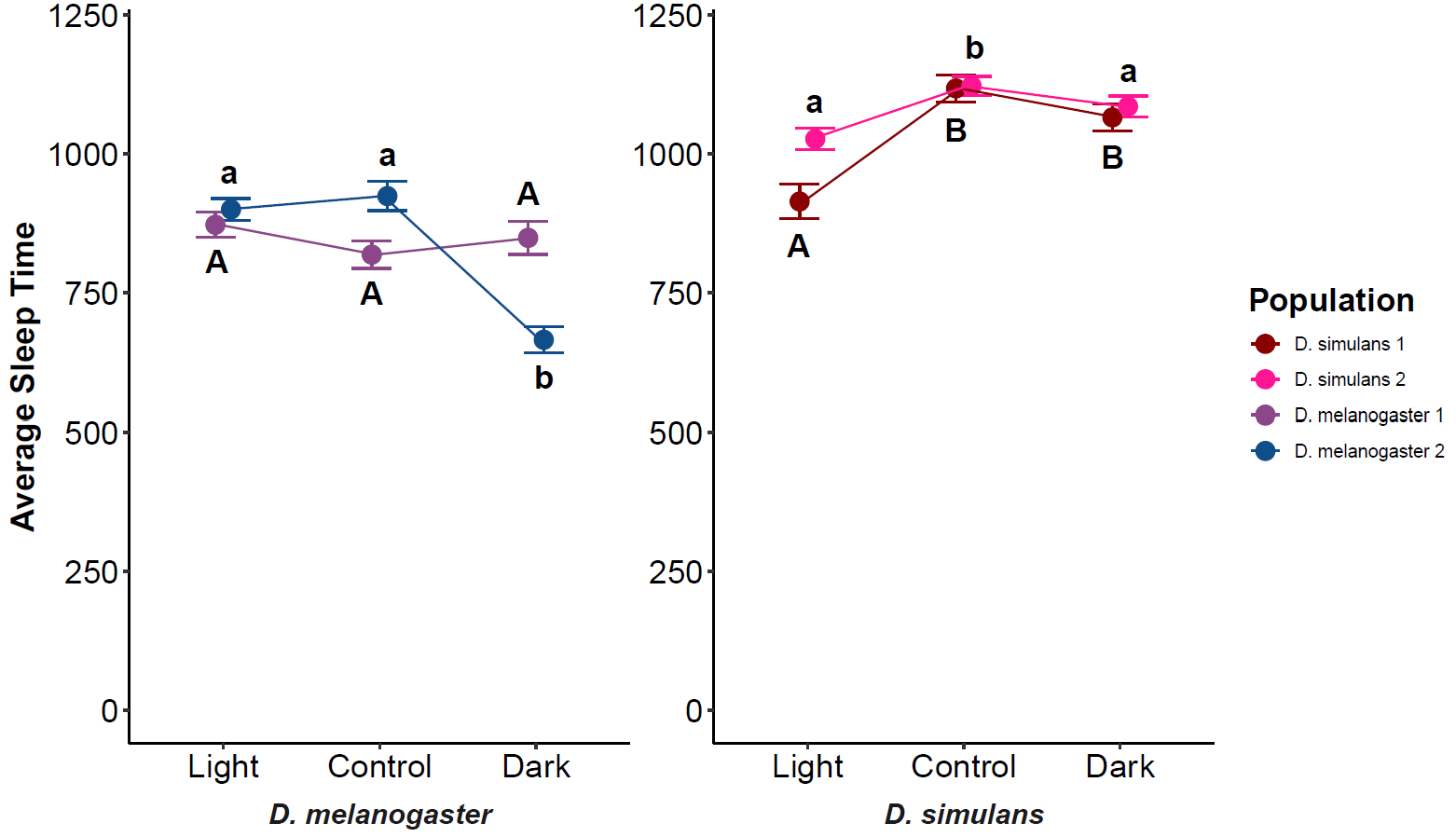

Figure S16: Male total sleep at generation 16 for A) *D. melanogaster* and B) *D. simulans*. The x-axis indicates the different treatments (L, C, and D). The y-axis represents the average sleep time (average time spent sleeping over three days). The letters above and below the data indicate significance, where groups with different letters are significantly different from each other. Uppercase letters indicate the significance of *D. mel*-1 and *D. sim*-1, and lowercase letters indicate the significance of *D. mel*-2 and *D. sim*-2 populations.

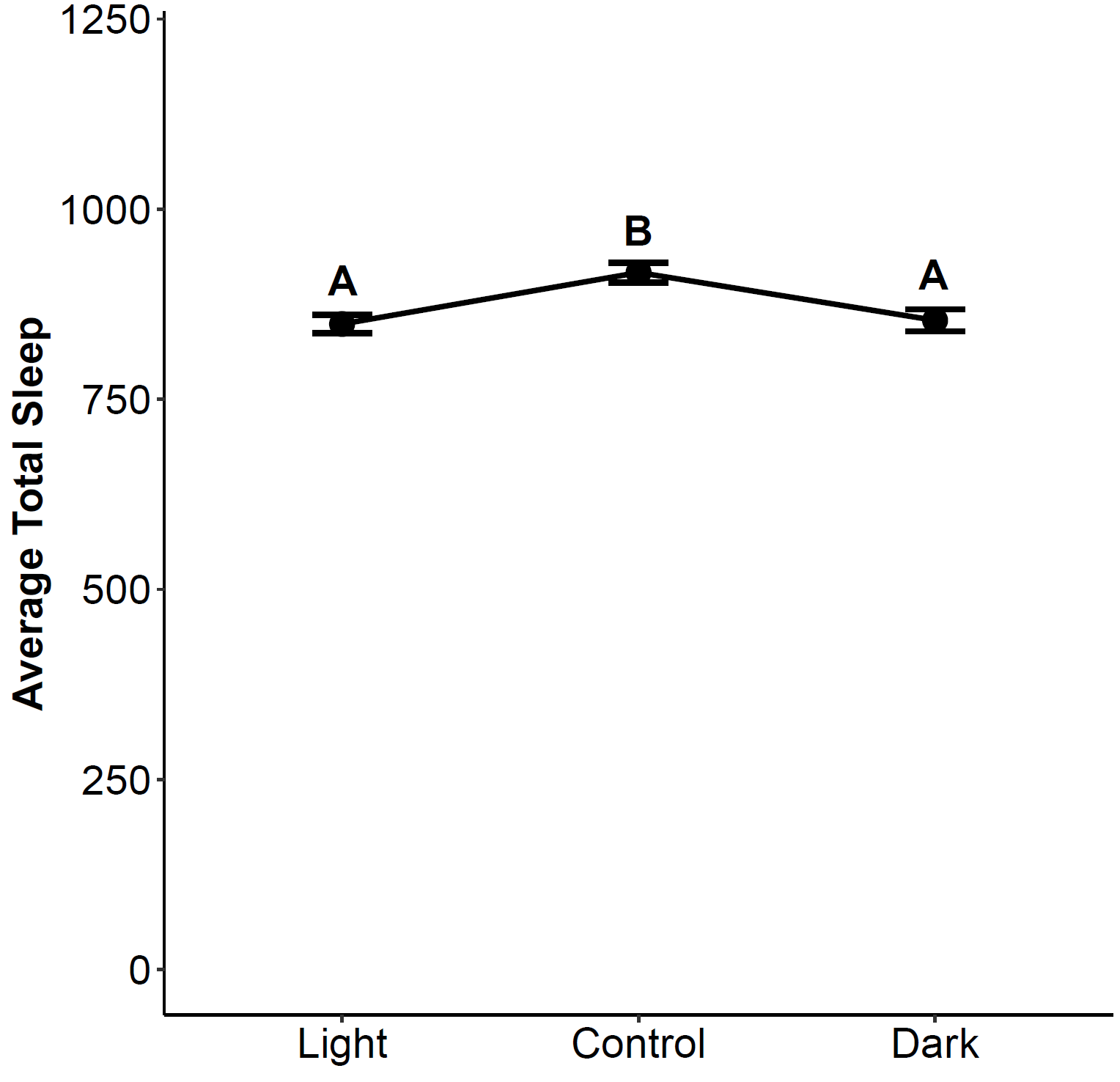

Figure S17: Female total sleep at generation 16. The x-axis indicates the different treatments (L, C, and D). The y-axis represents the average sleep time (average time spent sleeping over three days). The letters above the data indicate significance, where groups with different letters are significantly different from each other.
